# Intercellular Signaling Pathways as Therapeutic Targets for Vascular Dementia Repair

**DOI:** 10.1101/2024.03.24.585301

**Authors:** Min Tian, Riki Kawaguchi, Yang Shen, Michal Machnicki, Nikole G. Villegas, Delaney R. Cooper, Natalia Montgomery, Jacqueline Haring, Ruirui Lan, Angelina H. Yuan, Christopher K. Williams, Shino Magaki, Harry V. Vinters, Ye Zhang, Lindsay M. De Biase, Alcino J. Silva, S. Thomas Carmichael

## Abstract

Vascular dementia (VaD) is a white matter ischemic disease and the second-leading cause of dementia, with no direct therapy. Within the lesion site, cell-cell interactions dictate the trajectory towards disease progression or repair. To elucidate the underlying intercellular signaling pathways, a VaD mouse model was developed for transcriptomic and functional studies. The mouse VaD transcriptome was integrated with a human VaD snRNA-Seq dataset. A custom-made database encompassing 4053 human and 2032 mouse ligand-receptor (L-R) interactions identified significantly altered pathways shared between human and mouse VaD. Two intercellular L-R systems, Serpine2-Lrp1 and CD39-A3AR, were selected for mechanistic study as both the ligand and receptor were dysregulated in VaD. Decreased Seprine2 expression enhances OPC differentiation in VaD repair. A clinically relevant drug that reverses the loss of CD39-A3AR function promotes tissue and behavioral recovery in the VaD model. This study presents novel intercellular signaling targets and may open new avenues for VaD therapies.

## Introduction

Vascular dementia (VaD) constitutes approximately 25% of total dementia. It frequently coexists with Alzheimer’s disease (AD) in an additive or synergistic manner; 84% of aged subjects show morphological substrates of VaD in addition to AD pathology [1]. Despite its high prevalence, the precise underlying mechanisms of VaD remain poorly understood, which can be attributed to the lack of suitable preclinical animal models. VaD is characterized by multiple infarcts or ischemia in the periventricular and adjacent white matter (WM), leading to progressive deficits in memory and motor functions. However, the current animal models for VaD predominantly rely on globally induced cerebral hypoperfusion through vessel occlusion in rodents, genetically hypertensive brain and cerebrovascular disease, bilateral/asymmetrical carotid artery stenosis (BCAS/ACAS) or bilateral occlusion of the common carotid arteries (BCAO) in rat or mouse brains [2, 3]. These models suffer from several significant limitations, including in some cases high lethality, substantial variability in lesion size, and the presence of widespread neuronal death in gray matter, which is not observed in human VaD. There is an urgent and significant need for a replicable, robust, and more specific WM-focused mouse VaD model that encompasses the diverse etiologies observed in VaD patients.

The cells of the cerebral WM exist in a neurovascular niche, which supports cell homeostasis and injury response through cell-cell signaling. The neurovascular niche has membrane, soluble and extracellular matrix signals that communicate among its constituent endothelial cells (ECs), pericytes, astrocytes and oligodendrocyte progenitor cells (OPCs) [4, 5] in many brain diseases to dictate survival, recovery or further disease progression [6]. The full cell-cell signaling systems in VaD are not known. This VaD cell-to-cell “interactome” may provide starting points for candidate therapeutic systems in this disease.

The current study developed a VaD model using the common C57BL/6J (C57) mouse strain from a WM stroke model using immunocompromised mice [7]. This mouse model reproduces the cellular, circuit, and behavioral impairments in human VaD. Cell type-specific RNA-Seq reveals activation of WM-associated aging genes in VaD mouse model, and identifies a unique transcriptome of WM as compared to the same cell types in gray matter (cortex). Intercellular signaling pathways dysregulated in the VaD neurovascular niche were identified from a custom ligand-receptor database. The potential candidates in the same cell types were compared between mouse and human VaD [8] to identify key signaling systems for functional studies. An important extracellular matrix component, Serpine2, and its receptor Lrp1 were identified as elevated during VaD. Knocking down Serpine2 enhanced OPC differentiation towards myelinated oligodendrocytes in VaD. Another key gene system, identified as EC/microglia-microglia signaling through CD39-A3AR, was synergistically downregulated in the conjunction of VaD and aging in human and mouse. An A3AR specific agonist, currently in phase III trials for psoriasis, promoted tissue repair and behavioral recovery in the mouse VaD model in a delayed treatment. These findings shed light on the intercellular signaling pathways that could serve as therapeutic targets for VaD.

## Results

### A mouse model that recapitulates multiple features of human vascular dementia (VaD)

Confluent infarcts were induced in the cerebral white matter (WM) of C57 mouse brains by intracranial injections of the vasoconstrictor L-NIO (dihydrochloride) (Fig. 1a). The resultant lesion was in several hundred micrometers along the medial-lateral and anterior-posterior axes in corpus callosum (CC) and cingulum (Fig. 1a). This lesion covered the most common sites of WM damage in human VaD [9]. It produced significant axon and myelin loss, astrocyte (Fig. 1b, Supplementary Fig. 1a) and microglial activation, OPC proliferation (Fig. 1c), and pericyte responses (Supplementary Fig. 1b).

**Figure 1.**
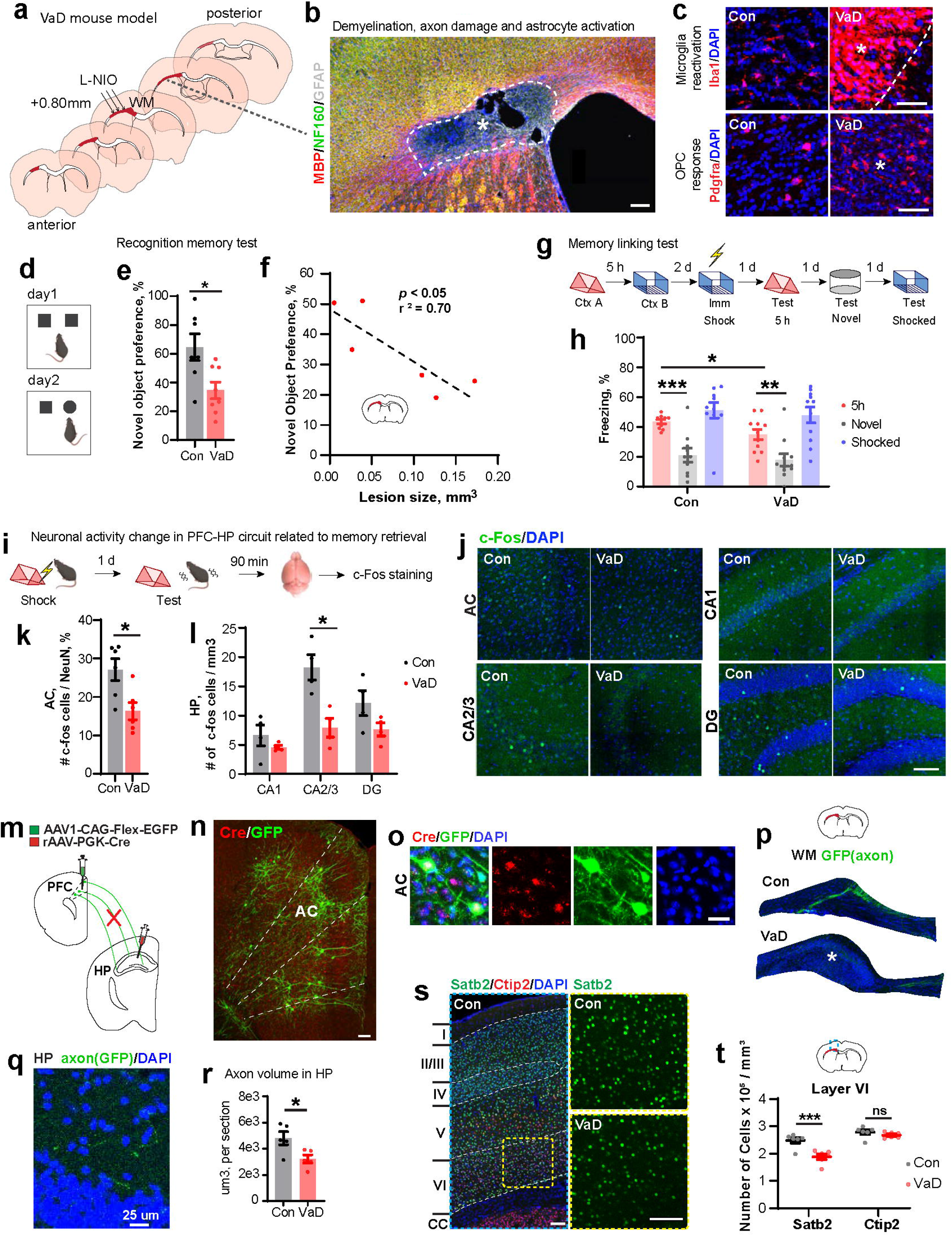
A mouse model that mimics vascular dementia (VaD) in human. (a) Schematic shows the induction of VaD in mouse cerebral white matter by 3 intracranial injections of vasoconstrictor L-NIO; lesion location highlighted in maroon. (b) A representative immunostaining image of infarct core in VaD mouse model shows the demyelination (degradation of MBP, red), axon damage (breakdown of NF160, green), and astrocyte activation (white). Scale bar = 100μm. DAPI, blue. (c) Representative immunostaining images showing the activation of microglia (Iba1, red), and OPC (Pdgfrα, red) response in VaD lesion (star sign shows the center, dashed line shows the border). Scale bars = 50μm. DAPI, blue. (d) Schematic of the Novel object recognition test. (d) Quantification of novel object preference (%). Con vs. VaD, * *p* < 0.05, t test. (f) Regression of novel object preference (%) against lesion size (mm^3^). Simple linear regression, *p* < 0.05, r^2^ = 0.70. (g) Schematic shows the paradigm of memory linking test. (h) Quantification of freezing (%) in associated (A, red), novel (gray), and shocked (B, blue) contexts. Con (A) vs VaD (A), * *p* < 0.05; Con (A) vs VaD (novel), *** *p* < 0.001; VaD (A) vs VaD (novel), ** *p* < 0.01. Each dot represents an individual animal. Two-way repeated measures ANOVA. (i) Schematic shows the test of neuronal activity in prefrontal cortex-hippocampus (PFC-HPC) circuits related to memory retrieval using Fear conditioning test and immediate early gene (c-Fos) staining. (j) Representative images show the difference of c-Fos (green) signal in AC, hippocampus (including CA1, CA2/3, DG subregions) in control and VaD brains. DAPI, blue. (k) Quantification of c-Fos cells in AC. Con vs. VaD, * *p* < 0.05, t test. (l) Quantification of c-Fos cells in HPC. CA2/3, Con vs. VaD, * *p* < 0.05, t test. (m) Schematic shows the retrograde labeling of HPC projecting neurons from PFC by injections of retroAAV-PGK-cre virus in HPC, and AAV1-CAG-Flex-EGFP in PFC. (n) A representative immunostaining image shows the labeling of GFP (green), and Cre recombinase (red) in prefrontal cortex including anterior cingulate (AC) subregion. Scale bar = 50μm. Dashed lines show the border of PFC subregions. (o) Representative images show the colocalization of Cre expression and retrogradely labeled neurons by GFP (greed). DAPI, blue. Scale bar = 25μm. (p) Schematic and images show the region of infarct core in VaD model, and axon bundles (labeled with GFP, green) passing the regions in control and VaD brains. Star sign indicates the core of infarct lesion. (q) A representative image shows the axon processes (labeled with GFP, green) in the hippocampus. Scale bar = 25μm. DAPI, blue. (r) Quantification result of axon volume in HPC labeled with GFP. Con vs. VaD, * *p* < 0.05, t test. (s) Representative images show the immunostaining of Satb2 (green), Ctip2 (red) in Layer I-VI above infarct lesion of VaD mouse model, and the comparison of Satb2+ cell density in Layer VI in control and VaD brains. Scale bars = 100μm. DAPI, blue. (t) Schematic shows the region of brains (blue dashed box) for quantification of Satb2+ and Ctip2+ cells. Quantification result of Satb2+ and Ctip2+ numbers in Layer VI above infarct lesion. Satb2, Con vs VaD, *** *p* < 0.001; Ctip2, Con vs VaD, ns = not significant.

To test the cognitive deficits in this model, a two-day Novel Object Recognition (NOR) test (Fig. 1d) was employed [10]. The memory impairment in VaD was significant (Fig. 1e), and the extent of deficit was inversely correlated to the WM lesion size (Fig. 1f). To explore memory deficits observed in VaD patients at early stages [11, 12], the mice were tested in a contextual fear conditioning memory linking protocol [13] (Fig. 1g), an hippocampal-dependent test that evaluates the linking or integration of context memories across a 5-hour interval. A significant impairment in this task in the VaD cohorts was observed (Fig. 1h).

VaD is fundamentally a disease of disconnection, damaging axonal tracts in the subcortical WM. In human VaD, periventricular WM hyperintensities may cause anatomic damage to the cingulum and CC [9, 14-16], which connect the prefrontal cortex (PFC) to many brain regions including hippocampus (HPC) (a crucial region for learning and memory [17, 18]). To test this circuit in the mouse VaD model, we measured its activation and anatomical connectivity. Neuronal activation in PFC and HPC as measured by c-Fos (immediate early gene) levels quantified after memory retrieval (contextual fear conditioning) (Fig. 1j-l), which indicated a significant neuronal activity decrease in anterior cingulate (AC) in PFC (Fig. 1k), and in CA2/3 in HPC (Fig. 1l) due to PFC-HPC circuit damage in VaD. Anatomical connectivity was labeled by injections of AAV1-CAG-Flex-EGFP and retroAAV-PKG-Cre into the dorsal-medial PFC and HPC respectively (Fig. 1m). CA1, CA2/3 and DG subregions in HPC were labeled with Cre recombinase (supplementary Fig. 1c). PFC neurons were double-labeled with Cre and GFP (green fluorescence protein) (Fig. 1n, 1o) indicating a successful retrograde tracing. PFC axonal processes extended through subcortical WM (Fig. 1p, Con) and then projected toward the HPC (Fig. 1q). The ischemic lesion reduced the axon projection volume in WM (Fig. 1p, VaD, Supplementary Fig. 1d) and HPC (Fig. 1r), indicating the interruption of neural circuits that may subserve a memory function.

In multiple sclerosis, demyelinating lesions produce selective loss of neurons in overlying cortex expressing Cux2, a transcriptional factor (TF) and a cortical layer identity marker [19]. Assessment of TFs and cortical layer markers in VaD revealed that Satb2 (Fig. 1s, 1t) and Cux1 (Supplementary Fig. 1e), in lesion adjacent cortical layer VI, were significantly reduced (Fig. 1t), but not the layer VI marker Ctip2 (Fig. 1s, 1t), the general neuronal marker NeuN, or the general cellular marker DAPI (Supplementary Fig. 1e). In cortical regions that do not overlie the VaD lesion (layer II-V), no TFs were reduced (Supplementary 1f-1g), nor the number of neurons (Supplementary Fig. 1h, 1i). These indicated that VaD lesions caused transcriptomic changes in the adjacent brain area without affecting the total number of neurons, which may serve as a marker of disease effect in connected neuronal circuits.

### Cell-type specific RNA-Seq reveals activation of WM-associated aging genes in VaD mouse model

To identify the transcriptome of the cells in the neurovascular nice, intersecting viral and mouse transgenic approaches were used in translating ribosome affinity purification (TRAP) because it allows to study at great sequencing depth compared to scRNA or snRNAseq [20, 21]. Ribotag profiles the mRNA that is being transcribed, the “translatome”, which is more closely linked to the actual proteome in the cell than the total RNA sequencing of other approaches [22, 23]. To isolate mRNA from endothelial cells (ECs), intersectional viral approach (PHP.eB-CAG-Flex-Rpl22-HA) and transgenic mouse strain (Tie2-Cre) were employed in conjunction with TRAP [24] (Fig. 2a, 2b, 2i). Tbx18-CreER::Rpl22-HA mice were used to specifically isolate pericyte transcriptome using TRAP (Fig. 2c, 2d, 2i). Ng2-CreER::Rpl22-HA mice were used to specifically isolate OPCs (Fig. 2e, 2f, 2i). Lenti-GfaABC1D-Rpl22-HA was used to isolate astrocyte (Fig. 2g, 2h, 2i). Immunostaining with HA and cell type markers demonstrated specific labeling of EC (Glut1+), pericytes (Pdgfrβ+ or CD13+), OPCs (Olig2+Pdgfra+), astrocytes (GFAP+) respectively (Fig. 2a, 2c, 2e, 2g, Supplementary Fig. 2a, 2b, 2d, 2g, 2h). Real-time PCR further confirmed the specific enrichment of each cell type (Fig. 2b, 2d, 2f, 2h). Co-staining of OPC (Pdgfrα+) and myelinated oligodendrocyte (Aspa+) showed that HA labeling was highly specific to OPCs in VaD, which may due to limited differentiation after injury (Supplementary Fig. 2f, 2g). Although most canonical cell type specific markers reported in cortex [25] were enriched after TRAP, many of them were diminished in WM cells, e.g. OPC genes Chga, Nptx2, Olfm1, Grik3, and Garba3, etc., were reduced after TRAP in Ng2-CreER::Rpl22-HA mice (Fig. 2j), suggesting a distinct transcriptomic profile of WM cells. PCA analysis distinguishes the input and pulldown fractions from the control and VaD samples into different clusters (Fig. 2k, Supplementary Fig. 2i-k), establishing the validity of the pulldown approach. Differentially expressed genes (DEGs) were identified with an FDR < 0.1 (Fig. 2l).

**Figure 2.**
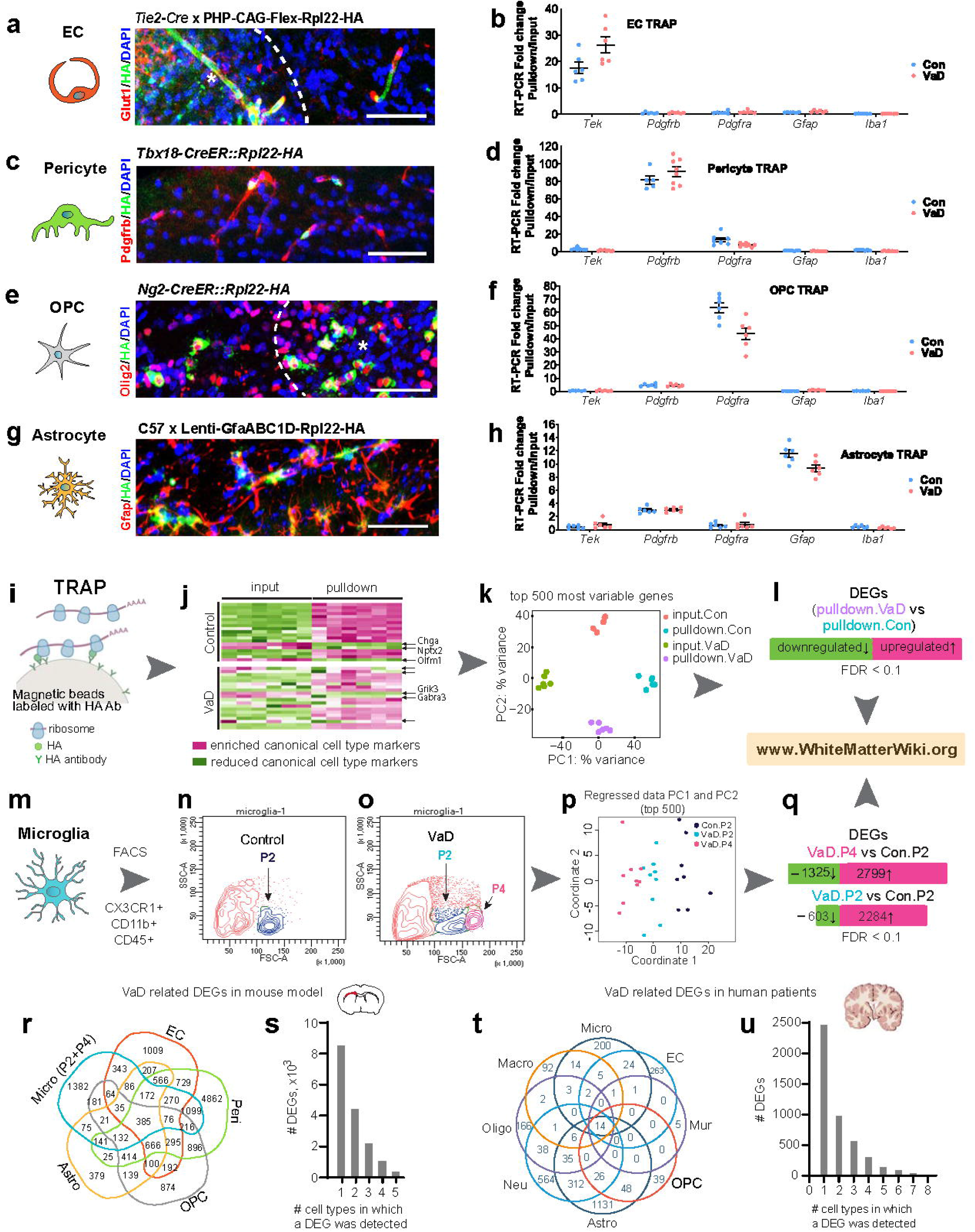
Cell type specific RNA-Seq of VaD associated DEGs in 5 white matter glia cells. (a, b) Purification of endothelial cell (EC) transcriptome by translating ribosome affinity purification (TRAP). (a) Representative immunostaining image shows the specific labeling of EC by retro-orbital injection of PHP-CAG-Flex-Rpl22-HA virus into Tie2-cre transgenic mice. Glut1 (red), HA (green), DAPI (blue); Star sign shows the VaD lesion core, dashed line shows the boarder. Scale bar = 50μm. (b) Real-time PCR results show the enrichment (fold change after TRAP, pulldown vs input) of only EC cell type marker gene (*Tek*) but not microglia (*Iba1*), astrocyte (*Gfap*), pericyte (*Pdgfrβ*), or OPC (*Pdgfrα*) in control and VaD. (c, d) Purification of pericyte transcriptome by TRAP. (c) Representative immunostaining image shows the specific labeling of pericyte using transgenic Tbx18-creER::Rpl22-HA mice. Pdgfrβ (red), HA (green), DAPI (blue); Scale bar = 50μm. (d) Real-time PCR results show the enrichment of only pericyte indicated by cell type marker gene (*Pdgfrβ*) but not OPC (*Pdgfrα*), astrocyte (*Gfap*), microglia (*Iba1*), or EC (*Tek*) in control and VaD. (e, f) Purification of OPC transcriptome by TRAP. (e) Representative immunostaining image shows the specific labeling of OPC using transgenic Ng2-creER::Rpl22-HA mice. Olig2 (red), HA (green), DAPI (blue); Scale bar = 50μm. (f) Real-time PCR results show the enrichment of only OPC indicated by cell type marker gene (*Pdgfrα*) but not pericyte (*Pdgfrβ*), astrocyte (*Gfap*), microglia (*Iba1*), or EC (*Tek*) in control and VaD. (g, h) Purification of astrocyte transcriptome by TRAP. (g) Representative immunostaining image shows the specific labeling of astrocyte by intracraniel injection of Lenti-GfaABC1D-Rpl22-HA virus into C57 mice. (h) Real-time PCR results show the enrichment of only astrocyte indicated by cell type marker gene (*Gfap*) but not OPC (*Pdgfrα*), pericyte (*Pdgfrβ*), microglia (*Iba1*), or EC (*Tek*) in control and VaD. (i) Schematic of how TRAP purify ribosomal mRNA using HA antibody. (j) Schematic shows white matter pulldown fraction after TRAP enrich/diminish canonical cell type marker genes in control and VaD. Diminished canonical cell type markers are indicated by arrows to the right. (k) Representative PCA result (from the OPC RNA-Seq results using Ng2-CreER::Rpl22-HA and TRAP) shows the clustering of input.Con, pulldown.Con, input.VaD, and pulldown.VaD fractions. Each dot represents an individual animal. (l) Schematic shows the identification of DEGs related to VaD by comparing pulldown.VaD and pulldown.Con samples in each cell type with FDR < 0.1. (m) Schematic shows the purification of microglia cells by FACS using CX3CR1, CD11b, CD45 antibodies from white matter. (n) Representative FACS plot shows the collection of P2 fraction as microglia sample from control brain. (o) Representative FACS plot shows the collection of P2 and P4 fraction as microglia samples from VaD brains. (p) PCA analysis of microglia samples from control and VaD brains. Each dot represents an individual animal. (q) Schematic shows the identification of microglial DEGs related to VaD by comparing VaD.P4 vs Con.P2, and VaD.P2 vs Con.P2 samples in each cell type with FDR < 0.1. The RNA-Seq results are available through www.WhiteMatterWiki.org. (r) Venn diagram shows the overlap of DEGs across 5 glia cell types in mouse VaD model. (s) Column graph shows the number of VaD associated DEGs expressed in one, or two, or multiple cell type(s) in mouse VaD model. (t) Venn diagram shows the overlap of DEGs across 8 glia cell types in human VaD. (u) Column graph shows the number of VaD associated DEGs expressed in one, or two, or multiple cell type(s) in human VaD.

To isolate microglia, a fluorescence-activated cell sorting (FACS) protocol for small WM regions using CX3CR1, CD11b, and CD45 antibodies was applied [26] (Fig. 2m). Based on cell scatter size, microglial obtained from the control brain was designated as P2 (Fig. 2n), while the same fraction and an additional fraction from VaD brain are named P2 and P4 respectively (Fig. 2o). RNA was extracted and sequenced, then clustered in PCA analysis (Fig. 2p). Border-associated macrophage (BAM) genes [27] were trace or undetectable indicating that the all FACS sorted cells were pure microglia (Supplementary Fig. 2l). Thousands of microglial DEGs were identified (VaD.P4 vs. Con.P2, and VaD.P2 vs. Con.P2) (Fig. 2q). Gene expression of 5 WM cells in control and VaD brains are accessible at www.WhiteMatterWiki.org. Taken together, most of the VaD related DEGs were cell type specific, and half of which are dysregulated in only one cell type in both mouse (Fig. 2r, s) and human (Fig. 2t, u) (Supplementary Table 1).

Glial aging is particularly accelerated in WM compared with cortical regions, while corpus callosum shows the most profound and earliest shifts towards aging [28]. Substantial numbers of WM-associated aging genes were changed in 5 types of glial cells (Fig. 3a) (Supplementary Table 2), mostly up-regulated, indicating that the VaD model in the young adult mouse caused a molecular expression profile that is seen in aging. Microglia exhibited the most unique shift of WM-associated aging genes (Fig. 3b), and most substantial fold change in VaD (Fig. 3c). Notably, the cluster of P4 microglia, exhibited a bigger number and greater change of WM-associated aging genes compared to P2-cluster microglia (Fig. 3c). A key WM-associated aging gene, C4b [28, 29], which is a complement component and major schizophrenia risk factor [30], is specifically up-regulated in OPC and microglia in VaD (Fig. 3c). Function analysis of KEGG pathways showed that microglial transcriptomic changes in the VaD model were associated with dysregulation of major molecular pathways, including those involved in Alzheimer’s disease (Fig. 3d). GO enrichment analysis also indicated that microglia in VaD model were involved in dysregulated molecular functional pathways including ATP binding (Fig 3e).

**Figure 3.**
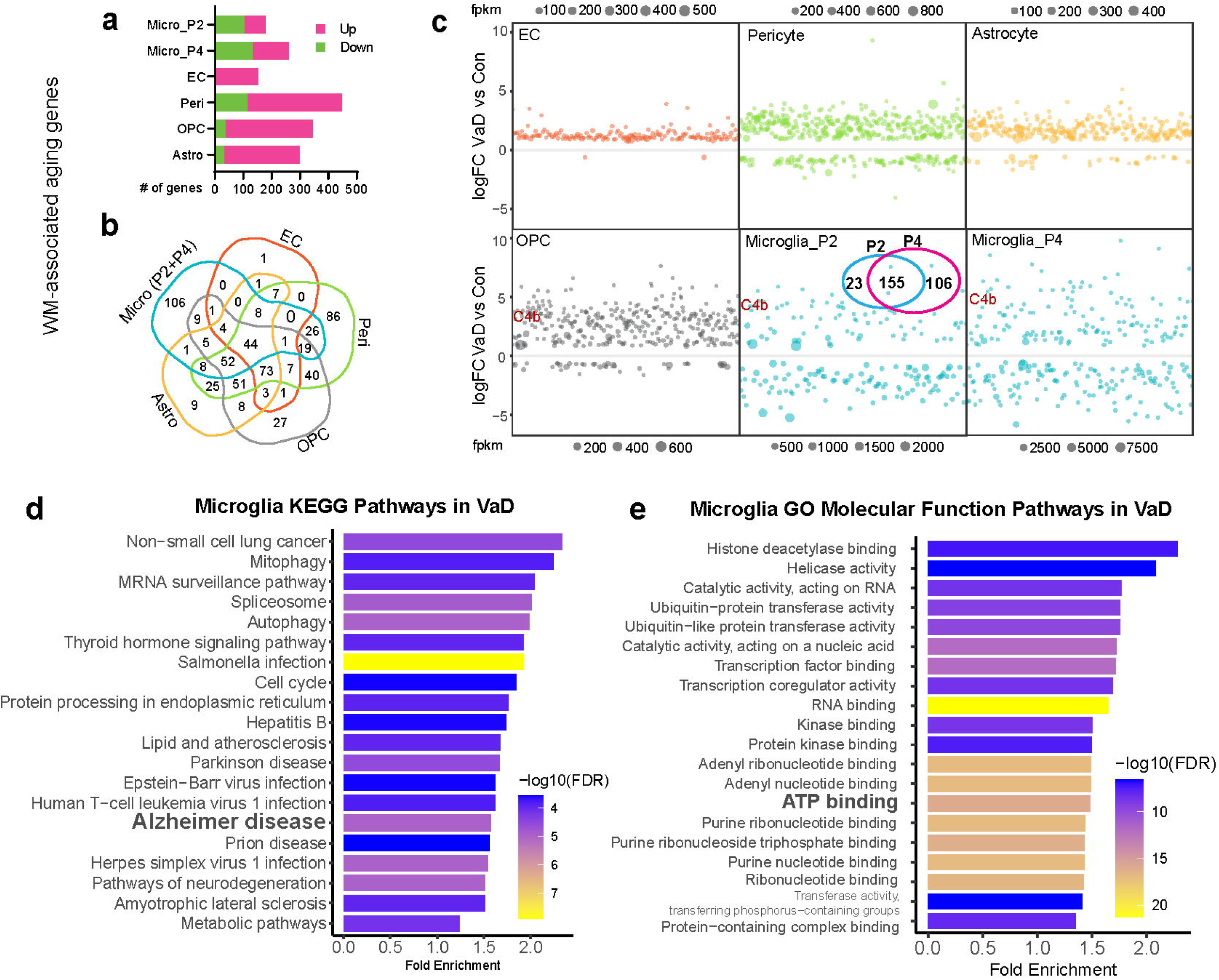
Cell type-specific white matter associated aging gene changes in VaD mouse model. (a) Number of WM-associated aging genes that are either up- or down-regulated in VaD in 5 glia cell types. (b) Venn graph shows the number of overlapped aging genes across 5 glia cell types. (c) Dot plots show the log fold change of aging genes in each cell type, VaD vs Con. The size of dots represents the expression level (FPKM) of genes. Venn plot in the square of Microglia_P2 shows the number of overlap and different aging gene changes in P4 vs P2 fractions in VaD. C4b is highlighted in maroon. (d) Microglia KEGG pathways analysis of aging genes changed in VaD mouse model. (e) Microglia GO molecular function pathways analysis of aging genes changed in VaD mouse model.

Many cell types have specific transcriptomes in distinct brain regions [26, 31-33]. A set of cell-type and WM-specific TFs/marker genes were identified (Supplementary Fig. 3a). Foxc1, Sox4, Atoh8, and Zfp871 are examples of distinct WM cell-type specific TFs compared to cortex and whole brain (Supplementary Fig. 3b, 3c). TFs are crucial for regulating gene expression. These findings suggest a distinct transcriptional control profile of WM cells against other brain regions. Cell-type marker genes specific for WM were also identified (Supplementary Fig. 3d). Tmem212, Slc1a1, Thbs4, and Nav3 are highly expressed in WM EC, OPC, astrocyte, and microglia respectively but not in cortical datasets (Supplementary Fig. 3e, 3f). Tmem212 is related to cerebral small vessel disease, and associated with enlarged periventricular space in magnetic resonance imaging (MRI) in aged populations [34]. VaD significantly increased Tmem212 expression in mouse [FPKM, 61.3±10.4 (control) vs 118.5±8.3 (VaD)]. Rank-rank hypergeometric overlap (RRHO) analyses [35] comparing WM cell type transcriptomes with cortical [25] and whole brain [36] show lower similarity of transcriptional profiles in EC and microglia specifically (Supplementary Fig. 4a-d). Furthermore, gene set enrichment analysis (GSEA) [37, 38] showed distinctive gene markers and functional hallmarks of WM EC and microglia compared with the same cell types in cortex (Supplementary 4e-h). These results highlight the importance of assembling a WM-specific cell transcriptome as a critical first step in identifying disease-associated genes in VaD, as well as other WM diseases.

**Figure 4.**
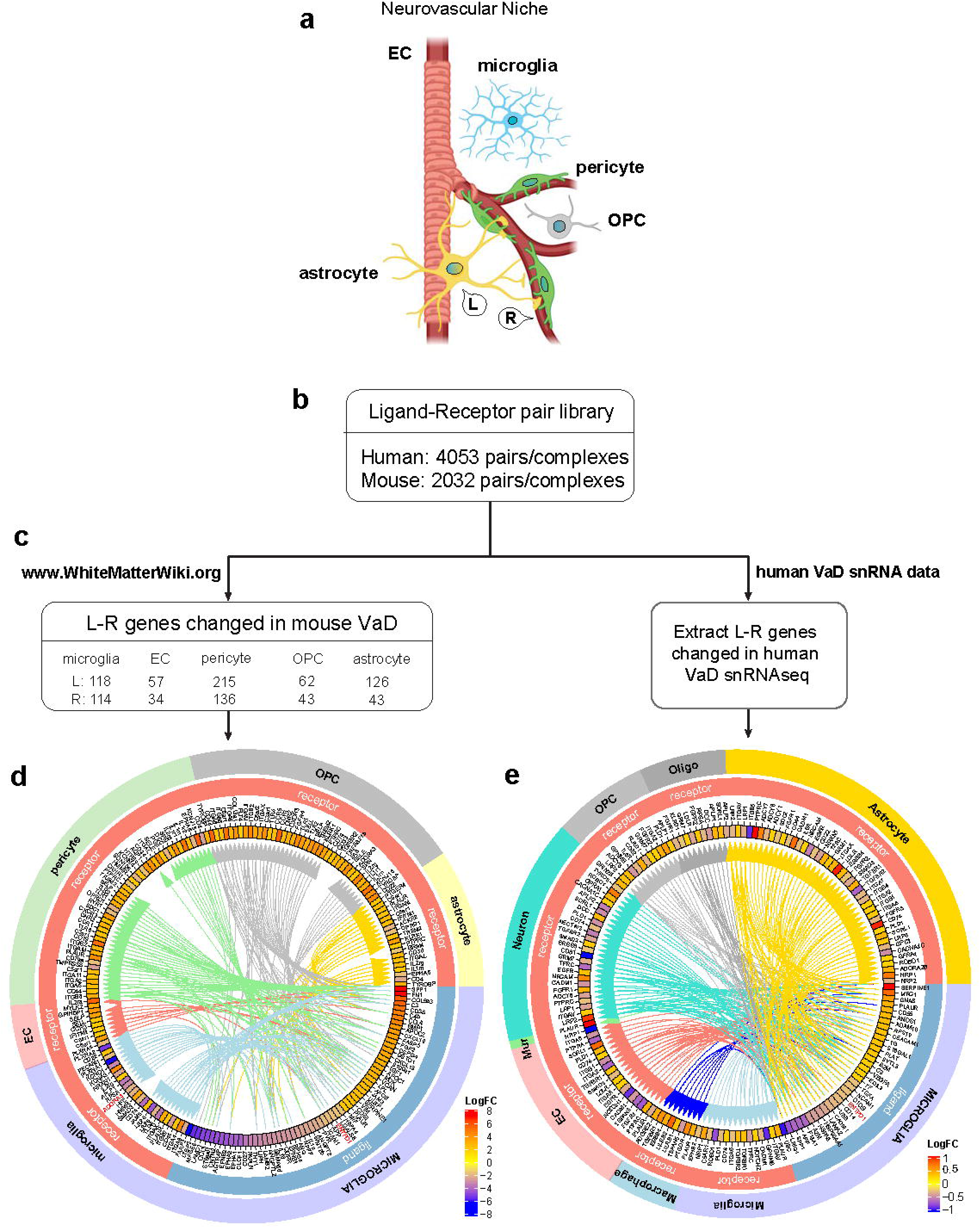
Identification of ligand-receptor (L-R) interactions in VaD neurovascular niche using L-R database in mouse VaD model and human patients. (a) Schematic of cell-cell interaction through L-R pairs in the neurovascular niche, generated by BioRender. (b) A custom-made L-R pair database which contains 4053 pairs in human and 2032 pairs in mouse. (c) Identification of L-R genes using L-R database from cell type-specific RNA-Seq of mouse VaD model and human VaD snRNA-Seq results. For example, from the mouse VaD RNA-Seq result, 118 ligands (including 84 microglia ligands were identified from the human ligand pool and another 34 microglia ligands) were identified from the mouse ligand pool. (d) Circos plot shows the intercellular interactome in mouse VaD brain from microglia to all the cell types. Note: genes with LogFC between -1.5 and 2 were excluded due to limited space in Circos plots. The name of genes will be indiscernible if all the genes are included. (e) Circos plot shows the intercellular interactome in human VaD brain from microglia to all the cell types. The Log fold change of each L/R gene is indicated in the gradient color box. The interacting cells are connected by arrows through L-R pairs.

### Constructing a map of the intercellular interactome in VaD

To determine the cell-cell signaling systems within the WM neurovascular niche (Fig. 4a), the VaD cell-type specific transcriptomes were analyzed for ligand-receptor (L-R) systems that are conjointly and differentially regulated. First, a L-R database was assembled by merging three major L-R libraries [39-41] covering 4053 human and 2032 mouse L-R pairs/complexes (Fig. 4b), which is currently the largest of its kind (Supplementary Table 3). Subsequently, the number of ligand and receptor genes related to mouse VaD were identified in each cell type. For example, from the microglia RNA-Seq data in mouse VaD, 118 ligands were identified including 84 from human L-R pool and additional 34 from mouse L-R pool (Fig. 4c). Human VaD related L-R genes were also identified from our published snRNA-Seq dataset [8].

To determine the intercellular interactome during VaD recovery, cells were linked together via L-R interactions (Supplementary Tables 4, 5), which can be visually presented by Circos plots (Fig. 4d, 4e, Supplementary Fig. 5). In mouse VaD, the top microglial ligands (LogFC > 2 or < -1.5) identified interfacing the human L-R pool, were connected towards corresponding receptors in other cell types, where SPP1 was the most increased ligand, and FARP2 was the most decreased ligand (Fig. 4d). In human VaD, the microglial ligands were identified interfacing the human L-R pool and connected towards corresponding receptors in other cell types (Fig. 4e).

**Figure 5.**
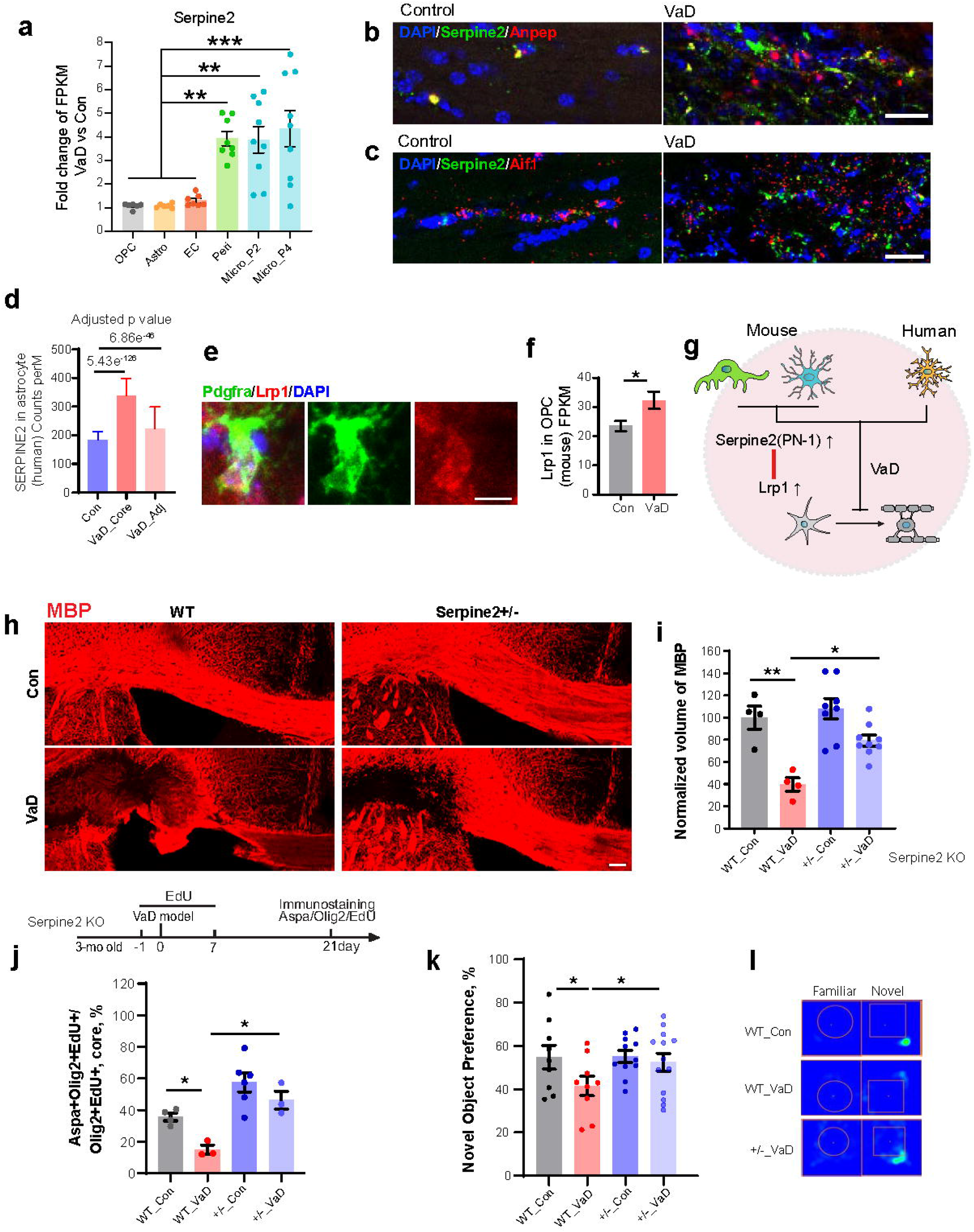
Intercellular Serpine2-Lrp1 pathway inhibits OPC maturation in VaD tissue repair. (a) The increase (FPKM, fold change) of Serpine2 expression in 5 glial cell types in VaD mouse model. (b) RNAscope images showing the Serpine2 (green) expression in pericyte (Anpep, red) cells in control and VaD white matter. Scale bar = 25μm. (c) RNAscope images showing the Serpine2 (green) expression in microglia (Iba1, red) cells in control and VaD white matter. Scale bar = 25μm. (d) The expression of SERPINE2 in human periventricular white matter under normal and VaD conditions. (e) Immunostaining images show the expression of Lrp1 in OPC in white matter. (f) The difference of Lrp1 expression (FPKM) in OPC in control and VaD mouse brain. * *p* < 0.05. (g) Schematic shows the hypothesis of intercellular Serpine2-Lrp1 signaling from pericyte/microglia (in mouse) or astrocyte (in human) to OPC, which inhibit differentiation of OPC towards myelinated oligodendrocyte. (h) Representative images show the immunostaining of MBP (red) in WT and Serpine2+/-animals under control and VaD conditions. Scale bar = 100μm. (i) Quantification of MBP volume in white matter of four animal cohorts: WT_con, WT_VaD, Serpine2+/-_Con, and +/-_VaD. WT_Con vs WT_VaD, ** *p* < 0.01; WT_VaD vs +/-_VaD, * *p* < 0.05. Each dot represents an individual animal. Two-way repeated measures ANOVA. (j) Schematic shows the paradigm of EdU labeling of OPCs (Olig2+) differentiated into myelinated oligodendrocytes (Aspa+Olig2+EdU+), and the quantification of new born OPCs differentiated in VaD lesion (Aspa+Olig2+EdU+/Olig2+EdU+, %). WT_Con vs WT_VaD and WT_VaD vs +/-_VaD, * *p* < 0.05. Each dot represents an individual animal. Two-way repeated measures ANOVA. (k) Result of novel object recognition test using WT and Serpine2 +/-animals. WT_Con vs WT_VaD and WT_VaD vs +/-_VaD, * *p* < 0.05. Each dot represents an individual animal. Two-way repeated measures ANOVA. (l) Representative heat maps show the exploration towards novel objects in WT_Con, WT_VaD, and +/-_VaD cohorts.

### Categorization of L-R genes as potential candidates for VaD treatment

In order to screen potential targets for VaD treatments, a series of molecular identity and significance criteria was used to categorize the DEGs in ascending order of priority for study (Supplementary Fig. 6a). First, the candidate must be a ligand (Tier 5). Second, the ligand candidates from the mouse VaD data set had to also be significantly regulated in human VaD (Tier 4). Next, ligand candidates comparably changed in only one or two cell types were prioritized (Tier 3). Then, those with significant change in corresponding receptors were identified (Tier 2). Finally, the candidates with a reported function related to neurological activity were classified (Tier 1) (Supplementary Table 6). In this selection scheme, Tier 1 L-R genes are the most attractive candidates under the scope of an intercellular interactome for VaD study.

**Figure 6.**
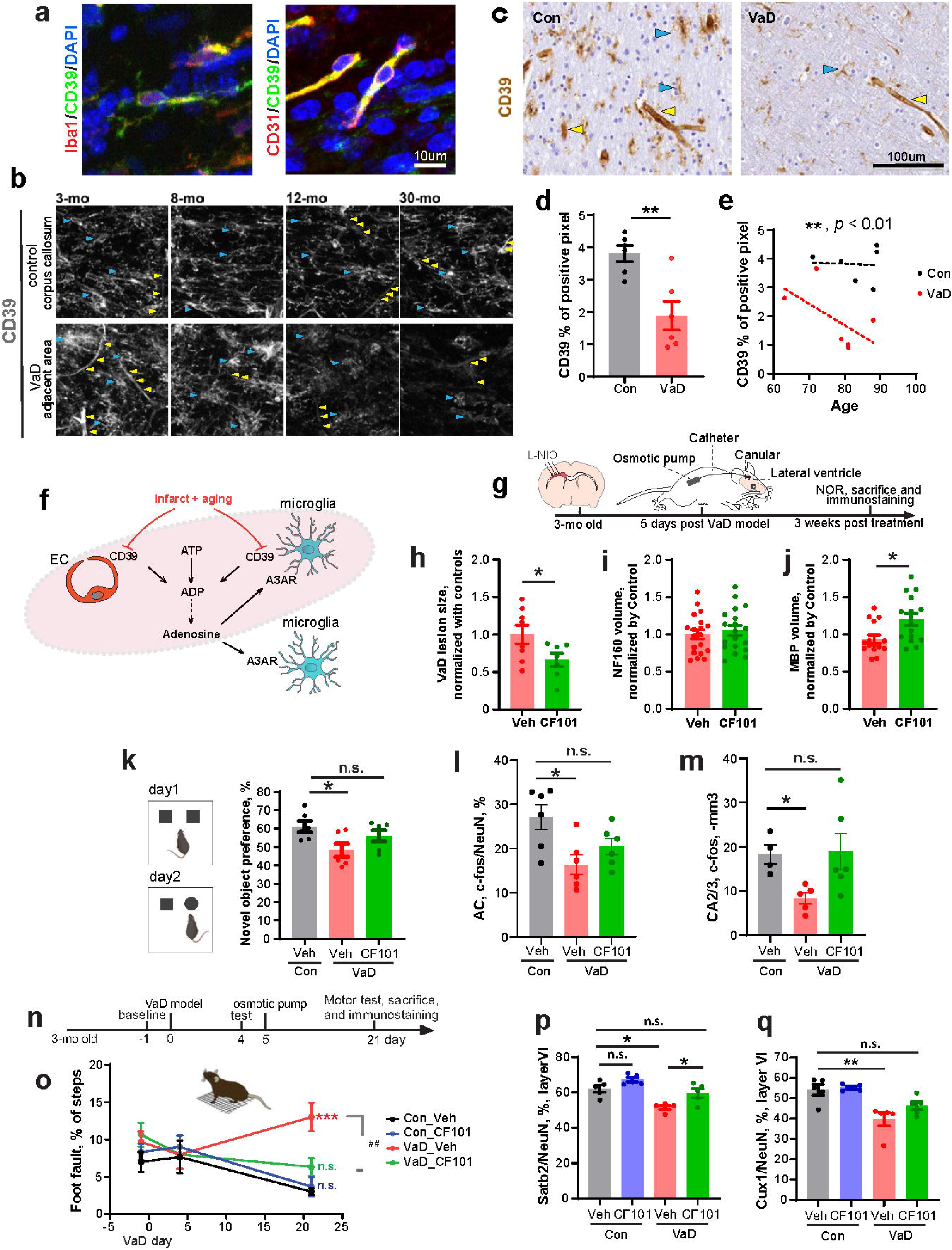
Intercellular CD39-A3AR signaling pathway is a potential target for enhancing VaD tissue and behavior recovery. (a) Representative immunostaining images of CD39 (green) showing the specific expression in microglia (Iba1, red), and EC (CD31, red) in mouse cerebral white matter. DAPI, blue; scale bar = 10μm. (b) Representative images showing the change of CD39 immunoreactivity in control and VaD brains at different ages (3-, 8-, 12-30-months). (c) Representative images of CD39 IHC in normal and VaD human brains. (d) Quantification of CD39 immunoreactivity in human periventricular white matter by percentage of positive pixel per ROI. ** *p* < 0.01, t test. (e) Regression of CD39 immunoreactivity against ages of samples in control and VaD brains. Simple linear regression, VaD vs Con ** *p* < 0.01. (f) Schematic shows the hypothesis of intercellular CD39(EC/microglia) to A3AR (microglia) signaling in VaD lesion and adjacent area that are downregulated by the combination of infarct and aging. (g) Schematic shows the paradigm of CF101 treatment 5-days after VaD induction, and the examination of cognitive function and tissue repair 3-weeks after treatment. (h) Quantification of VaD lesion size. Each dot represents an individual animal. * *p* < 0.05, t test. (i) Quantification of axon volume as evaluated by NF160 immunostaining. (j) Quantification of myelin volume as evaluated by MBP immunostaining. Each dot represents an individual animal. * *p* < 0.05, t test. (k) Result of novel object recognition test. Con_Veh vs VaD_Veh, * *p* < 0.05; Con_Veh vs VaD_CF101, n.s. = not significant. Each dot represents an individual animal. Two-way repeated measures ANOVA. (l) Quantification of c-Fos/NeuN (%) in anterior cingulate (AC). Con_Veh vs VaD_Veh, * *p* < 0.05; Con_Veh vs VaD_CF101, n.s. = not significant. Each dot represents an individual animal. Two-way repeated measures ANOVA. (m) Quantification of c-Fos/NeuN (%) in hippocampal CA2/3 subregion. Con_Veh vs VaD_Veh, * *p* < 0.05; Con_Veh vs VaD_CF101, n.s. = not significant. Each dot represents an individual animal. Two-way repeated measures ANOVA. (n) Schematic shows the paradigm of CF101 treatment 5-days after VaD induction, and the examination of motor function and the change of subtypes of neurons using immunostaining 3-weeks after treatment. (o) The results of the grid walk task. Three weeks post CF101 treatment, Con_Veh vs VaD_Veh, *** *p* < 0.001; Con_Veh vs Con_CF101, and Con_Veh vs VaD_CF101, n.s. = not significant. 8 animals per group. (p) Quantification of Satb2+/NeuN (%) in Layer VI above infarct lesion. Con_Veh vs VaD_Veh, and VaD_Veh vs VaD_CD101 * *p* < 0.05; Con_Veh vs VaD_CF101, n.s. = not significant. Each dot represents an individual animal. Two-way repeated measures ANOVA. (q) Quantification of Cux1+/NeuN (%) in Layer VI above infarct lesion. Con_Veh vs VaD_Veh, ** *p* < 0.01; Con_Veh vs VaD_CF101, n.s. = not significant. Each dot represents an individual animal. Two-way repeated measures ANOVA.

To classify the L-R candidates in terms of functions, ECM and G-protein coupled-receptor (GPCR) were identified as two major groups of candidates (Supplementary Fig. 6b, 6c). Microglia and pericyte are more involved in the regulation of ECM components, including the Tier 1 candidates: ECM regulating gene Entpd1 (CD39) and Serpine2 (PN-1) (Supplementary Fig. 6b). Microglia also exhibited higher dysregulation of GPCRs, including the Tier 1 receptor: Adora3 (A3AR) (Supplementary Fig. 6c).

### Serpine2-Lrp1 intercellular pathway inhibits OPC maturation in VaD tissue repair

Serpine2 was significantly increased in pericytes and microglia in mouse VaD (Fig. 5a-c) and astrocytes in human VaD (Fig 5d). Serpine2-encoded protease nexin-1 (PN-1) plays a role in multiple sclerosis [42] and Alzheimer’s disease [43, 44]. It binds to low density lipoprotein receptor-related protein 1 (Lrp1) to mediate functions in cancer [45] and vascular biology [46]. Lrp1 expression in OPC is higher than in myelinated oligodendrocytes [47]. Its expression in OPC was further validated using immunostaining (Fig 5e). RNA-Seq showed a significant increase of Lrp1 expression in VaD in OPC (Fig. 5f). Therefore, the Serpine2-Lrp1 axis was overall up-regulated, and may play a role in the differentiation of OPCs into myelinating oligodendrocytes which is required for VaD tissue repair (Fig. 5g). To investigate the function of Serpine2 on OPC differentiation, remyelination was evaluated in Serpine2 KO mice during VaD recovery. Homozygous knockouts were excluded, considering their significant epileptic activity and impairment of memory associated-long term potentiation (LTP) [48]. Immunostaining of myelin basic protein (MBP) showed that the reduction of myelin volume in VaD was ameliorated by reduced Serpine2 expression in heterozygous siblings (Fig. 5h, 5i). Quantification of myelinating oligodendrocytes (Aspa+Olig2+EdU+) differentiated from newly formed OPCs (Olig2+ EdU+) in the infarct core indicated that the differentiation of newborn OPC after VaD was increased by Serpine2 deficiency (Fig. 5j). Strikingly, the NOR task showed that reduced Serpine2 expression rescued the memory deficit in VaD (Fig. 5k, 5l). To summarize, Serpine2 plays a key role in the WM damage that underlies VaD, and its loss in this model promotes progenitor responses toward myelination and restores memory function.

### CD39-A3AR intercellular pathway is a potential target for tissue and behavior repair in VaD

CD39 (Entpd1) is an ectonucleotidase that plays a significant role in regulating the balance of extracellular ATP [49]. It has potential as a cancer immunotherapeutic target. Blockade of CD39 may prevent cancer cell adhesion to the ECM [50]. Cell type-specific RNA-Seq shows that CD39, and its receptor A3AR (Adenosine A3 receptor, gene name Adora3) were specifically expressed in microglia, and significantly reduced in VaD (Supplementary Fig. 7a, 7b). CD39 plays a role in the negative feedback control of neuronal activity by microglia through A1AR [51]. However, A1AR is not significantly changed in VaD (Supplementary Fig. 7e). Extracellular signaling in CD39-A3AR occurs as ATP is converted by CD39 to ADP and AMP; AMP is converted by CD73 (Nt5e) to adenosine; adenosine activates A3AR and is degraded by adenosine deaminase (Ada). RNA-Seq data indicated low expression of Nt5e (FPKM ≤ 3), and no significant change in Nt5e and Ada (Supplementary Fig. 7f, 7g). These data position extracellular ATP and its regulation by CD39 in VaD.

The cellular localization and protein levels in the CD39/A3AR system were determined in mouse and human, and as a function of age and VaD. Co-staining of CD39 with cell specific markers show that CD39 was specifically expressed in microglia and EC (Fig. 6a). Aging alone (3-mo to 30-mo in the mouse) did not lead to reduction of CD39 expression, while aging and VaD together significantly reduced CD39 expression in EC and microglia (Fig. 6b). Human VaD snRNA-Seq showed that microglial and endothelial CD39 expression in the WM lesion core and adjacent area was significantly reduced (Supplementary Fig. 7c, 7d, derived from [8]). Immunohistochemistry (Fig. 6c) further validated a significant reduction (50.6%) of CD39 expression in VaD brains (Fig. 6d). In contrast, the immunoreactivity of Glut1 (EC marker) was reduced only by 20.3%, which may be due to ischemic injury in vessels, while Iba1 (microglia marker) was not significantly changed (Supplementary Fig. 7h-k). Therefore, reduced CD39 immunoreactivity in human VaD is caused by its impaired expression in EC and microglia, including some decreased volume of blood vessels. Notably, the reduction of CD39 expression was inversely related to age in human VaD (Fig. 6e), which indicates that aging and VaD synergistically reduce CD39 expression.

These data suggest that intercellular CD39-A3AR signaling is an endogenous mechanism associated with VaD: Endothelial and microglial CD39 enhance the conversion of ATP into adenosine, which modulates microglia through A3AR; Infarct and aging together reduce CD39 expression in EC and microglia during VaD, which lead to impaired signaling to microglia in the infarct core and adjacent area, and thereafter retarded tissue and behavioral repair (Fig. 6f). In the studies of ischemic stroke, A3AR plays a neuroprotective role in the central nervous system [52]. Chronic administration of an A3AR agonist before global brain ischemia improved post-ischemic cerebral blood circulation, survival, and neuronal preservation [53]. However, the possibility of a delayed action of this signaling axis during the progression of WM ischemia/VaD has not been determined.

Immunostaining validated the expression of A3AR in WM microglia (Supplementary Fig. 8a) and the reduction of Adora3 in microglia was validated by RNAScope (Supplementary Fig. 8b, 8c). To investigate whether increasing an affected CD39-A3AR signaling is beneficial to VaD recovery, CF101, an A3AR-specific agonist, was delivered either acutely or in a delayed manner after VaD induction. CF101 is 50-fold more potent in A3AR action than with A1AR and A2bAR [54], and has been safely used in humans [55] in recent phase III clinical trials for psoriasis [56, 57]. Acute administration of CF101 from 30-min post stroke, for 1-2 weeks, by direct infusion into the lateral ventricle, reduced blood brain barrier (BBB) leakage (Supplementary Fig. 8d-f). However, unlike large artery stroke, periventricular WM microinfarcts in the human are often asymptomatic and progresses for years before development of VaD symptoms, so that any medication in VaD is likely to be a delayed treatment. Delayed CF101 delivery (Fig. 6g) produced a significant reduction of lesion size (Fig 6h). Although measures of axonal projections were not changed (Fig 6i), markers of myelination were significantly increased in affected WM by delayed CF101 treatment (Fig. 6j). To the best of our knowledge, this is the first finding of a drug that reduces lesion progression or enhances tissue repair in such a delayed treatment. Importantly, the NOR memory test showed that CF101 significantly reduced memory deficits in VaD animals (Fig. 6k), which also included restored c-Fos expression in PFC (AC) and HPC (CA2/3) (Fig. 6l, 6m).

Human VaD is associated with gait and motor abnormalities that predict dementia status and are associated with morbidity [58-60]. Using grid-walking, which tests limb motor control, delayed CF101 treatment successfully rescued the motor deficit in VaD (Fig. 6n, 6o). The reduced expression of neuronal transcription factors, Satb2 and Cux1, in the deep layers of motor somatosensory cortex, adjacent to the WM VaD lesion is a marker of secondary injury effect in this model (Fig. 1s, 1t; Supplementary Fig. 1e). Immunostaining using the animal cohorts after grid-walking tests showed that the reduced Satb2 and Cux1 expression, were significantly ameliorated by CF101 treatment (Fig. 6p, 6q).

## Discussion

Vascular dementia (VaD) produces disability and reduces quality of life, with a rising incidence due to an aging population. VaD arises from compromised cerebral blood flow, particularly ischemia and/or infarction in white matter (WM), which are often initially asymptomatic, resulting in tissue damage and cognitive decline. Unlike Alzheimer’s disease with established transgenic models, VaD lacks suitable counterparts, hindering therapeutic progress. The VaD mouse model induced by WM ischemia faithfully reproduces human VaD’s complex pathophysiology, featuring consistent lesions, in the most common location. It mirrors human VaD, exhibiting neural circuit damage and cognitive deficits. Akin to multiple sclerosis, where demyelination leads to decreased expression of specific neuronal transcription factors or subtypes [19], this VaD model also shows specific reduction of Satb2 and Cux1in neurons near lesions. While offering advantages, the model does not incorporate or mimic related genetic diseases, blood-brain barrier dysfunction [61], hippocampal cell death [62], small haemorrhages, and vascular amyloid deposition [63].

This study using TRAP brings new evidence which underscores a unique WM’s transcriptomic profile compared to gray matter in a cell type specific manner [64-66]. This study also fills the gap of a comprehensive WM transcriptomic analysis in normal and VaD brains (www.WhiteMatterWiki.org). Moreover, additional WM-specific cell type markers and transcription factors were identified when compared to cell-specific cortical tissue [25] and whole brain datasets [36].

For translational relevance, it is ideal to identify intercellular molecular signaling systems that are differentially regulated in both mouse and human VaD: a VaD “interactome”. In the present study, target screening criteria include whether the gene is significantly regulated in both human and mouse RNA-Seq, whether the change is specific to one or two particular cell type(s), and whether the known function is implicated in CNS. Several the highest value (Tier 1) L-R candidates were studied in our VaD mouse model. This is a first report of the role of Serpine2-Lrp1 in OPC differentiation, and the ability of downregulating Serpine2 to enhance tissue repair (remyelination) and memory recovery in VaD. The limitation of this Serpine2 study is that conventional knockout animals are used, which means cell type specific modification will be the future direction for further investigation. CD39-A3AR exhibits specific expression in endothelial cells and microglia. Notably, the expression of CD39 remains unaffected solely by the aging process but is influenced by the combined effects of aging and VaD. A3AR is the functional receptor of CD39, and it exhibits specific expression in microglia and experiences a significant reduction in VaD. Notably, the adult brain exhibits limited restorative capabilities, particularly in WM. Initially, infarct lesions in WM are often asymptomatic, yet progress into adjacent areas, resulting in more severe impairments [67]. Delayed treatment with an A3AR agonist significantly enhanced tissue repair and functional recovery. This delayed delivery holds clinical relevance to the disease presentation. CF101, a highly selective A3AR agonist undergoing phase III clinical trials for psoriasis, exhibits minimal non-specific effects on other adenosine receptors. The approaches in this set of studies were a discovery-based progression from large-scale molecular expression analysis in human and mouse VaD. The current investigations represent the first demonstration of a pharmacological approach capable of enhancing tissue repair in the VaD brain.

## Author contributions

M. Tian and S. T. Carmichael initiated, designed the study, and drafted the manuscript. M. Tian conducted and analyzed most of the experiments, including construction of custom L-R database and drawing of Circos plots. R. Kawaguchi conducted the DEGs analysis for cell type specific RNA-Seq. Y. Shen conducted the FC and NOR tests. M. Machnicki, produced PHP.eB-CAG-Flex-Rple22-HA and Lenti-GfaABCD-Rpl22-HA viruses. N. G. Villegas, D. R. Cooper, N. Montgomery, J. Haring, R Lan, and A. H. Yuan, conducted part of immunostaining and image quantification. C. K. Williams, S. Magaki, and H. V. Vinters, helped with human sample processing, IHC and quantification. L. M. De Biase and M. Tian conducted the microglia FACS. Y. Zhang directed the website development, and the participated in Serpine2 study. A. J. Silva helped with project designing and behavior test designing. M. Tian and R. Kawaguchi have the direct access to RNA-Seq data. S. T. Carmichael is the senior and corresponding author.

## Acknowledgement

This project was supported by Ressler Family Foundation, NIH R37NS102185, The Dr. Miriam and Sheldon G. Adelson Medical Research Foundation, UCLA Eli and Edythe Broad Center of Regenerative Medicine and Stem Cell Research, including support from the Steffy Family Trust. We thank Dr. Daniel T. Gray, Dr. André Sousa for advice; Ms. Pooja Nair, Dr. Mitchell Krawczyk for technical support.

---

**Supplementary Figure 1 Other characteristics of mouse VaD model (related to Figure 1)**. (a) Immunostaining images showing the damage of axon (NF160, green), activation of astrocyte (GFAP, gray), and damage of myelin (MBP, red). DAPI, blue; Star sign indicates the lesion core; scale bar = 100μm. (b) Immunostaining images showing the pericyte (CD13, red) response in VaD lesion. DAPI, blue; Star sign indicates the lesion core; scale bar = 50μm. (c) Immunostaining image shows the injection of retroAAV-PGK-cre in the hippocampus spread across all the subregions (CA1, CA2/3, and DG) as indicated by the expression of cre recombinase (red). (d) Quantification of Cux1+, NeuN+, DAPI+ cell number in cortical Layer VI under control and VaD conditions. Cux1, Con vs VaD, * *p* < 0.05, t test, each dot represents an individual animal. (f) Quantification of Cux1+ cell numbers in Layer II/III, IV, and V. Each dot represents an individual animal. (g) Quantification of Satb2+ cell numbers in Layer V. Each dot represents an individual animal. (h) Quantification of Ctip2+ cell numbers in Layer V. Each dot represents an individual animal. (i) Quantification of NeuN+ cell numbers in Layer II/III, IV, and V. Each dot represents an individual animal. (j) Summary of the VaD mouse model with human VaD pathological characteristics.

**Supplementary Figure 2 Cell-type specific HA labeling for TRAP, and low macrophage contamination in FACS sorted microglia (related to Figure 2)**. (a) Immunostaining images show that EC HA (green) labeling, using Tie2-cre x PHP-CAG-Flex-Rpl22-HA, does not get into microglia (Iba1, red), astrocyte (Gfap, red), pericyte (Pdgfrβ, red), or oligodendrocyte lineage (Olig2, red). DAPI, blue; Scale bar = 50μm; Star sign indicates the lesion core; Dashed line shows the border of infarct. (b) Immunostaining images show that pericyte HA (green) labeling, using Tbx18-creER::Rpl22-HA transgenic mice, does not get into microglia (Iba1, red), astrocyte (Gfap, red), EC (CD31, red), or oligodendrocyte lineage (Olig2, red). DAPI, blue; Scale bar = 50μm; Star sign indicates the lesion core; Dashed line shows the border of infarct. (c) The paradigm of tamoxifen injection and VaD induction in mice cohorts for pericyte and OPC labeling. (d) Immunostaining images show that OPC HA (green) labeling, using Ng2-creER::Rpl22-HA transgenic mice, does not get into microglia (Iba1, red), astrocyte (Gfap, red), pericyte (CD13, red), or EC (Glut1, red). DAPI, blue; Scale bar = 50μm; Star sign indicates the lesion core; Dashed line shows the border of infarct. (e) Quantification of HA+Olig2+/HA+ (%) in the cerebral white matter of Ng2-creER::Rpl22-HA transgenic mice. (h) Immunostaining images show that astrocyte HA (green) labeling, using C57 x Lenti-GfaABC1D-Rpl22-HA, does not get into microglia (Iba1, red), EC (CD31, red), pericyte (CD13, red), or oligodendrocyte lineage (Olig2, red). DAPI, blue; Scale bar = 50μm; Star sign indicates the lesion core. (f) Schematic shows the region of cerebral white matter where the oligodendrocyte lineage cells were investigated in VaD and intact conditions. (g) Representative images show the colocalization of HA expression with OPC (Pdgfrα, red), total lineage (Olig2, red), and myelinated oligodendrocyte (Aspa+, red) in intact brain, infarct core, and peri-infarct white matter in VaD. Scale bar = 50μm. (i-k) PCA results show the clustering of input.Con, pulldown.Con, input.VaD, and pulldown.VaD fractions after TRAP enrichment of pericyte (j), EC (k), and astrocyte (l); Each dot represents an individual animal. (l) Comparison of macrophage marker (Mrc1, Pf4, F13a1) and microglia marker (Tmem119, P2ry12, Cx3cr1, and Aif1) in FACS sorted cells in terms of FPKM.

**Supplementary Figure 3 Identification of white matter specific transcriptional factors and cell markers**. (a) A partial list of white matter specific TFs in 5 glial cell types, comparing their normalized specificity in white matter and cortex/whole brain. (b) The expression (FPKM) of example TFs: Foxc1 in pericytes, Mycl in OPCs, Atoh8 in astrocytes, and Zfp871 in microglia. (c) These TFs are not specific in corresponding cell types in the whole brain or cortical RNA-Seq. (d) A partial list of WM specific cell type markers, comparing their normalized specificity in white matter and cortex/whole brain. (e) The expression (FPKM) of example WM specific cell type markers: Tmem212 in EC, Slc1a1 in OPC, Thbs4 in astrocyte and Nav3 in microglia. (f) These markers are not specific in cortical cells.

**Supplementary Figure 4 RRHO and GSEA analyses of white matter RNA-Seq dataset compared with cortical/whole brain RNA-Seq datasets indicate specific transcriptomic characteristics of white matter cells**. (a) Pearson correlation coefficient comparing white matter with cortex cells using RRHO. White matter OPC and EC are fairly distinctive from cortical OPC and EC respectively. (b) Rank-Rank scatter plot shows the EC-related gene-expression signatures with medium overlap in white matter and cortex. (c) Pearson correlation coefficient comparing white matter with whole brain cells using RRHO. White matter microglia and EC are medium to small correlation with whole brain microglia and EC respectively. (e) GSEA analysis comparing EC from white matter and cortical cell RNA-Seq shows the top up-regulated (left) and down-regulated (right) genes in white matter. Five samples from white matter and 2 samples from cortex were compared. (e) GSEA analysis comparing microglia from white matter and cortical cell RNA-Seq shows the top up-regulated (left) and down-regulated (right) genes in white matter. Five samples from white matter and 2 samples from cortex were compared. (f) GSEA enrichment plots show the white matter EC is highly enriched in reactome translation and rRNA processing functions. (g) GSEA enrichment plots show the white matter microglia is highly enriched in fatty acid metabolism, hypoxia, and complement functions.

**Supplementary Figure 5 Circos plots show the intercellular interactome in mouse and human VaD (related to Figure 4)**. (a) Signaling from microglia to other cells that are changed in mouse VaD, identified by mouse pool of L-R library. (b) Signaling from microglia to other cells that are changed in human VaD, identified by mouse pool of L-R library. (c) Signaling from EC to other cells that are changed in mouse VaD, identified by human pool of L-R library. (d) Signaling from EC to other cells that are changed in mouse VaD, identified by mouse pool of L-R library. (e) Signaling from EC to other cells that are changed in human VaD, identified by human pool of L-R library. (f) Signaling from EC to other cells that are changed in human VaD, identified by mouse pool of L-R library. (g) Signaling from pericyte to other cells that are changed in mouse VaD, identified by human pool of L-R library. (h) Signaling from pericyte to other cells that are changed in mouse VaD, identified by mouse pool of L-R library. (i) Signaling from mural cell to other cells that are changed in human VaD, identified by human pool of L-R library. (h) Signaling from mural cell to other cells that are changed in human VaD, identified by mouse pool of L-R library. (k) Signaling from OPC to other cells that are changed in mouse VaD, identified by human pool of L-R library. (l) Signaling from OPC to other cells that are changed in mouse VaD, identified by mouse pool of L-R library. (m) Signaling from OPC to other cells that are changed in human VaD, identified by human pool of L-R library. (n) Signaling from OPC to other cells that are changed in human VaD, identified by mouse pool of L-R library. (o) Signaling from astrocyte to other cells that are changed in mouse VaD, identified by human pool of L-R library. (p) Signaling from astrocyte to other cells that are changed in mouse VaD, identified by mouse pool of L-R library. (q) Signaling from astrocyte to other cells that are changed in human VaD, identified by human pool of L-R library. (r) Signaling from astrocyte to other cells that are changed in human VaD, identified by mouse pool of L-R library. The Log fold change of each L/R gene is indicated in the gradient color box. The interacting cells are connected by arrows through L-R pairs.

**Supplementary Figure 6 Screening criteria for L-R candidates and the identification of VaD associated ECM/GPCR genes**. (a) The L-R pairs are classified into 6 tiers by whether the ligand is changed in human RNA-Seq, whether the dysregulated ligand expression is specific in one or two cell type(s) (based on mouse RNA-Seq data in control or VaD brain), whether the expression of corresponding receptor is also changed, and whether the L-R is interesting in brain function. One sample L-R in Tier5 is Serpine2 (↑)-Lrp1 (↑); one sample in Tier1 is Entpd1 (↓)-Adora3(↓). (b) Bubble plot shows the expression of ECM genes in 5 glia cell types. The gradient color shows how they are changed (logFC) in VaD. The bubble size shows the FPKM value of each gene in control brains. (c) Bubble plot shows the expression of GPCR genes in 5 glia cell types. The gradient color shows how they are changed (logFC) in VaD. The bubble size shows the FPKM value of each gene in control brains.

**Supplementary Figure 7 (related to Figure 6)**. (a) Expression level (FPKM in RNA-Seq) of Entpd1 (CD39) in 5 glial cells types in control and VaD brains. (b) Expression level (FPKM in RNA-Seq) of Adora3 (A3AR) in 5 glial cells types in control and VaD brains. (c) The expression level (counts perM read in snRNA-Seq) of Entpd1 (CD39) in microglia of normal brain periventricular white matter, VaD core and adjacent white matter tissue of human patients. (d) The expression level (counts perM read in snRNA-Seq) of Entpd1 (CD39) in EC of normal brain periventricular white matter, VaD core and adjacent white matter tissue of human patients. (e) Representative images of Glut1 IHC in normal and VaD human brains, which indicate the change of vessel volume in VaD. (g) Quantification of Glut1 immunoreactivity by positive pixel per ROI. \**p* < 0.05, t test. (f) Representative images of Iba1 IHC in normal and VaD human brains, which show the volume of microglia. (h) Quantification of Iba1 immunoreactivity by positive pixel per ROI. (i) Expression level (FPKM in RNA-Seq) of Nt5e (CD73) in 5 glial cells types in control and VaD brains. (j) Expression level (FPKM in RNA-Seq) of Ada (adenosine deaminase) in 5 glial cells types in control and VaD brains. (k) Expression level (FPKM in RNA-Seq) of Adora1 (A1AR) in 5 glial cells types in control and VaD brains.

**Supplementary Figure 8 (related to Figure 6)**. (a) Immunostaining images show the expression of A3AR (green) in microglia (Iba1, red). Star sign shows the core of VaD lesion; Scale bar = 10μm. (b) RNAscope images showing the Adora3 (green) expression in microglia (Iba1, red) cells in control and VaD white matter. DAPI = blue; Scale bar = 10μm. (c) Quantification of Adora3 dots per cell in control and VaD brains. Each dot represents an individual cell. *** *p* < 0.001, t test. (d) Schematic shows the CF101 treatment 30-min after VaD induction, and the examination of lesion size and BBB leakage using immunostaining 1∼2-weeks after treatment. (e) Quantification of VaD lesion size in VaD brains treated with vehicle or CF101. Each dot represents an individual animal. (f) Quantification of BBB leakage by immunoreactivity of mouse IgG. Each dot represents an individual animal; Veh vs CF101, * *p* < 0.05, t test.

**Supplementary Figure 9 RIN value, unique mapped reads (%), and number of reads for RNA-Seq samples**. (a) The values for EC samples. (b) The values for pericyte samples. (c) The values for OPC samples. (d) The values for astrocyte samples. Each dot represents an individual animal.

Supplementary Table 1. VaD related DEGs across cell types.

Supplementary Table 2. Aging related genes that are dysregulated in mouse VaD model.

Supplementary Table 3. L-R database.

Supplementary Table 4. VaD interactome identified by human L-R pool.

Supplementary Table 5. VaD interactome identified by mouse L-R pool.

Supplementary Table 6. Tier 1_L-R candidates.

