## Supplementary material for "Intercellular Signaling Pathways as Therapeutic Targets for Vascular Dementia Repair": Online Methods

### Animals

All experiments were performed in accordance with National Institutes of Health (NIH) animal protection guidelines and were approved by University of California, Los Angeles Chancellor's Animal Research Committee. Tie2-cre transgenic mice (B6.Cg-Tg(Tek-cre)1Ywa/J) were purchase from the Jackson Laboratories. C57BL/6J (C57) mice were purchase from the Jackson Laboratories or Taconic Biosciences. Tbx18-creER transgenic strain [1] (gift from Sylvia Evans, University of California, San Diego, CA) was crossed with Rpl22-HA transgenic mice (B6J.129(Cg)-Rpl22tm1.1Psam/SjJ, Jackson Laboratory) to obtain the Tbx18 (T-box transcription factor 18) reporter mouse line Tbx18-creER::Rpl22-HA animals. Ng2-creER transgenic strain [2] (gift from Akiko Nishiyama, University of Connecticut, Storrs, CT) was crossed with Rpl22-HA transgenic mice (B6J.129(Cg)-Rpl22tm1.1Psam/SjJ, Jackson Laboratory) to obtain the OPC reporter mouse line Ng2-creER::Rpl22-HA animals. *Serpine2* KO mice [3] were obtained from Dr. Ye Zhang at UCLA.

### Virus constructs

AAV1-CAG-FLEX-EGFP-WPRE (Addgene 51502, >7x$10^{12}$ vg/mL) and retroAAV-PKG-cre (Addgene 24593, 1.7x$10^{13}$/µL) were used for PFC-HP long projection study. PHP-CAG-FLEX-Rpl22-HA (4.11 x $10^{14}$ vg/mL) was produced by packing plasmid pAAV-CAG-FLEXon-Rpl22-3HA in to PHP.eB. Lenti-GfaABC1D-Rpl22-HA was produced by packing plasmid pZac2.1-GfaABC1D-Rpl22-HA (Addgene, 111811) into Lentivirus.

### Intracranial and retro-orbital virus injections

For intracranial virus injection, mice were anesthetized with 2% isoflurane and placed in a stereotaxic head frame on a heat pad. Artificial tears were applied to the eyes to prevent eye drying. A midline incision was made down the scalp, and a craniotomy was performed with a dental drill. A Nanoliter injector (World Precision Instruments) was used to infuse virus with Micro4 Controller (World Precision Instruments). Virus was infused at 50-100 nL/min. For PFC-HP long projection study, 10-fold dilution of retroAAV-PKG-cre and 3-fold dilution of AAV1-CAG-flex-EGFP were applied 7-days post VaD. The coordinates for 2 injections (each of 0.5µL of virus solution) in HP were: anterior-posterior (AP) -2.0mm, medio-lateral (ML) +1.7mm, dorsoventral (DV) −1.3mm (CA1) and 1.6mm (DG); the coordinates for 1 injection (0.3 µL of virus solution) in medial PFC was: AP +1.70mm, ML +0.45mm, DV −1.25mm. For astrocyte TRAP cell labeling, three injections (each of 0.5µL of Lenti-GfaABC1D-Rpl22-HA virus solution) were made in the following coordinates: AP +1.45mm, ML +2.83mm, DV −1.58mm; AP +0.15mm, ML +2.33mm, DV −1.58mm; and AP +0.15mm, ML +3.33mm, DV −1.60mm. After infusion, the capillary was kept at the injection site for 5 min before slow withdrawn. The incision was closed using VetBond (3M, No.1469SB). The mice were recovery on 37 °C heated blanket, and returned to home cage after woke up. Water with amoxicillin was applied for 1 week.

For retro-orbital virus injection, the protocol was adopted from Yardeni, et al [4]. Briefly, mice were anesthetized with 2% isoflurane through a funnel-shaped nose cone and placed on a 37 °C heated blanket. A 1mL insulin syringe with 27.5-gauge needle was prepared with 50 µL of PHP-CAG-FLEX-Rpl22-HA virus. The needle was placed so the bevel faces down to decrease the likelihood of damaging the eyeball. The needle was inserted follow the edge of the eyeball down until the need tip is at the base of the eye. The virus is injected slowly and smoothly. After the injection is complete, the needle is slowly and smoothly withdrawn. Mice with obvious bleeding or injectate leakage were removed. Artificial tear was applied to protect the eyes after injection. The mice were recovery on 37 °C heated blanket, and returned to home cage after woke up.

**Tamoxifen preparation and administration**

100mg of tamoxifen (Sigma T5648) was dissolved in 4.5mL of prewarmed corn oil (Sigma C8267) at 50°C. The solution was vortexed and warmed repetitively until completely dissolved. The final solution was aliquoted and stored in -20°C until injection day. For Tbx18-creER::Rpl22-HA and Ng2-creER::Rpl22-HA transgenic strains, the mice were injected intraperitoneally at the dosage of 75 mg/Kg/day for 5 consecutive days started from the 7$^{th}$ day before VaD model.

**VaD mouse model**

VaD in the mouse was modified from white matter stroke model [5]. Briefly, the mice were anesthetized with 2% isoflurane and securely mounted onto a stereotaxic apparatus. Core body temperature of the mice was maintained at 36.5 to 37.5°C. A midline incision was made down the scalp, and a craniotomy was performed with a dental drill. A Nanoliter injector (World Precision Instruments) was used to infuse 27μg/μL of L-NIO (Sigma-Aldrich, 400-600) in sterile saline (Hospira) with Micro4 Controller injector (World Precision Instruments), at the speed of 100 nL/min. To avoid damage to motor cortex, the glass pipette (Wrld Precision Instruments, 1B100F-4) containing the L-NIO was inserted through the cortex of the frontal lobe into the underlying subcortical WM at an angle of 36°. Three injections (each of 0.3μL of L-NIO solution) were made in the following coordinates: AP +0.80mm, ML +2.00mm, DV −1.56mm; AP +0.80mm, ML +2.83mm, DV −1.60mm; and AP +0.80mm, ML +3.66mm, DV −1.61mm. Localized vasoconstriction leads to focal ischemia in the subcortical WM [5].

**Brain tissue microdissection and TRAP**

Mice were anesthetized with isoflurane and perfused with 20mL cold PBS to remove the circulating macrophages. Coronal forebrain sections (300μm thick) were collected using mouse coronal section block on cold PBS, and placed to a glass dissection surface under a stereoscope maintained at 4°C. Subcortical WM was microdissected from control and VaD brains using fine tipped forceps. Individual samples were transferred into an RNA low-binding 1.5mL tube and snap-froze on smashed dry ice. All the samples were stored in -80°C before TRAP procedure.

The TRAP protocol was modified from Heiman et al [6]. RNA low-binding tubes with microdissected subcotical WM tissue, one animal per tube, were placed on ice. 300μL of homogenate buffer was added into each tube. The tissue samples were homogenated using grinding pestles, followed by 30 repetitive pipetting using 200 μL tips. 25 μL of the homogenate was saved as input samples. The rest of the homogenate was centrifuged at 4°C and 10K rpm for 10 min. The supernatant was collected to another RNA low-binding tube and incubated with 3 μL of HA antibody (Covance, MMS-101P) with rotation at 4°C for 4 h. Wash the Dynabeads Protein G-magnetic (Thermo Fisher Scientific, 10004D) with homogenate buffer once and then incubated with homogenate-HA antibody mixture at 4°C over-night on rotator. The ribosome-associated mRNAs were bound to the Protein G magnetic beads at this point. Then, these tubes were placed in magnetic rack to let the beads precipitate. The supernatant was removed and the beads were washed with 400 μL/tube of high salt buffer, 3 times at 4°C on rotator. 300 μL of lysis buffer from RNA extraction kit NucleoSpin® RNA XS (Macherey-Nagel) was added into each tube. The tubes were vortexed for 30 seconds and the supernatant (pulldown samples) containing cell type specific ribosome-associated mRNA were stored in -80°C until RNA extraction.

**Subcortical white matter tissue dissociation and FACS**

The microglia FACS protocol was modified from previous report [7]. Briefly, C57BL/6J mice were anesthetized with isoflurane and perfused with 20mL cold PBS to remove the circulating macrophages. Subcortical WM was microdissected from control and VaD brains using fine tipped forceps, and minced using a scalpel under the stereoscope before being transferred Eppendorf tubes containing 1mL of Hibernate A solution (Brain Bits) stored on ice.

Microdissected WM tissues were gently dissociated in Hibernate A solution using sequential trituration with fire-polished glass pipettes with openings of decreasing diameter (final pipette ~0.4 mm diameter opening). Resulting cell suspensions were spun down, resuspended in 300μL 1x PBS and filtered through a 40μm mesh filter. Cells were then washed once with 1x PBS, resuspended in 300μL 1x PBS and filtered as above. Filtered cells were then incubated for 20 min on ice with the following antibodies: APC conjugated Rat anti-CD11b (1:100, BD Pharmingen), PE-Cy7 conjugated Rat anti-CD45 (1:400, BD Pharmingen), Brilliant Violet – 421 conjugated mouse anti-CX3CR1 (1:200, BioLegend). Throughout the experiment, samples were kept at 4°C on ice.

Samples were sorted using a FACS Aria cell sorter (BD Biosciences). The population of cells containing microglia could be readily identified based on forward scattering (FSC) and side scattering (SSC) properties. A gating strategy based on FSC and SSC width and height was used to select only single cells.

**RNA extraction and sequencing**

For EC, pericyte, OPC, and astrocyte, total RNA was isolated using the NucleoSpin® RNA XS (Macherey-Nagel), library preparation (Takara SMART-Seq v4 Ultra low Input RNA kit) and sequencing [NovaSeq (PE100/150)] were conducted by MedGenome. Input and pulldown samples were extracted parallelly on the same day. Samples with high RNA integrity number (>6) were used for library construction. More detailed information on the sequencing depth, RIN value and unique alignment rate can be found in Supplementary Fig 9. Microglia samples for RIN value test were separate from sequenced samples, with RIN value >6.0. Microglia total RNA was isolated using PicoPure$^{TM}$ kit (Thermo Fisher, KIT0204), library preparation [Nugen Ovation RNA Ultra Low Input (500 pg) + Kapa Hyper] and sequencing (PE 2x75) were conducted by the UCLA Neurosciences Genomics Core (UNGC). The uniquely mapped reads (%) for Control_P2, VaD_P2, and VaD_P4 are 68%-78%; The number of reads per sample are 32-56M. All RNA extractions were stored in RNase-free tubes at −80°C until further processing.

**Reverse-transcript PCR, pre-amplification, and quantitative real-time PCR**

Reverse transcription was performed using PrimeScript™ RT Reagent Kit (Takara, RR037A). Pre-amplification is performed using TaqMan$^{TM}$ PreAmp Master Mix (4391128). Quantitative real-time polymerase chain reaction (qRT-PCR) were performed using Premix Ex Taq$^{TM}$ (Takara, RR390A). All the performance was conducted following the vendor provided protocols. The PCR probes used in this study are listed below.

| Gene name | Taqman probe Cat# |
|---|---|
| Aif1 | Mm00479862_g1 |
| Gapdh | Mm99999915_g1 |
| Gfap | Mm01253033_m1 |
| Pdgfrα | Mm00440701_m1 |
| Pdgfrβ | Mm00435553_m1 |
| Tek | Mm00443243_m1 |

**Bioinformatics**

To identify DEGs, Hiset2 or STAR was used to align reads to the GRCm38 genome assembly. An FDR value threshold of 0.1 was imposed for the likelihood ratio test to select DEGs, using edgeR, limma, and voom packages. Expression level estimation was reported as fragments per kilobase of transcript Fragments Per Kilobase Million (FPKM) value (www.WhiteMatterWiki.org).

To construct custom-made L-R database, the lists of L-R interactions were downloaded from 3 major databases [8-10]. R package tidyr was used to merge the databases and remove the duplicates. The source database and species (human or mouse) were annotated. R package circlize was used to draw Circos plots.

To identification of WM specific transcriptional factors (TFs) and cell type markers, the mouse TFs were downloaded from KEGG (https://www.kegg.jp/brite/mmu03000), and the canonical cell type markers were adopted from Barres' database [11]. A stringent approach was employed to identify WM and cell type-specific TFs/markers based on, 1) FPKM at least two-fold higher than in other cell types, 2) enrichment fold change >1 (calculated as the fold-change of FPKM, pulldown vs input), and 3) exclusion of genes with an FPKM <10. The calculation of normalized specificity in WM, cortex, or the whole brain involves dividing the FPKM value in the target cell by the average FPKM value across all cell types (including the cell type being considered) and then further dividing this result by the total number of cell types. This computation yields a value within the range of 0 to 1, where 0 signifies no expression, and 1 indicates that the gene is exclusively expressed in the target cell type.

To identify ECM and GPCR genes in different cell types, gene ontology annotations of ECM and GPCR were downloaded from MGI (ECM: https://www.informatics.jax.org/go/term/GO:0031012; GPCR: https://www.informatics.jax.org/go/term/GO:0004930). Bubble plots were drawn based on the cell-type specific expression of ECM and GPCR genes, and their changes in VaD.

The Venn diagrams in Figs 2 and 3 were draw using R package ggvenn, or http://bioinformatics.psb.ugent.be/webtools/Venn/, or Venn.drawio.

Rank-rank hypergeometric overlap (RRHO) [12] analyses were performed using average FPKM from control samples of the WM dataset, compared with Barres' dataset [11], which was filtered to include only the genes with an FPKM greater than 1 for each cell type. These were also compared with Betsholtz's dataset [13], which was generated by bulk-normalizing counts to produce FPKM values.

Gene set enrichment analysis (GSEA) was done using GSEA 4.3.2 [14, 15] with MsigDB ver7.0.

KEGG and GO pathway analyses were done by ShinyGO0.77.

***In situ* hybridization**

Mouse brains were dissected and fast-frozen in OCT by dry ice without PFA fixation. 15 μm frozen sections were sliced using cryostat. *In situ* hybridization was performed using RNAscope Fluorescent Multiplex Reagent Kit V2 (ACD, 323110) according to the manufacturer's instructions. RNAscope Probe-Mm-Adora3-O1 (ACD, 461891) was used to detect Adora3 mRNA. Probe-Mm-Aif1-C2 (ACD, 319141-C2) was used as marker for microglia. Probe-Mm-Anpep-C3 (ACD, 417181-C3) was used as marker for microglia. Probe-Mm-Serpine2-C3 (ACD, 435241-C3) was used as marker for Serpine2 mRNA.

**Immunostaining using mouse brain sections and confocal images**

Animals were perfused transcardially with 0.1 M phosphate-buffered saline (PBS), followed by 4% paraformaldehyde (PFA). The brains were removed, post-fixed overnight in 4% PFA, and sectioned into consecutive 50-μm-thick slices using vibratome (Leica). Immunostaining was performed by brain section blocking in 5% normal donkey serum for 1 hour at room temperature, incubating in primary antibody overnight at 4°C, and in secondary antibody for 1 hour at room temperature. For the performance using CD39 and Pdgfrβ antibodies, antigen retrieval using sodium citrate buffer (10 nM, pH 6.0), in pressure cooker for 3 min was applied before blocking. All sections were counterstained with 4′,6-diamidino-2-phenylindole (DAPI). The antibodies used in this study are listed below.

| **Antibody** | **Vendor / Species / Cat #** | **Concentration** |
|---|---|---|
| A3AR | Abcam / Rabbit / ab197350 | 1:300 |
| Aspa | EMD Millipore / Rabbit / ABN1698 | 1:500 |
| CD13 | Abcam / Rat / ab33489 | 1:300 |
| CD31 | R&D / Rat / AF3628-SP | 1:300 |
| CD31 | Invitrogen / Rabbit / PA5-16301 | 1:300 |
| CD39 | Abcam / Rabbit / ab223842 | 1:500 |
| c-Fos | Cell Signaling / Rabbit / 2250 | 1:500 |
| Cre-Recombinase | Synaptic Systems GmbH / Guinea pig / 257004 | 1:500 |
| Ctip2 | Abcam / Rat / ab18465 | 1:500 |
| Cux1 | Proteintech / Rabbit / 11733-1-AP | 1:500 |
| Gfap | Invitrogen / rat / 13-0300 | 1:2000 |
| Glut1 | Millipore Sigma / Rabbit / 07-1401 | 1:500 |
| HA | Roche / Rat / 11867431001 | 1:300 |
| Iba1 | Wako / Rabbit / 019-19741 | 1:1000 |
| Iba1 | Abcam / Goat / Ab5076 | 1:500 |
| Lrp1 | Thermo Fisher Scientific / Rabbit / BS-2677R | 1:100 |
| MBP | Abcam / Rabbit / Ab40390-1001 | 1:500 |
| NeuN | Chemicon / Mouse / MAB377 | 1:1000 |
| NeuN | Synaptic Systems GmbH / Guinea pig / 266004 | 1:500 |
| NF160 | Abcam / Mouse / Ab7794-1001 | 1:500 |
| Olig2 | EMD Millipore / Rabbit / AB9610 | 1:500 |
| Pdgfrα | R&D / Goat / AF1062 | 1:500 |
| Pdgfrβ | R&D / Goat / AF1042 | 1:500 |
| Satb2 | Abcam / Rabbit / ab92446 | 1:1000 |
| Anti-Goat-IgG | Jackson ImmunoResearch / Donkey | 1:1000 |
| Anti-Guinea Pig-IgG | Jackson ImmunoResearch / Donkey | 1:1000 |
| Anti-Mouse-IgG | Jackson ImmunoResearch / Donkey | 1:1000 |
| Anti-Rabbit-IgG | Jackson ImmunoResearch / Donkey | 1:1000 |
| Anti-Rat-IgG | Jackson ImmunoResearch / Donkey | 1:1000 |

High-resolution confocal images in z-stacks were acquired (Nikon C2). Area measurements of the infarct core, WM axonal projections stained with NF160 and MBP were quantified with Imaris (Bitplane, version 10.0.0). The parameters for scanning were kept constant across treatment, and conditions.

**Immunohistochemistry (IHC) using human brain sections and whole slide imaging**

IHC protocol was modified from previous reports [16, 17]. Briefly, formalin-fixed periventricular white matter blocks of normal and VaD patients were obtained from the NIH brain bank. Samples were paraffin embedded, sectioned at 7 µm in thickness, immunostained with CD39, Glut1, or Iba1 primary antibodies followed by either horse anti-mouse or horse anti-rabbit secondary antibody conjugated to horseradish peroxidase (HRP), visualized with DAB as chromogen (Vector Laboratories, S-2012), and counterstained with hematoxylin. Slides were then scanned and digitized using the Pannoramic Midi 2 (Epredia). The intensity of immunoreactivity was analyzed using the Positive Pixel Count algorithm in the ImageScope program. The antibodies used in this study are listed below.

| **Antibody** | **Vendor / Species / Cat #** | **Concentration** |
|---|---|---|
| CD39 | Abcam / Rabbit / ab223842 | 1:500 |
| Glut1 | Millipore Sigma / Rabbit / 07-1401 | 1:500 |

| | | |
|---|---|---|
| Iba1 | Invitrogen / Mouse / MA5-27726 | 1:300 |
| Anti-Mouse-IgG-HRP | Vector Laboratories / Horse / MP-7402-15 | N/A |
| Anti-Rabbit-IgG-HRP | Vector Laboratories / Horse / MP-7401-50 | N/A |

Patient information is listed below.

| NIH brain bank sample # | Sex | Age | Diagnose |
|---|---|---|---|
| 21278 | Female | 84 | VaD/LOAD* |
| 298992 | Female | 86 | No dementia |
| 33987 | Male | 63 | No dementia |
| 17629 | Female | 81 | VaD |
| 17678 | Male | 89 | VaD |
| 53706 | Female | 79 | VaD |
| 36866 | Female | 79 | No dementia |
| 34913 | Male | 85 | No dementia |
| 236440/81764 | Female | 88 | VaD/LOAD* |
| 46146 | Male | 71 | No dementia |
| 9700 | Male | 72 | VaD |
| 73095 | Male | 81 | VaD |
| 54549 | Male | 86 | VaD |
| 63781 | Male | 63 | VaD |
| 236428/56356 | Female | 79 | No dementia |
| 86213 | Male | 89+ | No dementia |
| 11371 | Male | 89+ | No dementia |
| 1176 | Female | 85 | No dementia |
| 17230 | Female | 89+ | No dementia |
| 90996 | Female | 89+ | VaD |
| 51570 | Male | 89+ | VaD |
| *LOAD: late onset Alzheimer's disease. LOAD was diagnosed in addition to VaD after samples were request by authors. | | | |

**Osmotic pump and brain infusion kit implantation**

Osmotic mini-pumps (1002, 1004) were subcutaneously implanted and connected to right lateral ventricle through brain infusion kit 3 (Alzet). The brain infusion kits were fixed onto the skull with super glue (Loctite, 45198). For 4-week treatment, 1004 mini-pumps were used to delivery 400 μM of CF101 (in 1% DMSO in saline) at the speed of 0.11 μL/h; For 2-week treatment, 1002 mini-pumps were used to deliver 200 μM of CF101 at the speed of 0.23 μL/h.

**Behavior tests**

C57 mice with or without VaD were tested in Fear Conditioning (FC), Memory association, Novel Object Recognition (NOR), and Grid Walking tasks. All tests were performed 3-4 weeks after VaD. For FC, testing mice were first handling for 3 days (1 min/day) and then habituated to transportation and external environmental cues for 2 min in the experimental room each day for another 3 days. During FC test, mice explored the context for 2 min and then shocked for 2 s (0.65 mA). 58-s after the shock, mice were placed back in their home cage. 1 day later, the mice were returned to the same context for 5 min.

For NOR, the testing mice were allowed to explore an open field arena (30 cm x 30 cm) with 2 identical objects for 12 min on the first day. Then the one object was replaced to a novel object and the mice were allowed to explore the 2 different objects for 8 min. The exploration was recorded and the time to explore each object was analyzed by ANY-maze. The data were presented as exploration ratio of time spent exploring the novel object versus both objects. For Grid walking test, the mice were tested 1-day pre-, 4-

day and 21-day post-VaD. Behavior tests were scored by observers who were masked to the treatment group of the animals.

Memory linking is a process by which new information is linked to previously stored information in the brain. This linking process helps to store information in a structured way and makes it easier to retrieve later. By forming connections between new and old information, memory association enhances the strength of memory traces and helps to organize the information in a meaningful way, leading to more effective memory storage and recall. Compared to single memory, memory linking is more sensitive to changes in the microenvironment (e.g. aging) [18, 19]

For memory linking task, mice explored 2 different contexts (A and then B, counterbalanced) which were separated by 5h-7d. Mice explored each context for 10 min. For immediate shock, mice were placed in chamber B for 10s followed by a 2-s shock (0.65 mA). 58 seconds after the shock, mice were placed back in their home cage. For the context tests, mice were returned to the designated context (A, B and a novel context C, counterbalanced). The freezing was assessed via an automated scoring system (Med Associates) with 30 frames per second sampling; the mice needed to freeze continuously for at least one second before freezing could be counted.

**Statistical analysis and reproducibility**

The investigators who collected and analyzed the data including behavior, bioinformatic analysis, mouse brain immunostaining, human brain IHC, were blinded to the VaD and treatment conditions. Mice were randomly allocated to treatment condition using a randomized block experimental design (restricted randomization). All data are demonstrated in figures as means ± SEM. All statistical analyses were performed using GraphPad Prism 8. For behavior experiments, RNA-Seq, quantification of immunostaining and human IHC, n designates the number of mice or patients. Statistical significance was assessed by Student's t test, or one- or two-way ANOVA where appropriate, followed by the indicated post hoc tests for repeated measures. Significance levels were set to $P = 0.05$. Significance for comparisons: *$P < 0.05$; **$P < 0.01$; ***$P < 0.001$.
