## Supplementary figures and images for "Intercellular Signaling Pathways as Therapeutic Targets for Vascular Dementia Repair"

### Supplementary Figure 1

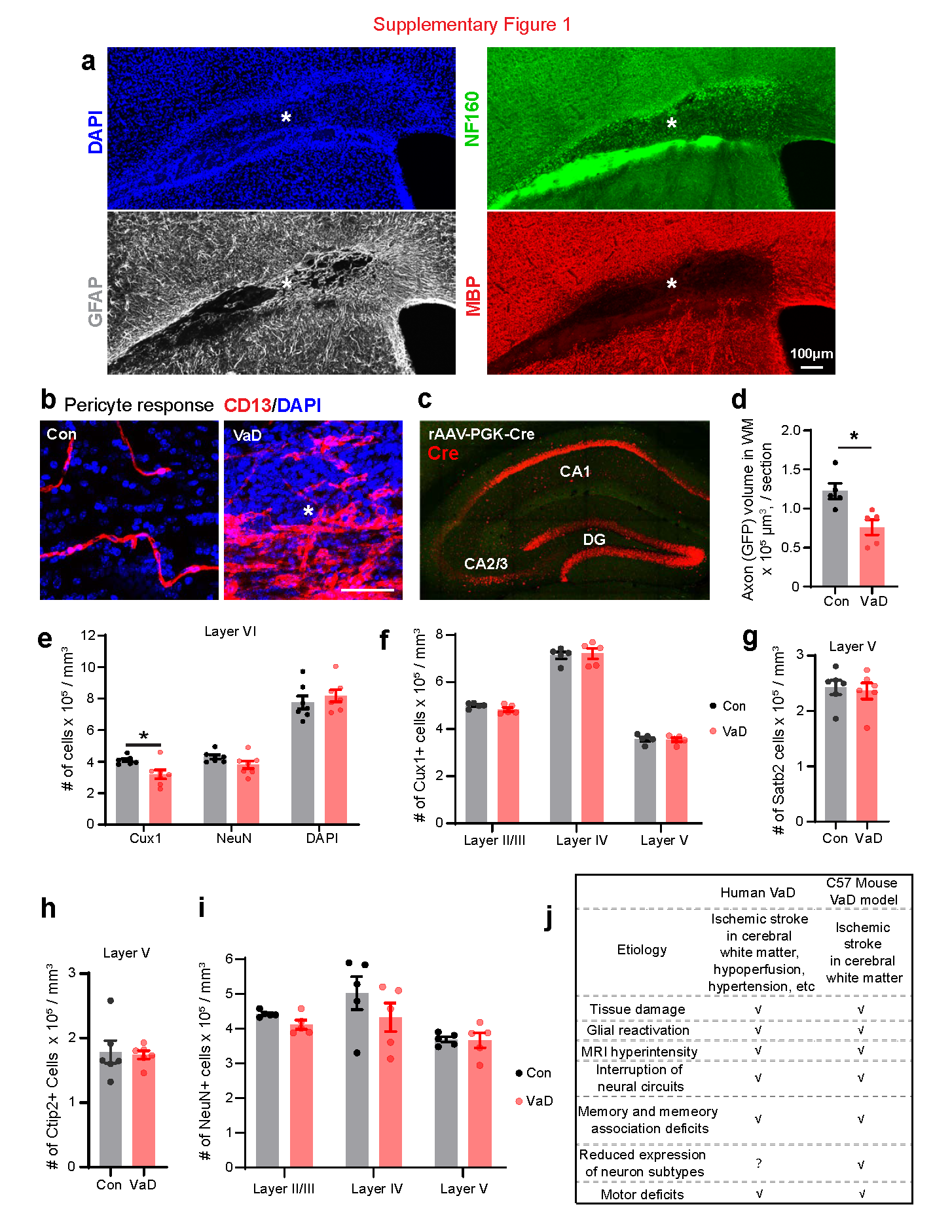

### Supplementary Figure 2

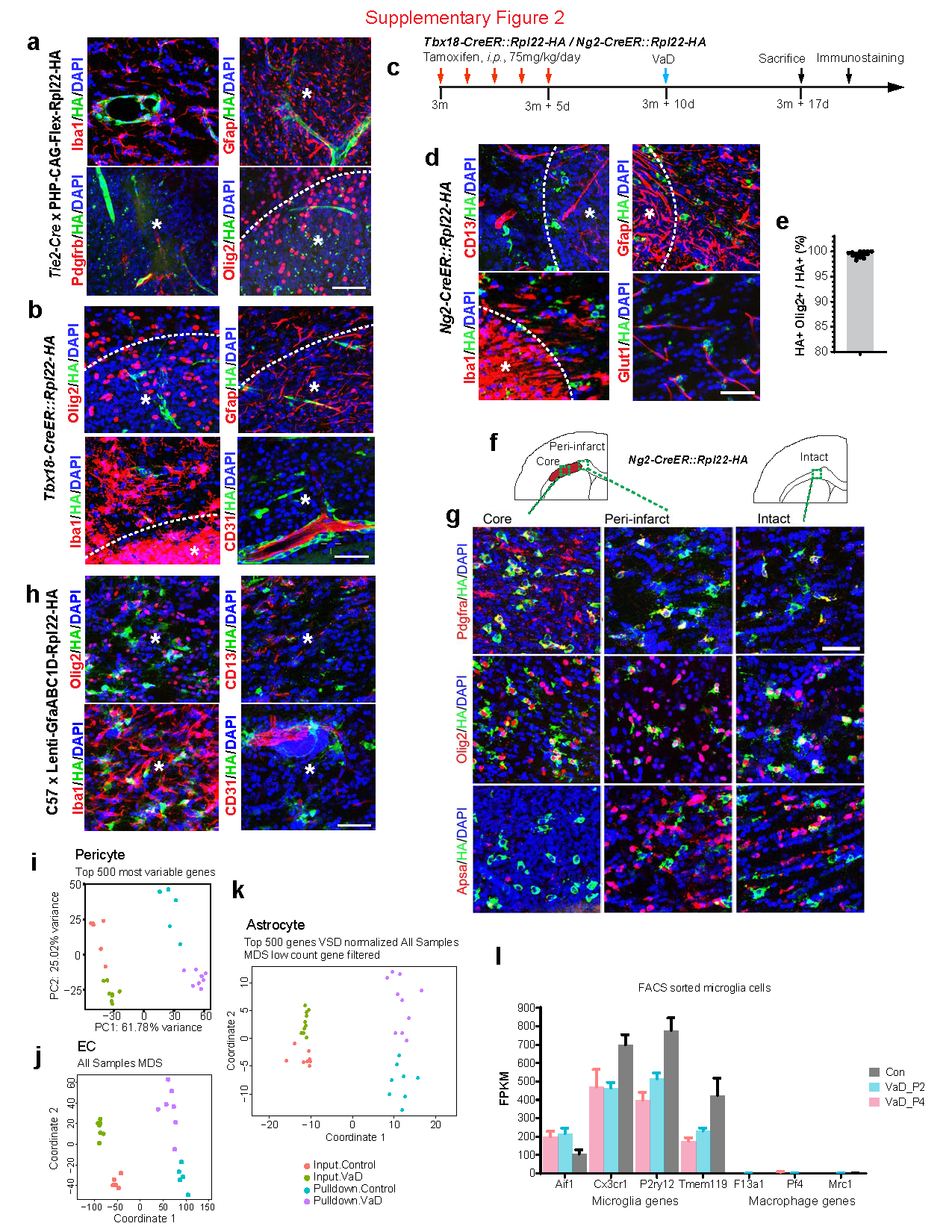

### Supplementary Figure 6

#### Supplementary Figure 6

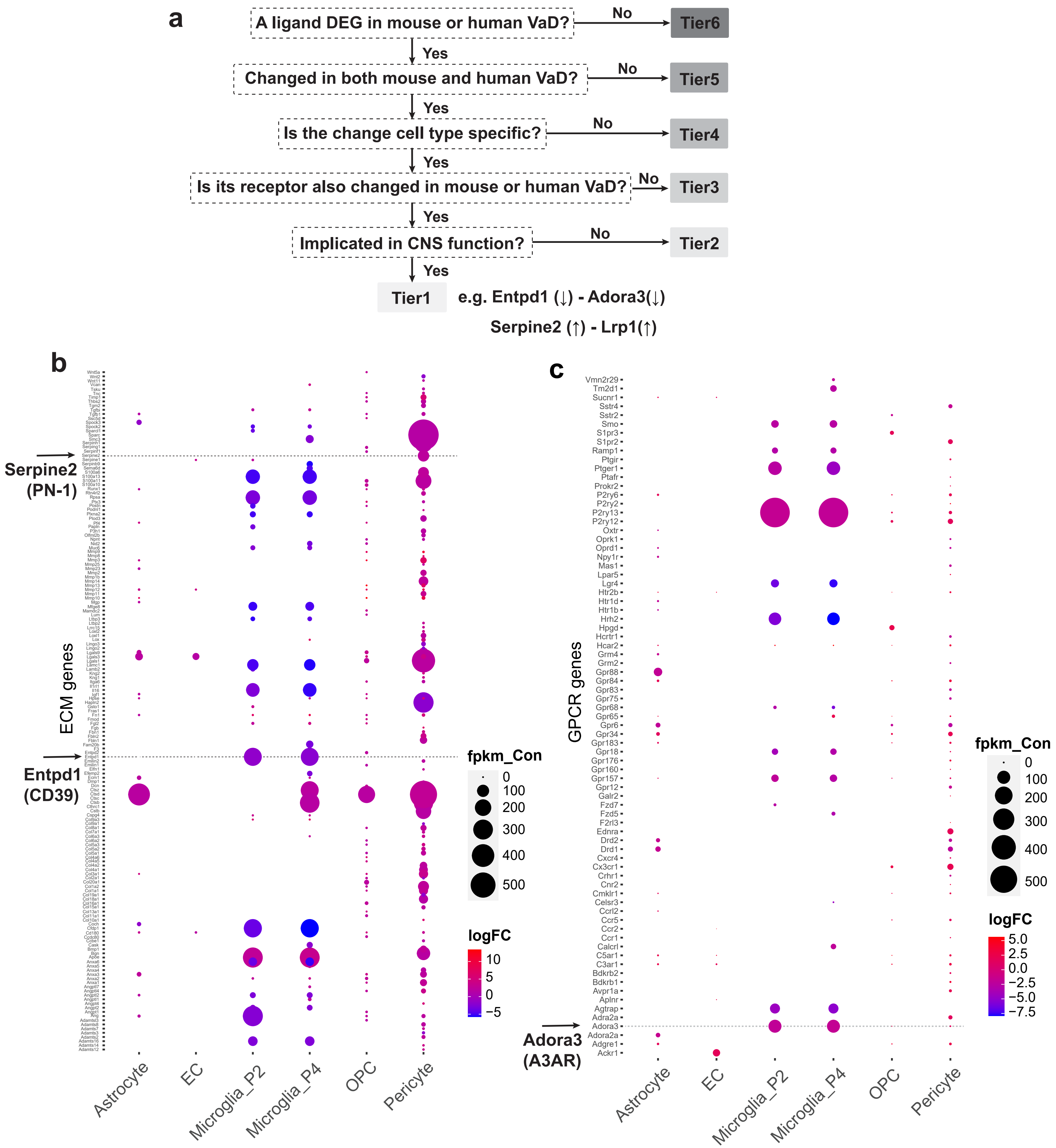

### Supplementary Figure 7

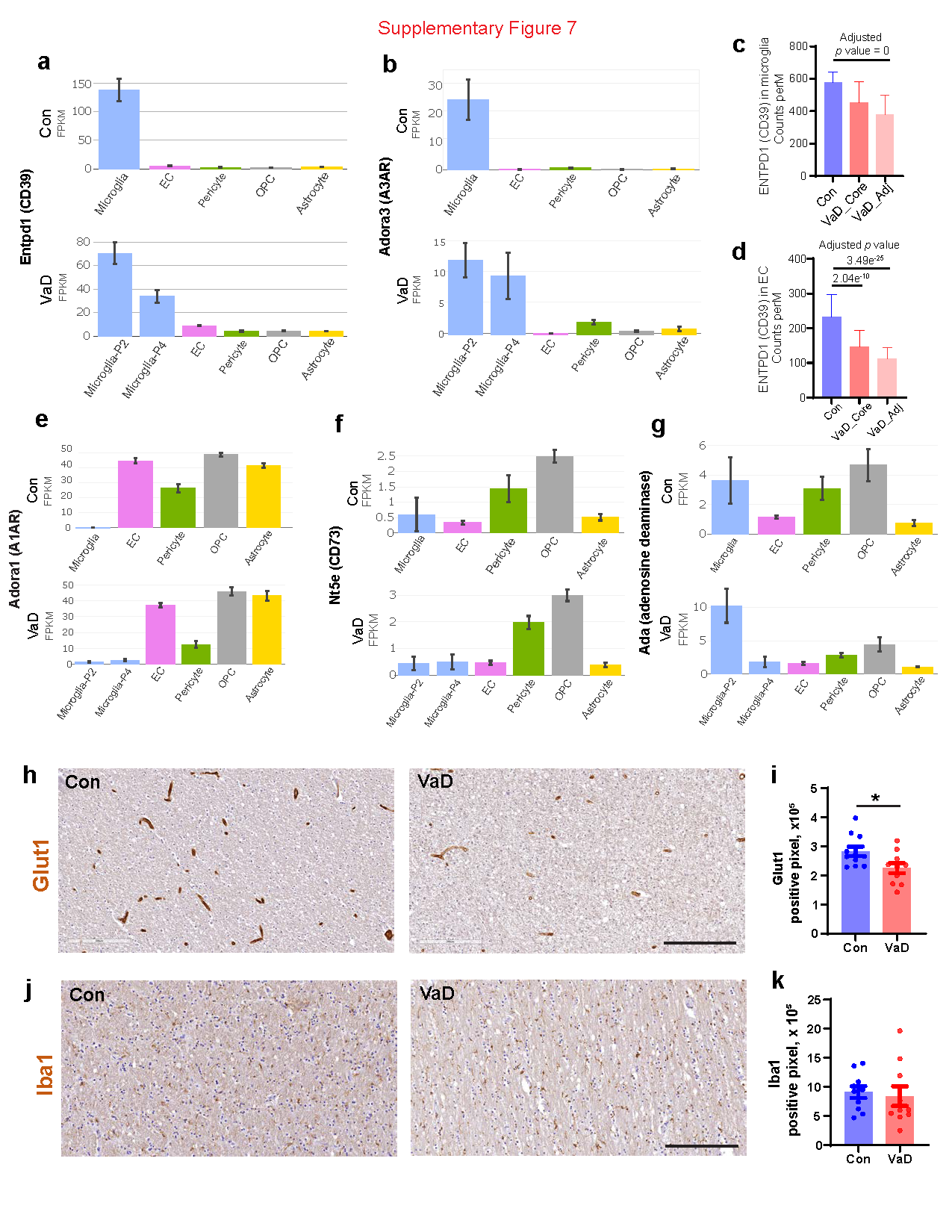

### Supplementary Figure 8

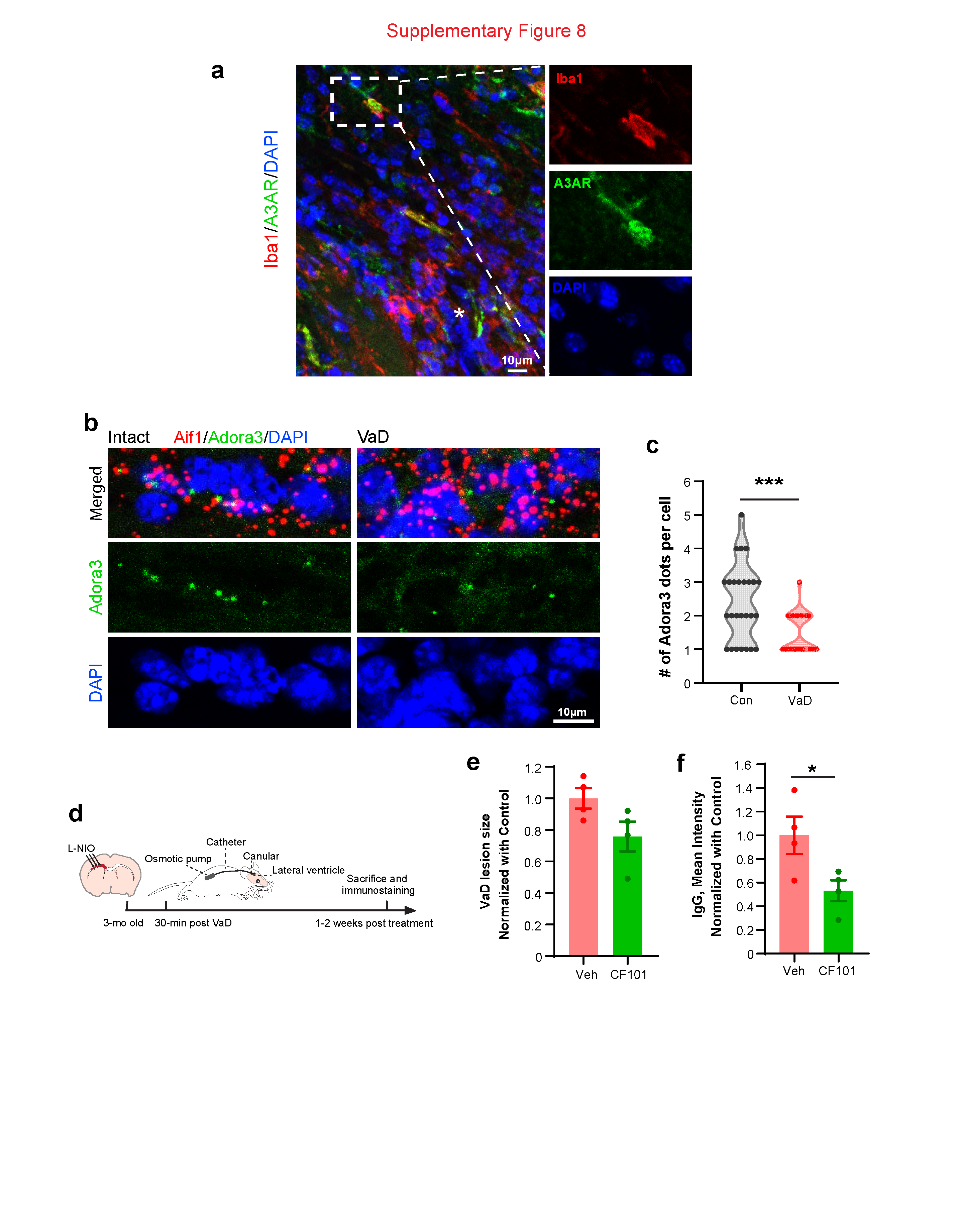
