## Supplementary Figure 3 for "Intercellular Signaling Pathways as Therapeutic Targets for Vascular Dementia Repair"

### a White matter specific transcriptional factor

| Cell type | TF | Normalized specificity in WM (0-1) | Normalized specificity in cortex / whole brain (0-1) |
|---|---|---|---|
| EC | Klf2 | 0.83 | 0.29 |
| EC | Npas1 | 0.74 | 0.00 |
| EC | Lhx6 | 0.73 | 0.05 |
| Peri | Irf3 | 0.42 | 0.09 |
| Peri | Hes1 | 0.54 | 0.10 |
| Peri | E4f1 | 0.49 | 0.06 |
| Peri | Zfp692 | 0.60 | 0.05 |
| Peri | Foxc1 | 0.88 | 0.11 |
| Peri | Crebzf | 0.53 | 0.06 |
| Peri | Zfp14 | 0.53 | 0.05 |
| Peri | Tbx3 | 0.83 | 0.08 |
| Peri | Foxf2 | 0.69 | 0.04 |
| Peri | Ebf1 | 0.75 | 0.10 |
| OPC | Mycl | 0.89 | NA |
| OPC | Sox4 | 0.73 | 0.12 |
| OPC | Tox3 | 0.55 | 0.40 |
| OPC | Hes5 | 0.53 | 0.19 |
| Astro | Atoh8 | 0.62 | 0.25 |
| Astro | Hopx | 0.55 | 0.23 |
| Micro | Mef2a | 0.89 | 0.16 |
| Micro | Rfx7 | 0.77 | 0.02 |
| Micro | Zfp292 | 0.71 | 0.02 |
| Micro | E2f3 | 0.75 | 0.13 |
| Micro | Prdm2 | 0.71 | 0.12 |
| Micro | Zfp871 | 0.70 | 0.04 |
| Micro | Zfp644 | 0.67 | 0.03 |
| Micro | Zfp142 | 0.67 | 0.09 |
| Micro | Elk4 | 0.65 | 0.09 |
| Micro | Foxj2 | 0.62 | 0.23 |
| Micro | Sp4 | 0.65 | 0.03 |
| Micro | Zfp788 | 0.54 | 0.01 |
| Micro | Clock | 0.57 | 0.04 |
| Micro | Zbtb44 | 0.54 | 0.07 |
| Micro | Zfp654 | 0.58 | 0.03 |
| Micro | Crebrf | 0.47 | 0.01 |
| Micro | Creb1 | 0.55 | 0.07 |
| Micro | Aebp2 | 0.46 | 0.23 |
| Micro | Arid1b | 0.50 | 0.10 |
| Micro | Prdm4 | 0.44 | 0.07 |
| Micro | Foxk2 | 0.46 | 0.11 |
| Micro | Zfp532 | 0.40 | 0.05 |

**b**

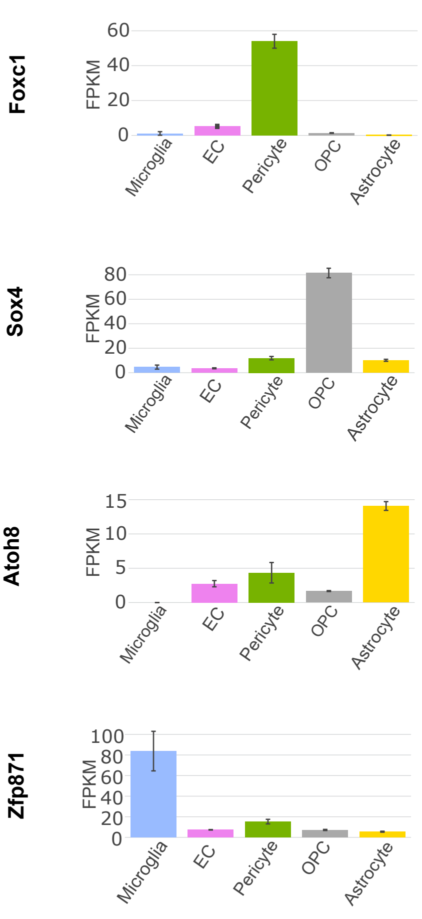


**c**

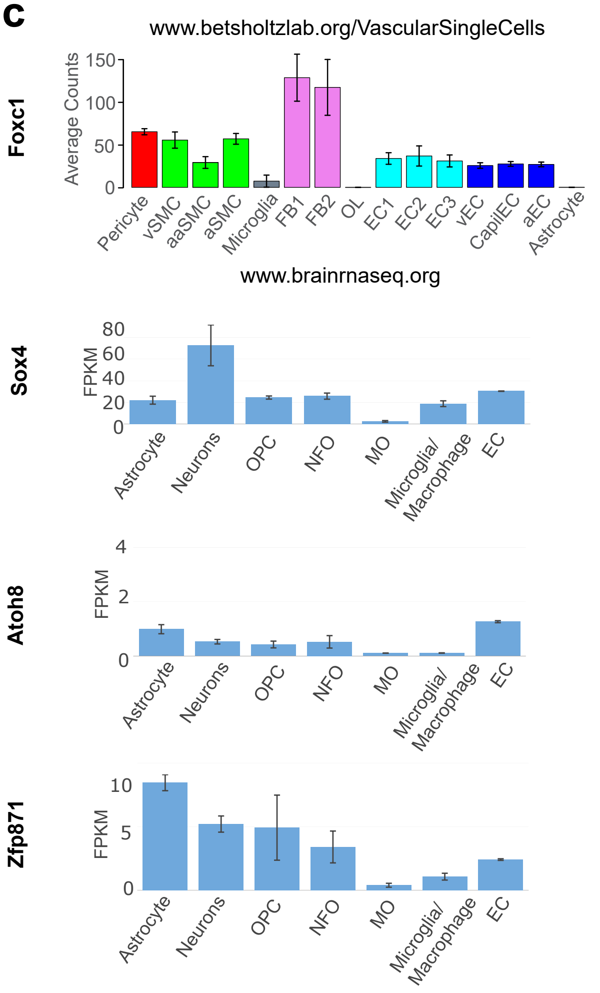


### d White matter specific cell marker

| Cell type | Marker | Normalized specificity in WM (0-1) | Normalized specificity in cortex / whole brain (0-1) |
|---|---|---|---|
| EC | Tmem212 | 0.71 | 0.30 |
| Peri | Aldh1a2 | 1.00 | 0.16 |
| Peri | Col1a1 | 0.98 | 0.16 |
| Peri | Col3a1 | 1.00 | 0.16 |
| Peri | Dcn | 1.00 | 0.17 |
| Peri | Dpep1 | 1.00 | 0.17 |
| Peri | Fam180a | 0.99 | 0.17 |
| Peri | Igf2 | 0.98 | 0.16 |
| Peri | Igfbp2 | 0.92 | 0.15 |
| Peri | Inmt | 1.00 | 0.17 |
| Peri | Lum | 1.00 | 0.17 |
| Peri | Mfap4 | 0.99 | 0.17 |
| Peri | Rgs4 | 0.76 | 0.16 |
| Peri | Slc22a6 | 1.00 | 0.15 |
| Peri | Tcf21 | 1.00 | 0.17 |
| Peri | Tgfbi | 0.95 | 0.16 |
| Peri | Tpm2 | 0.85 | 0.16 |
| OPC | Slc1a1 | 0.81 | 0.25 |
| Astro | Scara3 | 0.88 | 0.65 |
| Astro | Sdc4 | 0.77 | 0.32 |
| Astro | Thbs4 | 0.91 | 0.58 |
| Micro | Olfr172 | 1.00 | 0.14 |
| Micro | Pvrig-ps | 1.00 | NA |
| Micro | Ppnr | 1.00 | NA |
| Micro | H60b | 0.97 | 0.44 |
| Micro | Ighd | 0.97 | NA |
| Micro | Upk1b | 0.97 | 0.21 |
| Micro | Rab39 | 0.94 | 0.44 |
| Micro | Srgap2 | 0.93 | 0.39 |
| Micro | Dennd4a | 0.90 | 0.28 |
| Micro | Mef2a | 0.89 | 0.16 |
| Micro | Nav3 | 0.88 | 0.23 |
| Micro | Mgat4a | 0.91 | 0.38 |
| Micro | Bmp2k | 0.90 | 0.32 |
| Micro | Cttnbp2nl | 0.87 | 0.19 |
| Micro | Rn7sk | 0.93 | NA |
| Micro | Slc8a1 | 0.87 | 0.21 |
| Micro | Slfn8 | 0.89 | 0.55 |
| Micro | Tmem131 | 0.77 | 0.09 |
| Micro | Slc12a6 | 0.71 | 0.19 |

**e**

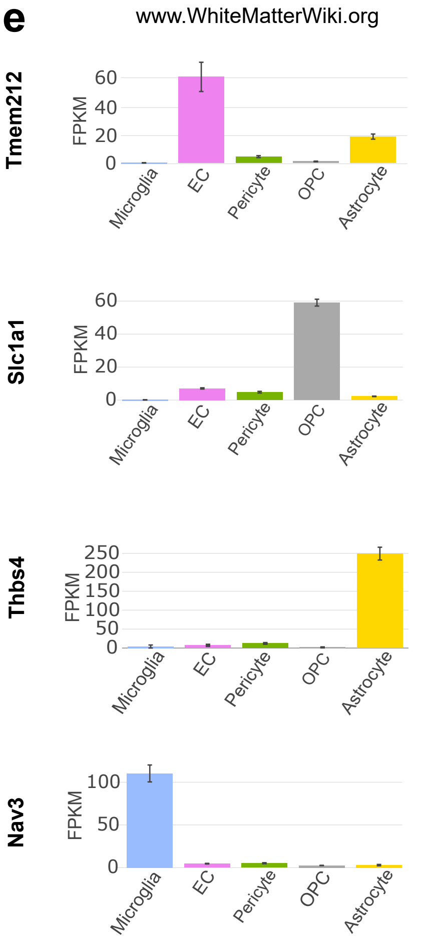


**f**

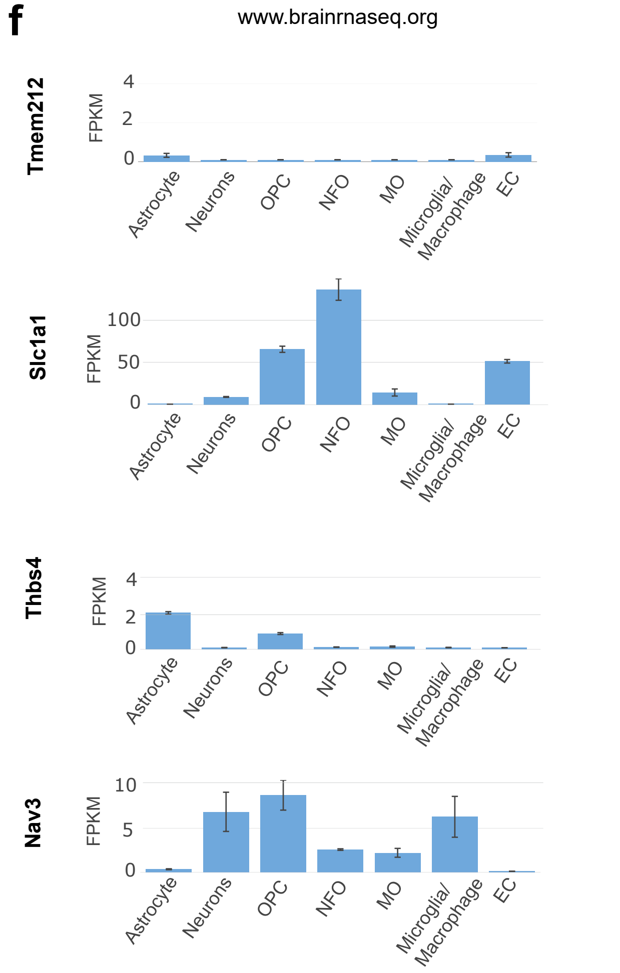
