## Supplementary Figure 4 for "Intercellular Signaling Pathways as Therapeutic Targets for Vascular Dementia Repair"

**a** Pearson Correlation coefficient

Medium correlation: 0.3 - 0.49

Small correlation: < 0.29

| | Cortex | | | | | | |
|---|---|---|---|---|---|---|---|
| **WM** | Micro | EC | Astro | Neuron | OPC | NFO | MO |
| Astro | 0.520 | 0.117 | 0.630 | 0.231 | 0.576 | 0.051 | 0.030 |
| Peri | 0.219 | 0.331 | 0.290 | 0.253 | 0.370 | 0.304 | 0.181 |
| OPC | 0.074 | 0.226 | 0.160 | 0.315 | 0.434 | 0.689 | 0.520 |
| Micro | 0.763 | 0.185 | 0.423 | 0.155 | 0.468 | 0.022 | 0.008 |
| EC | 0.167 | 0.385 | 0.234 | 0.480 | 0.400 | 0.153 | 0.104 |

**b**

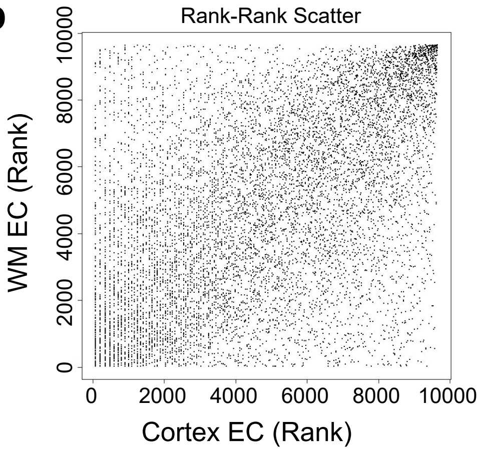


**c**

| | Whole brain | | | | | | | | | | | | | | |
|---|---|---|---|---|---|---|---|---|---|---|---|---|---|---|---|
| **WM** | Micro | aEC | capilEC | vEC | EC1 | EC2 | EC3 | Astro | OL | Peri | aaSMC | aSMC | vSMC | FB1 | FB2 |
| Astro | 0.590 | 0.099 | 0.116 | 0.089 | 0.133 | 0.093 | 0.104 | 0.842 | 0.110 | 0.121 | 0.180 | 0.153 | 0.160 | 0.114 | 0.171 |
| Peri | 0.291 | 0.176 | 0.162 | 0.138 | 0.208 | 0.158 | 0.151 | 0.344 | 0.642 | 0.557 | 0.408 | 0.277 | 0.568 | 0.413 | 0.301 |
| OPC | 0.109 | 0.114 | 0.081 | 0.094 | 0.099 | 0.099 | 0.083 | 0.152 | 0.649 | 0.106 | 0.136 | 0.116 | 0.146 | 0.101 | 0.079 |
| Micro | 0.449 | 0.178 | 0.168 | 0.167 | 0.183 | 0.181 | 0.171 | 0.350 | 0.050 | 0.290 | 0.205 | 0.139 | 0.279 | 0.093 | 0.227 |
| EC | 0.239 | 0.296 | 0.300 | 0.263 | 0.351 | 0.284 | 0.285 | 0.260 | 0.260 | 0.135 | 0.226 | 0.177 | 0.193 | 0.128 | 0.165 |

**d**

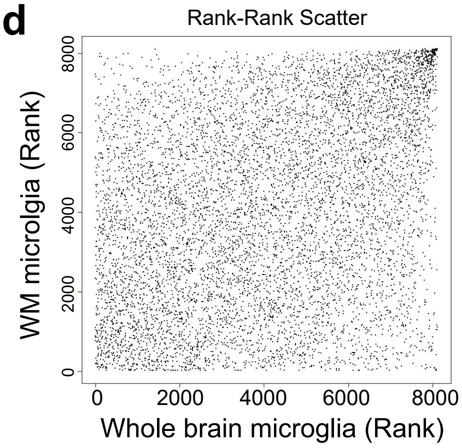


**e**

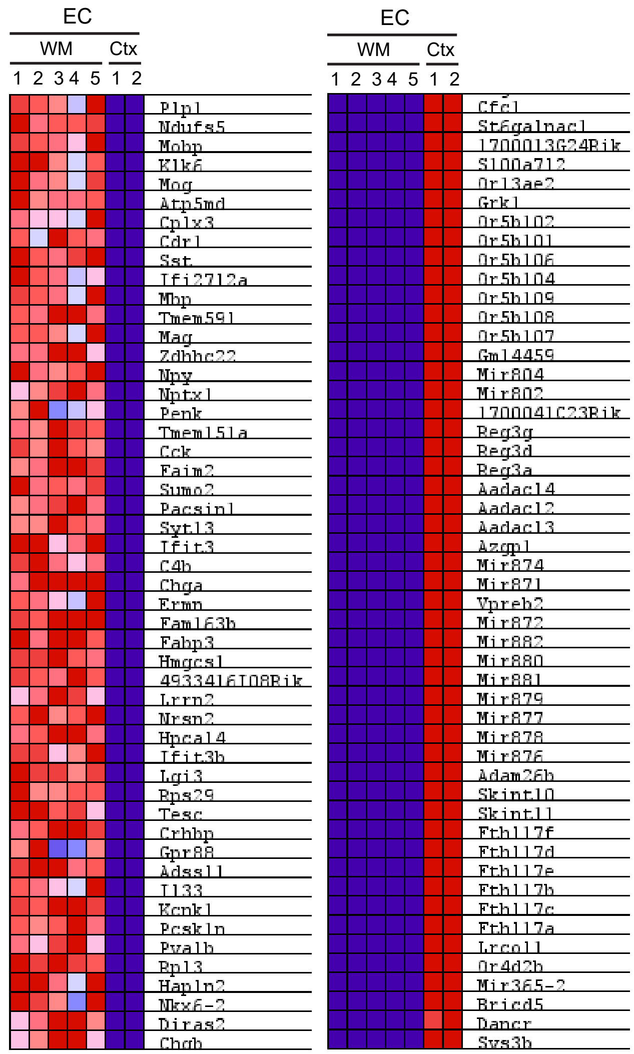


**g**

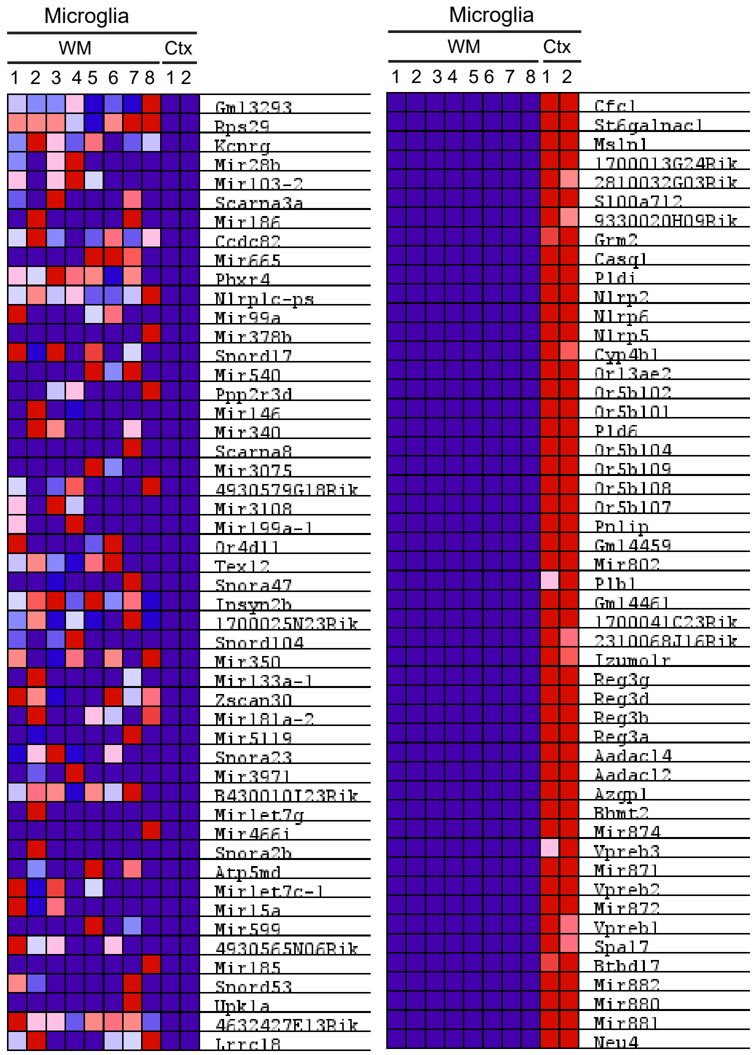


**f**

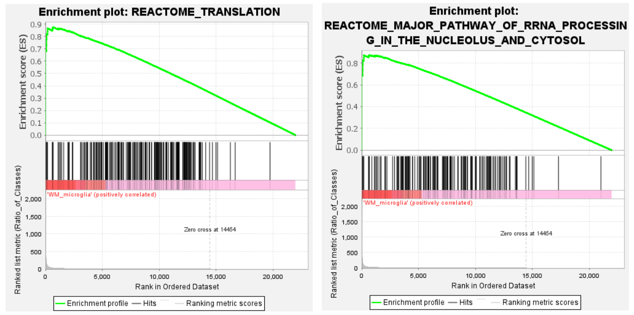


**h**

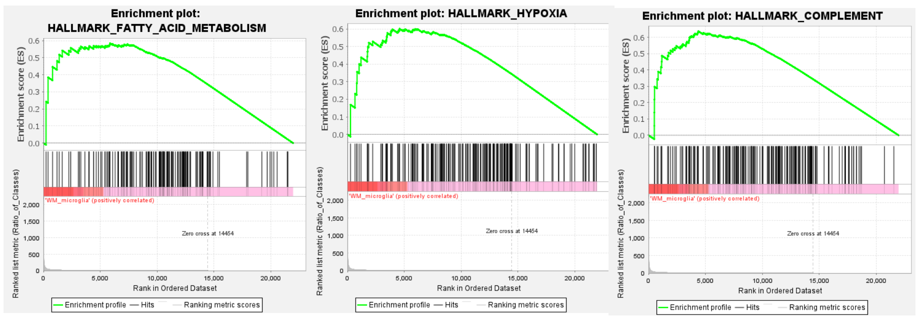
