## Supplementary Figure 5 for "Intercellular Signaling Pathways as Therapeutic Targets for Vascular Dementia Repair"

**a**

Microglia to other cells in mouse VaD (identified by mouse L-R pool)

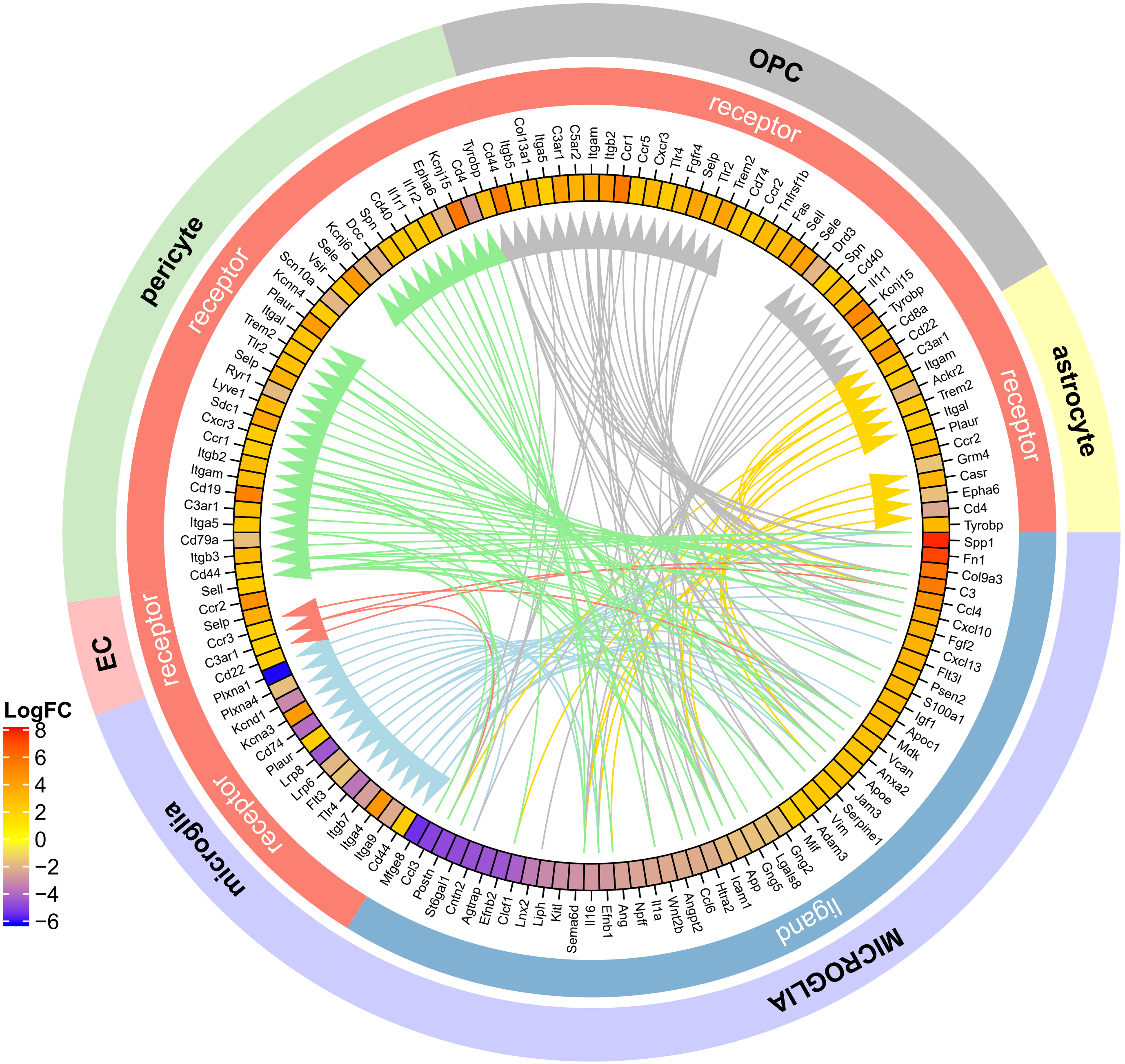

**b**

Microglia to other cells in human VaD (identified by mouse L-R pool)

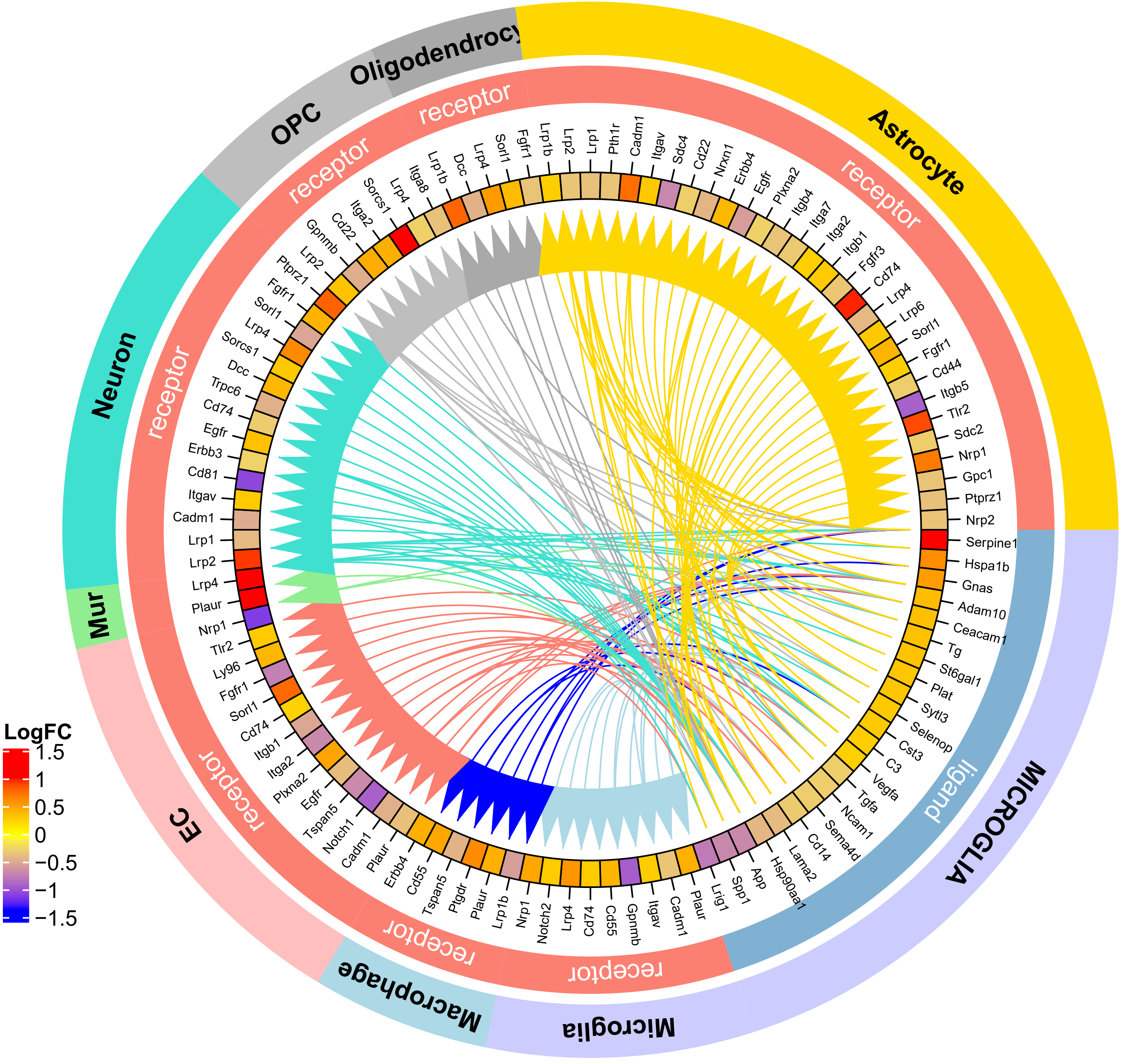

**c**

EC to other cells in mouse VaD (identified by human L-R pool)

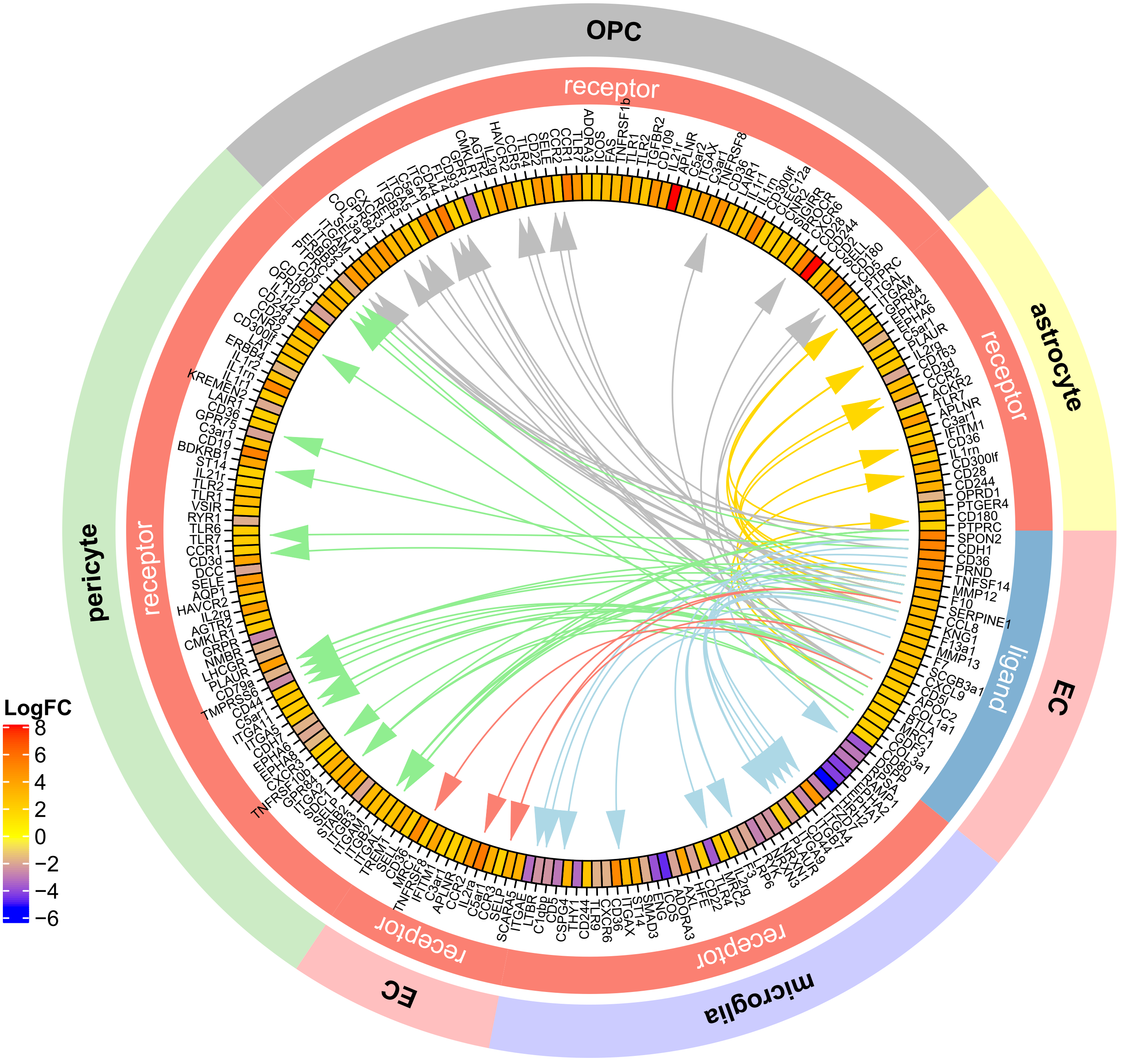

**d**

EC to other cells in mouse VaD (identified by mouse L-R pool)

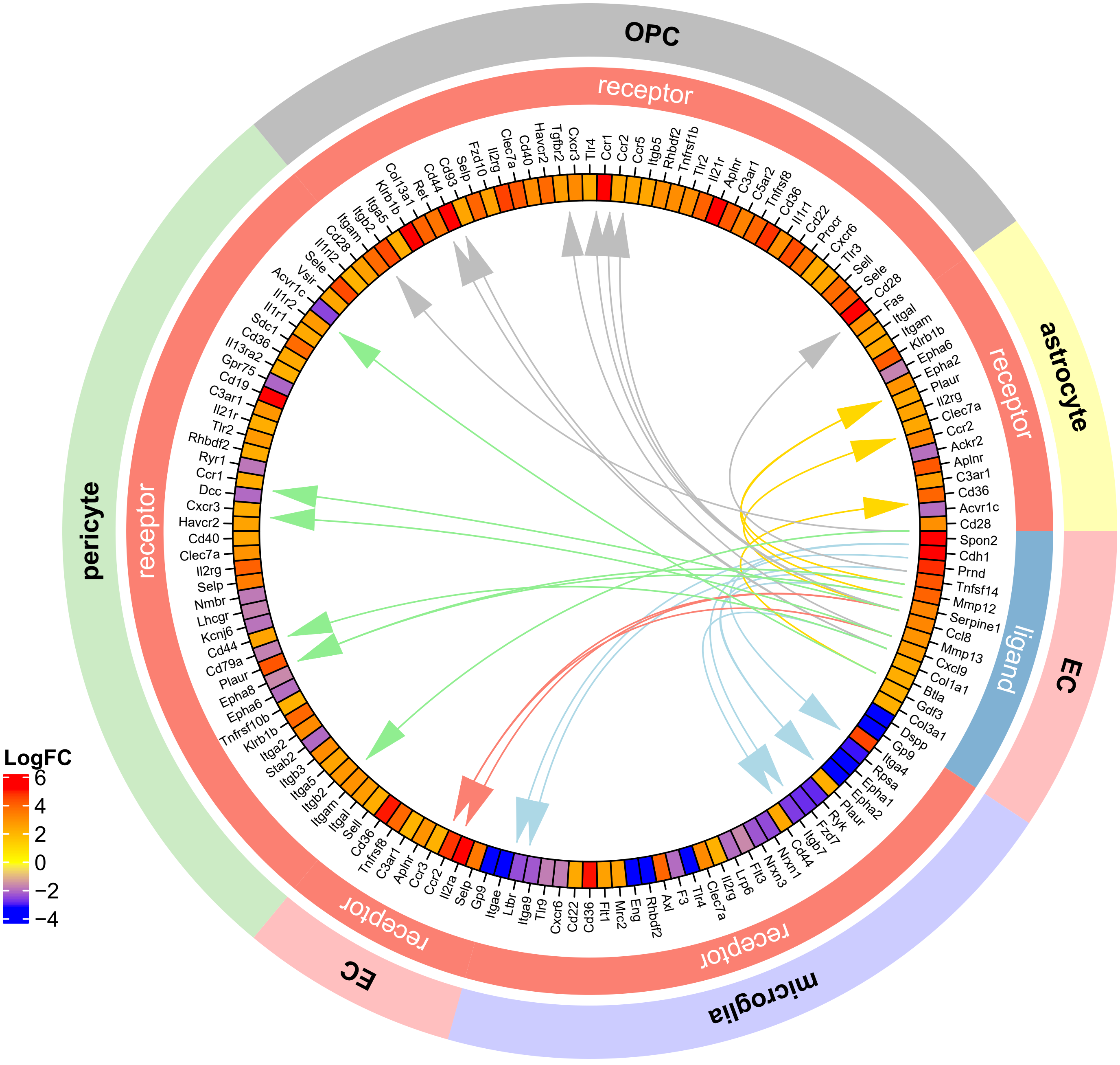

**e**

EC to other cells in human VaD (identified by human L-R pool)

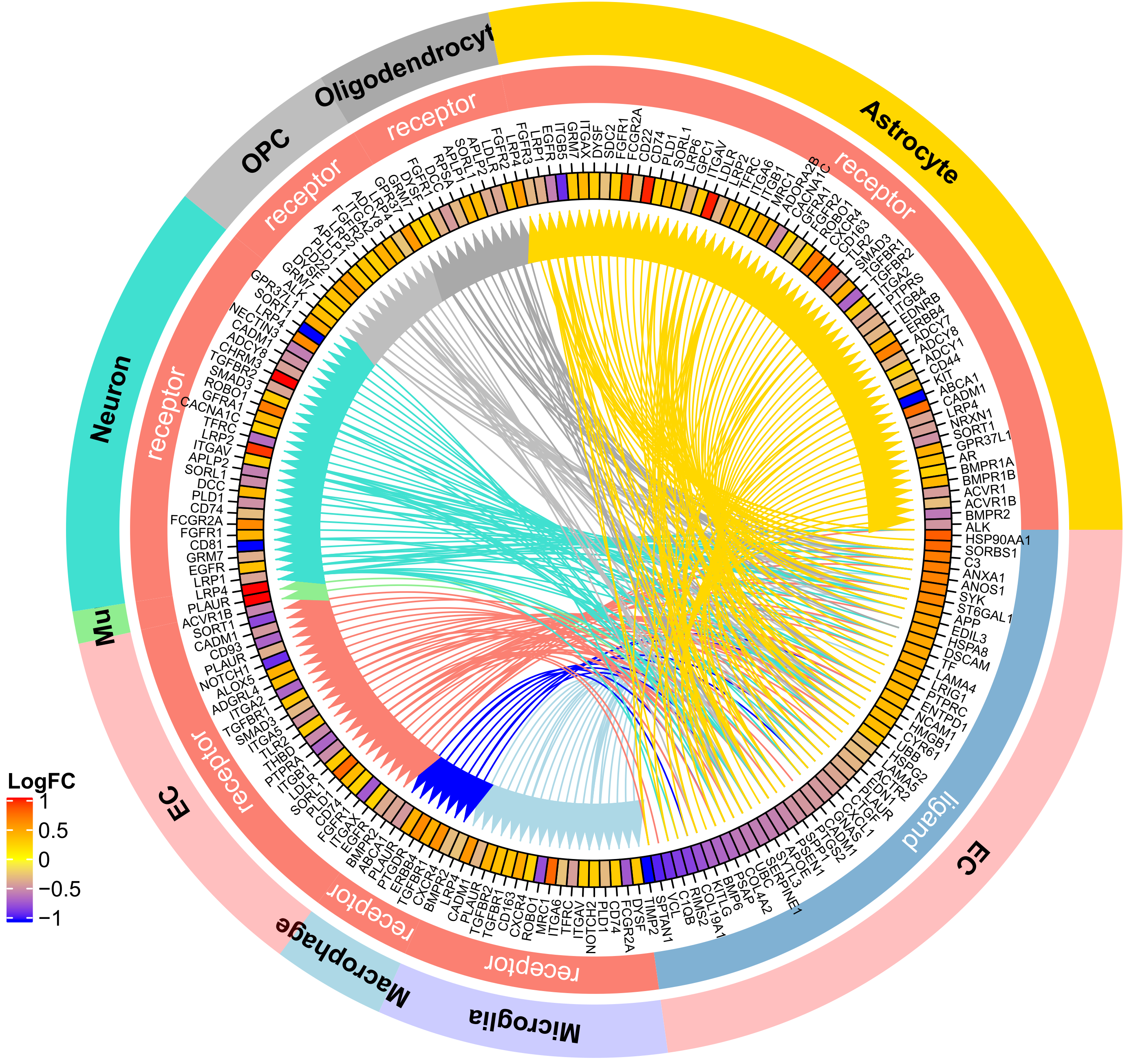

**f**

EC to other cells in human VaD (identified by mouse L-R pool)

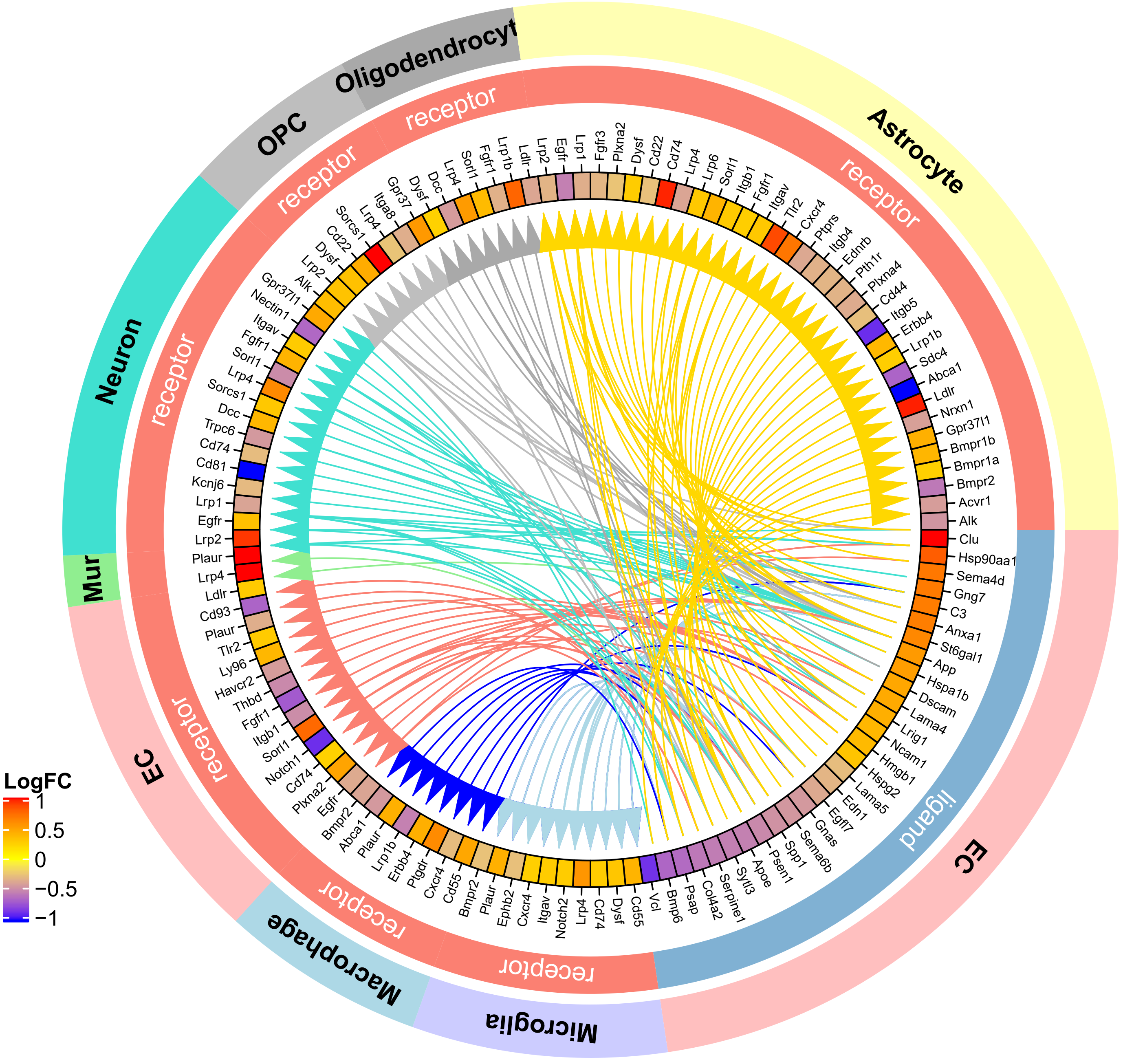

**g**

Pericyte to other cells in mouse VaD (identified by human L-R pool)

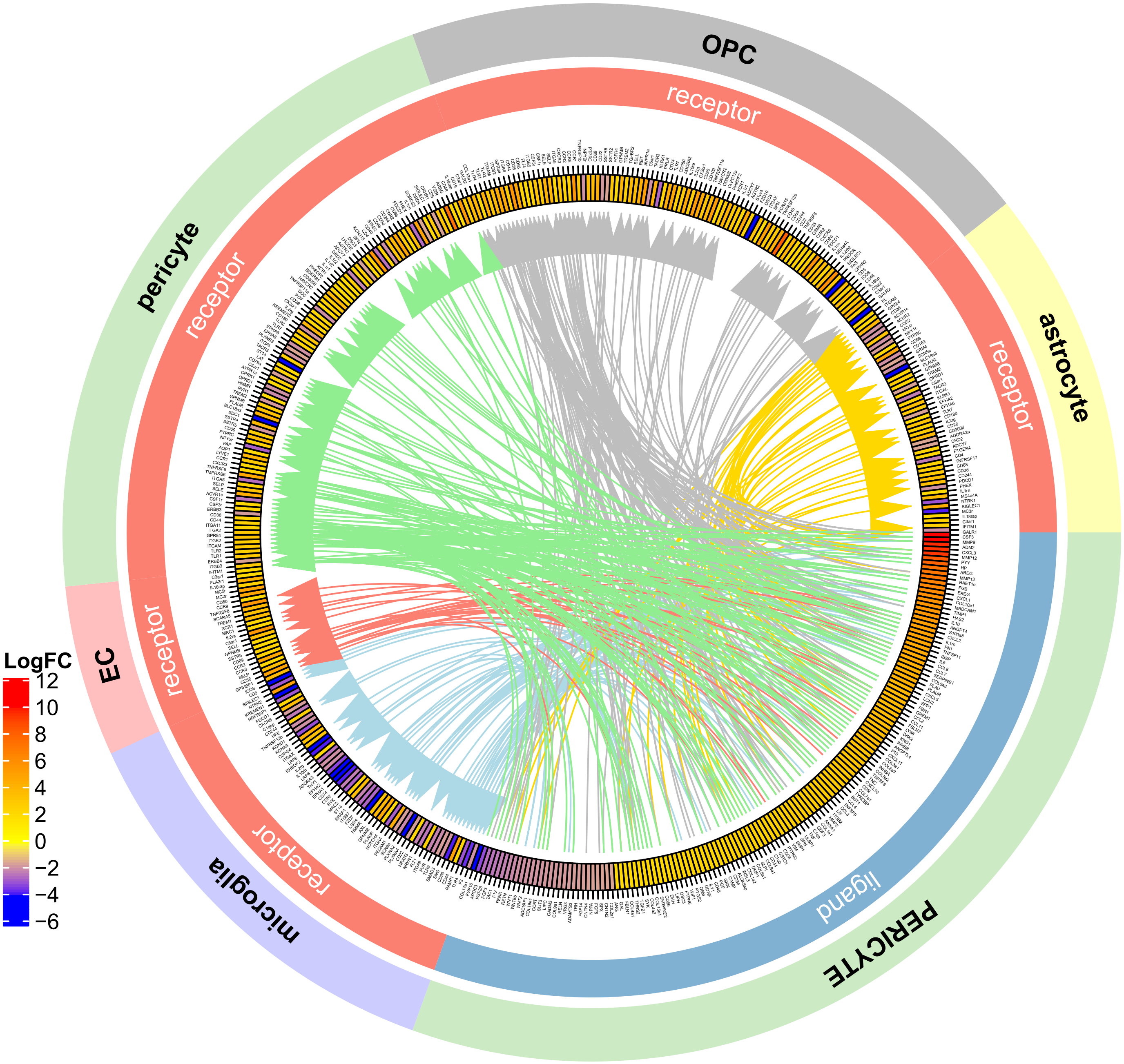

**h**

Pericyte to other cells in mouse VaD (identified by mouse L-R pool)

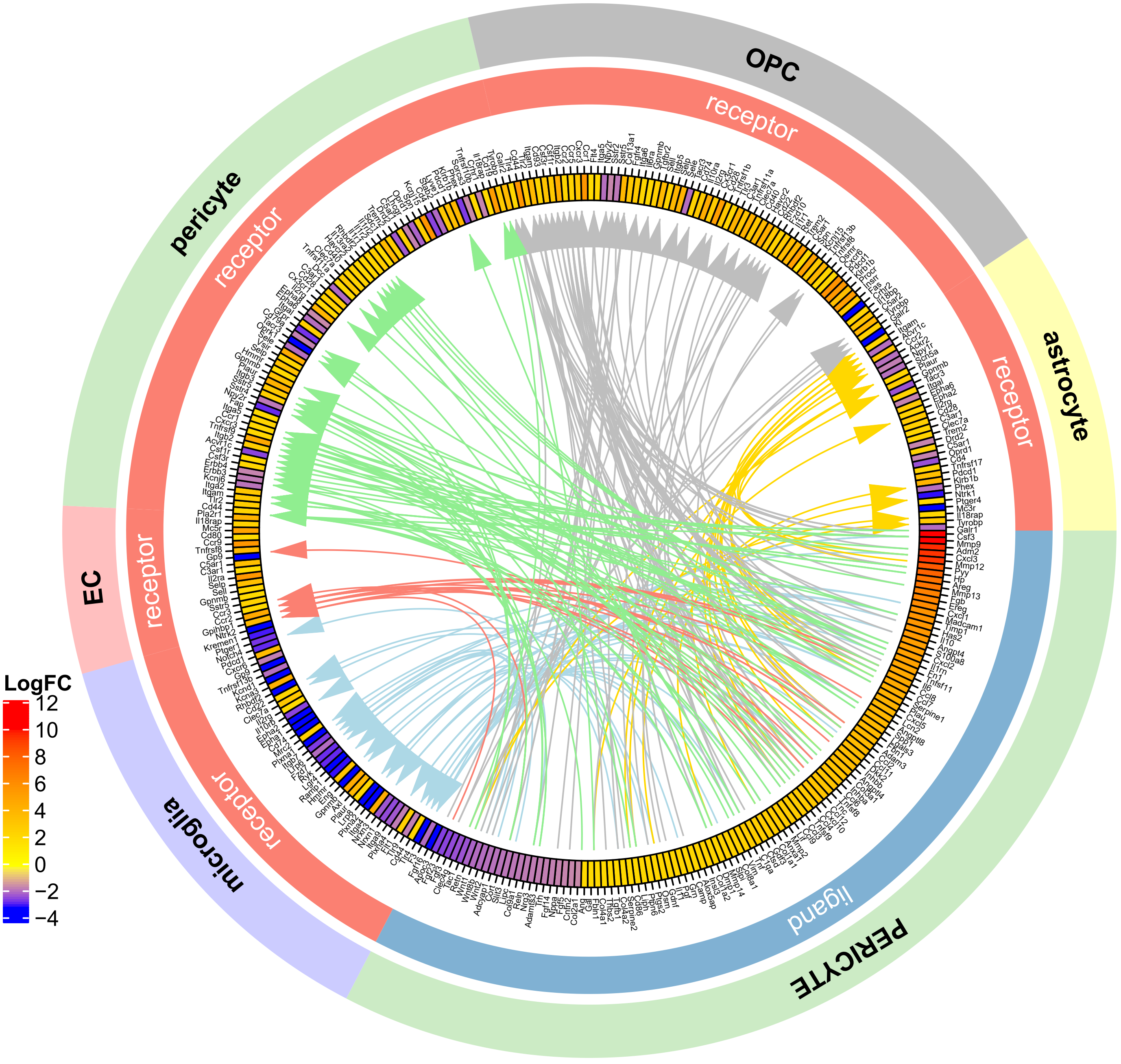

**i**

Mural cell to other cells in human VaD (identified by human L-R pool)

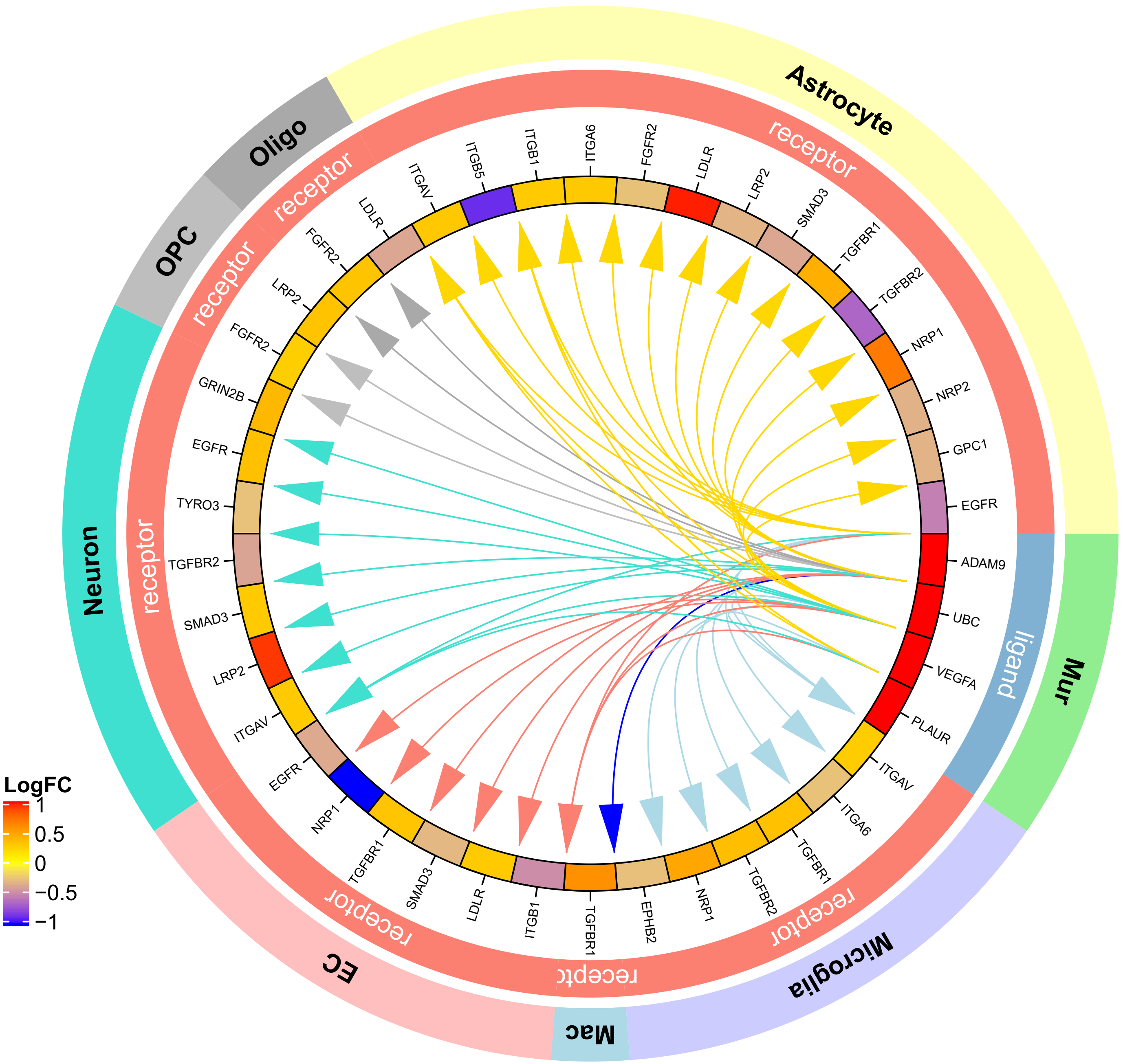

**j**

Mural cell to other cells in human VaD (identified by mouse L-R pool)

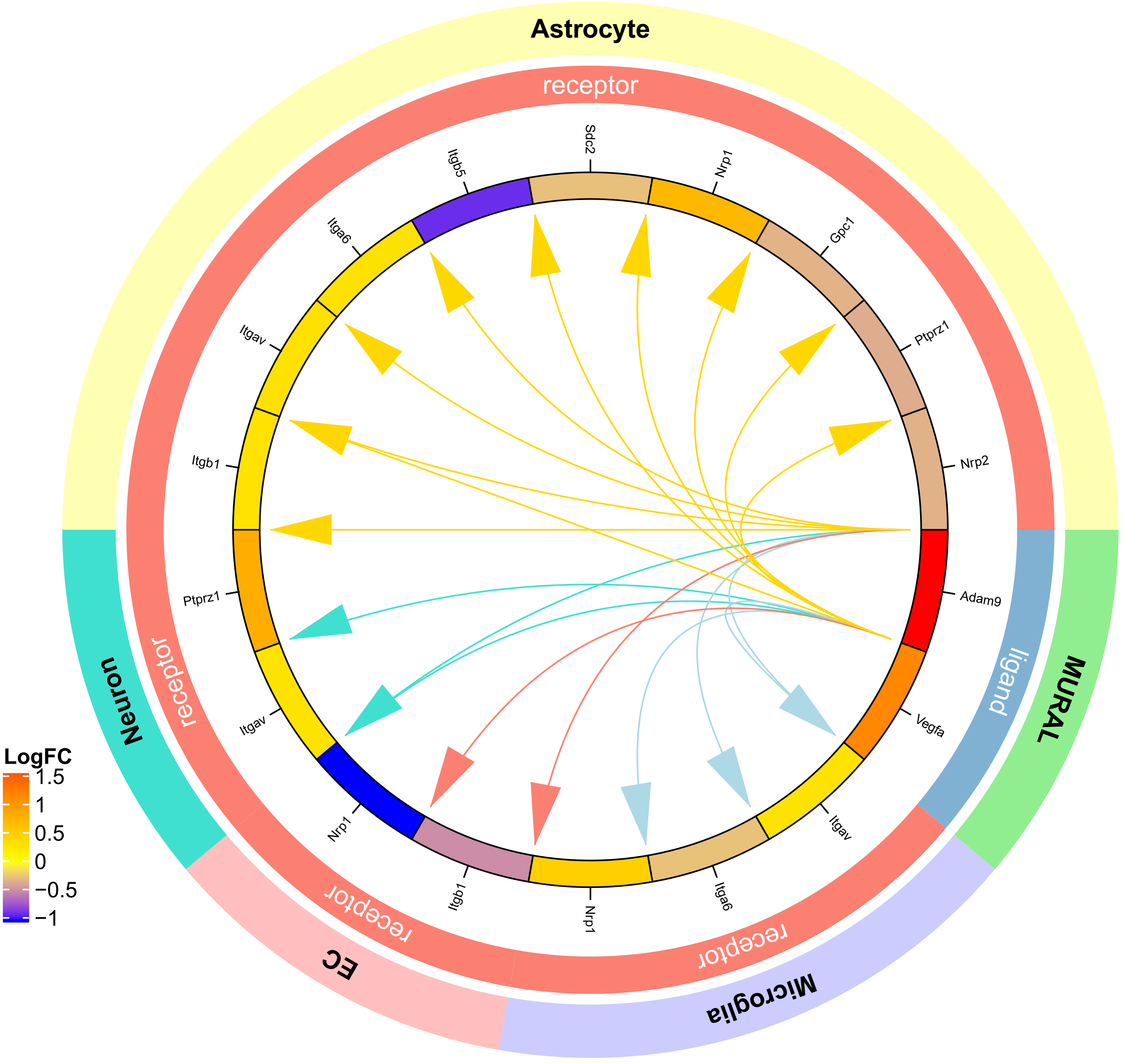

**k**

OPC to other cells in mouse VaD (identified by human L-R pool)

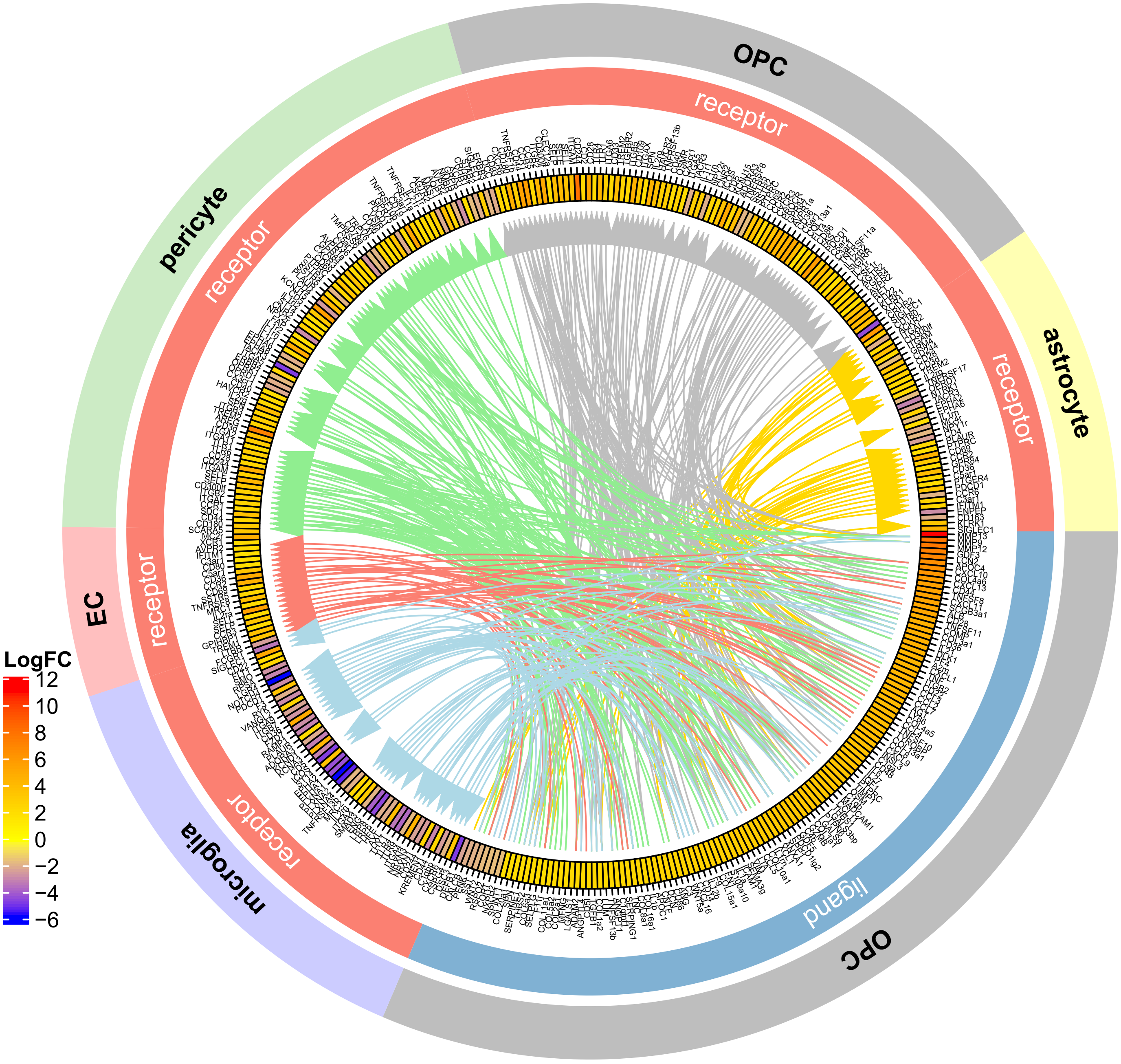

**I**

OPC to other cells in mouse VaD (identified by mouse L-R pool)

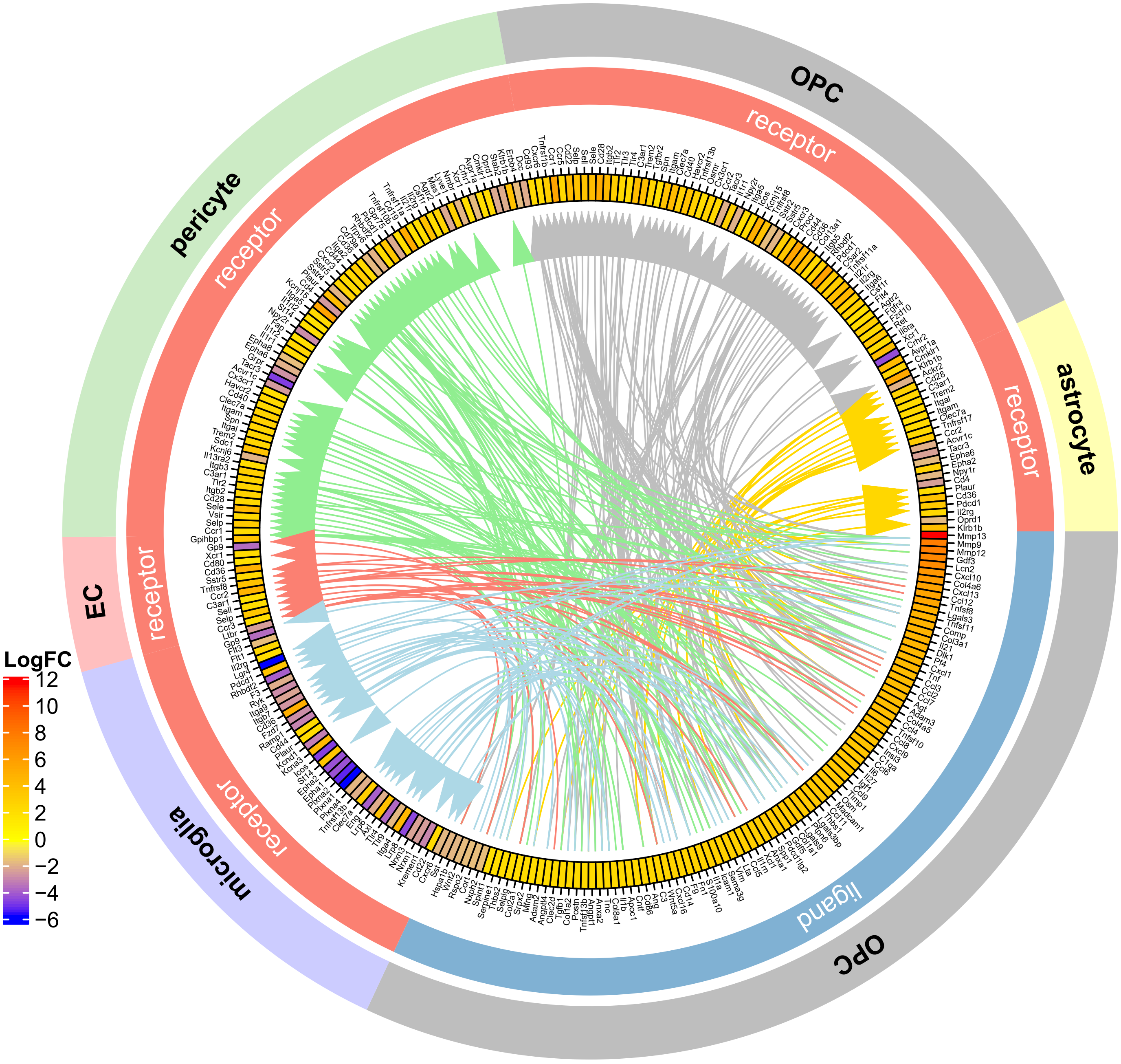

**m**

OPC to other cells in human VaD (identified by human L-R pool)

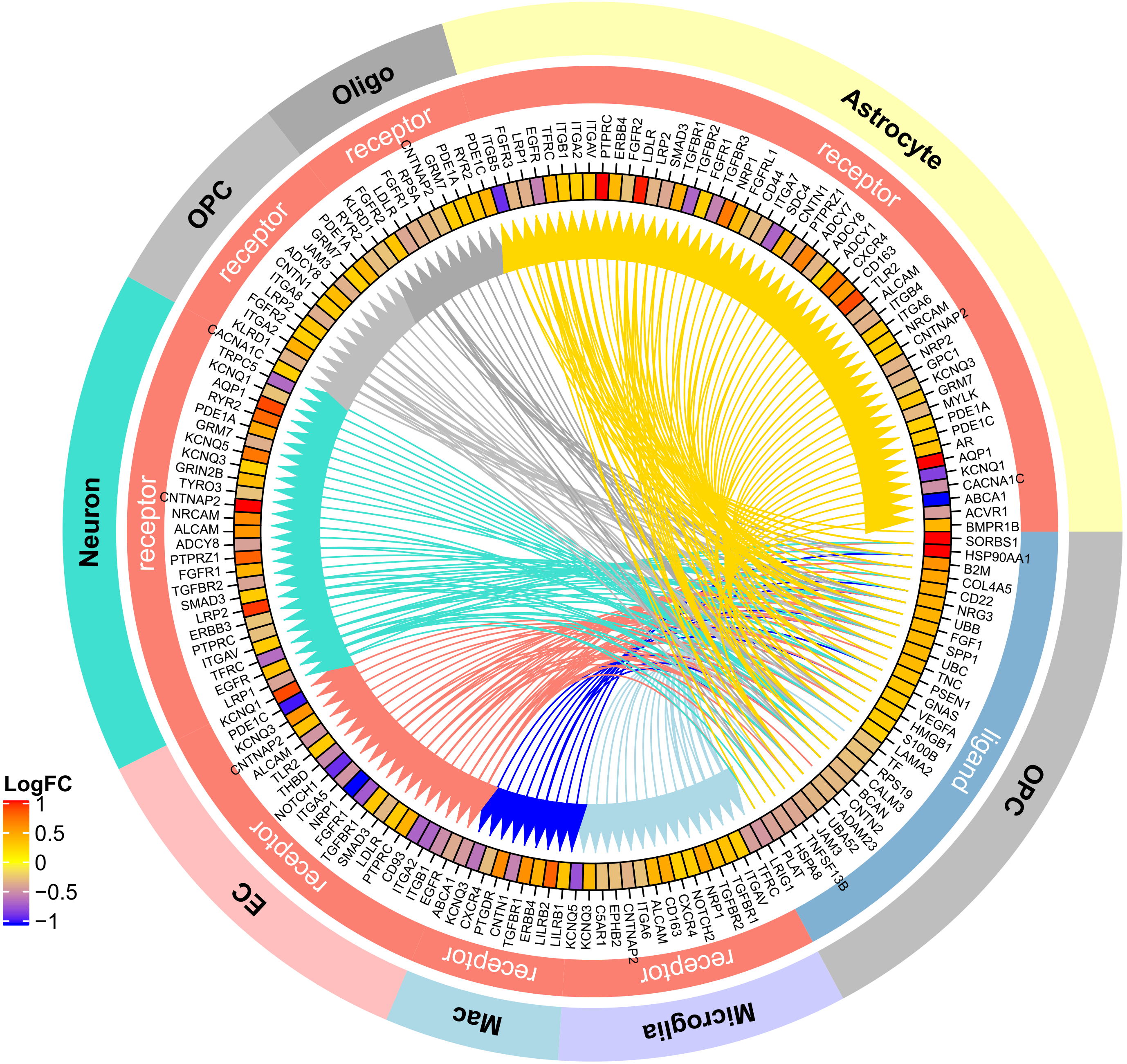

**n**

OPC to other cells in human VaD (identified by mouse L-R pool)

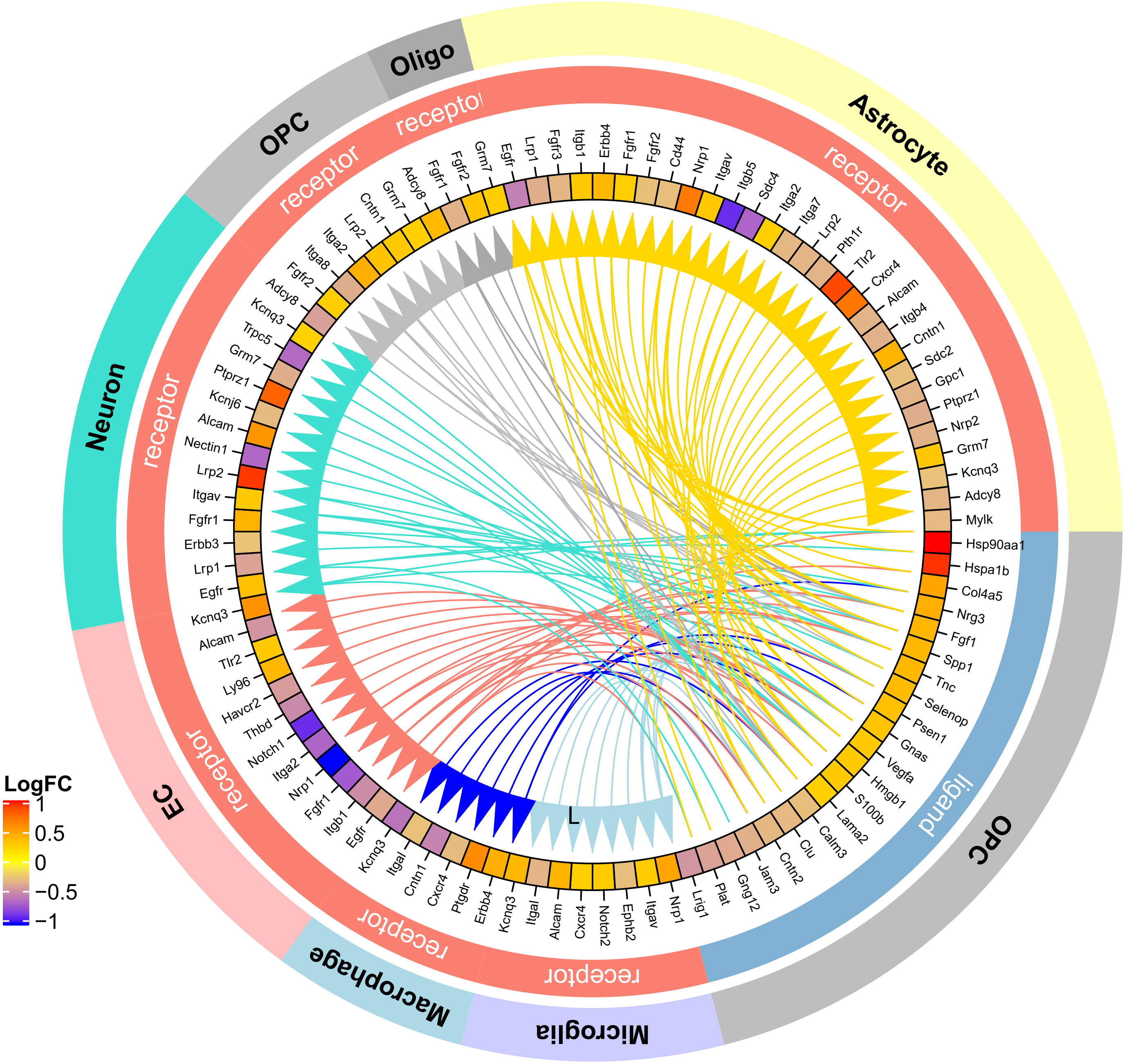

**o**

Astrocyte to other cells in mouse VaD (identified by human L-R pool)

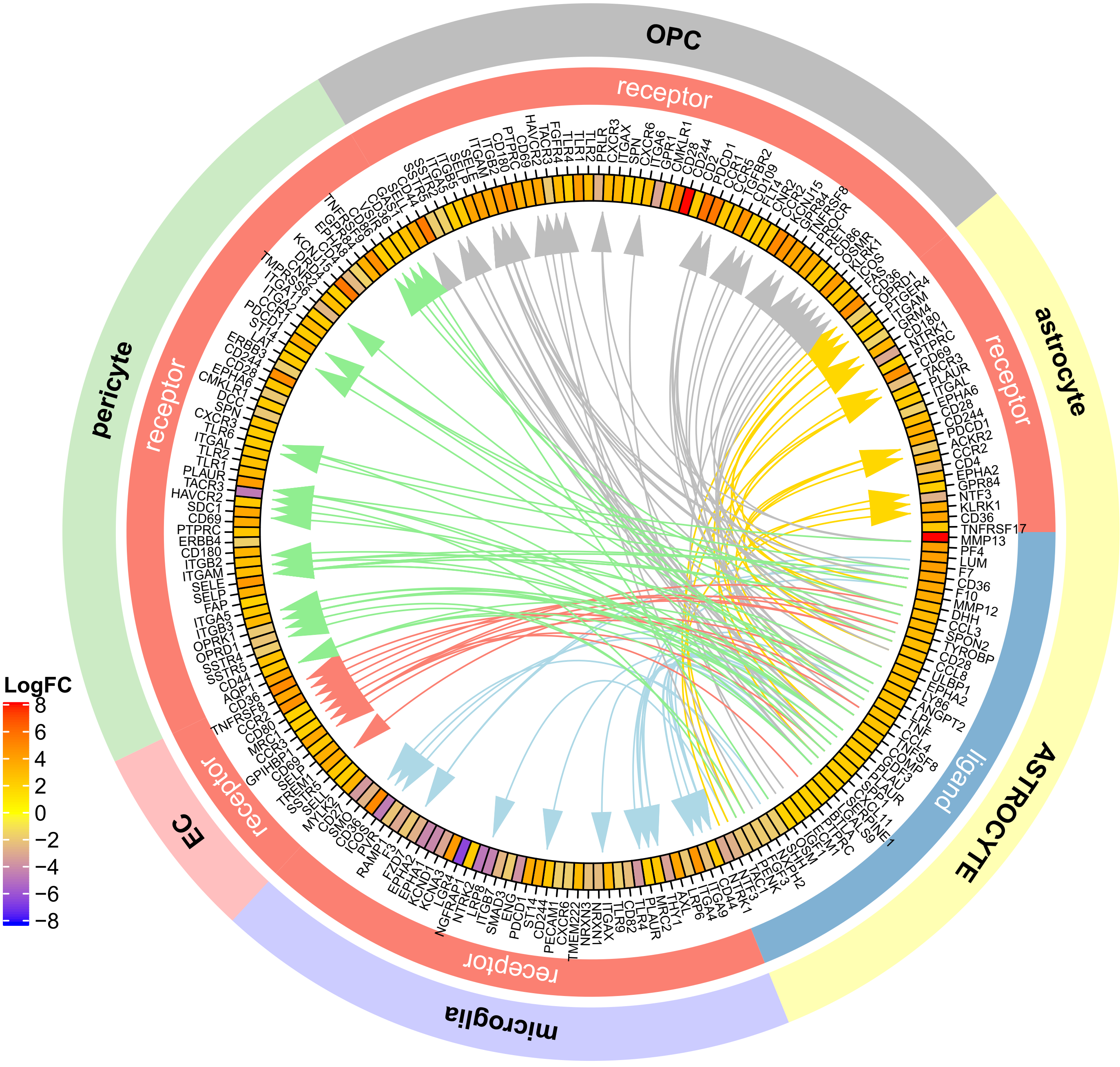

**p**

Astrocyte to other cells in mouse VaD (identified by mouse L-R pool)

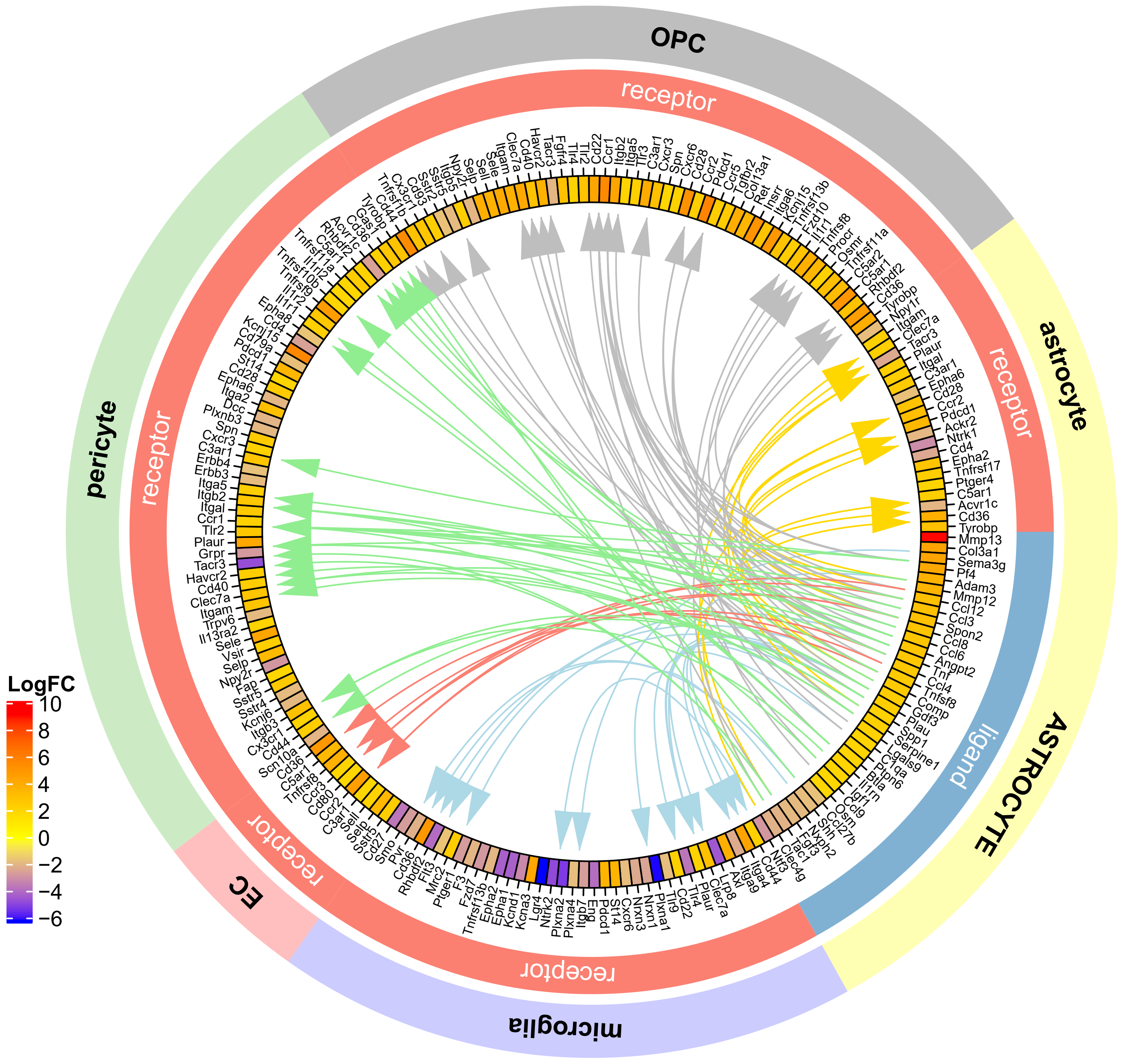

**q**

Astrocyte to other cells in human VaD (identified by human L-R pool)

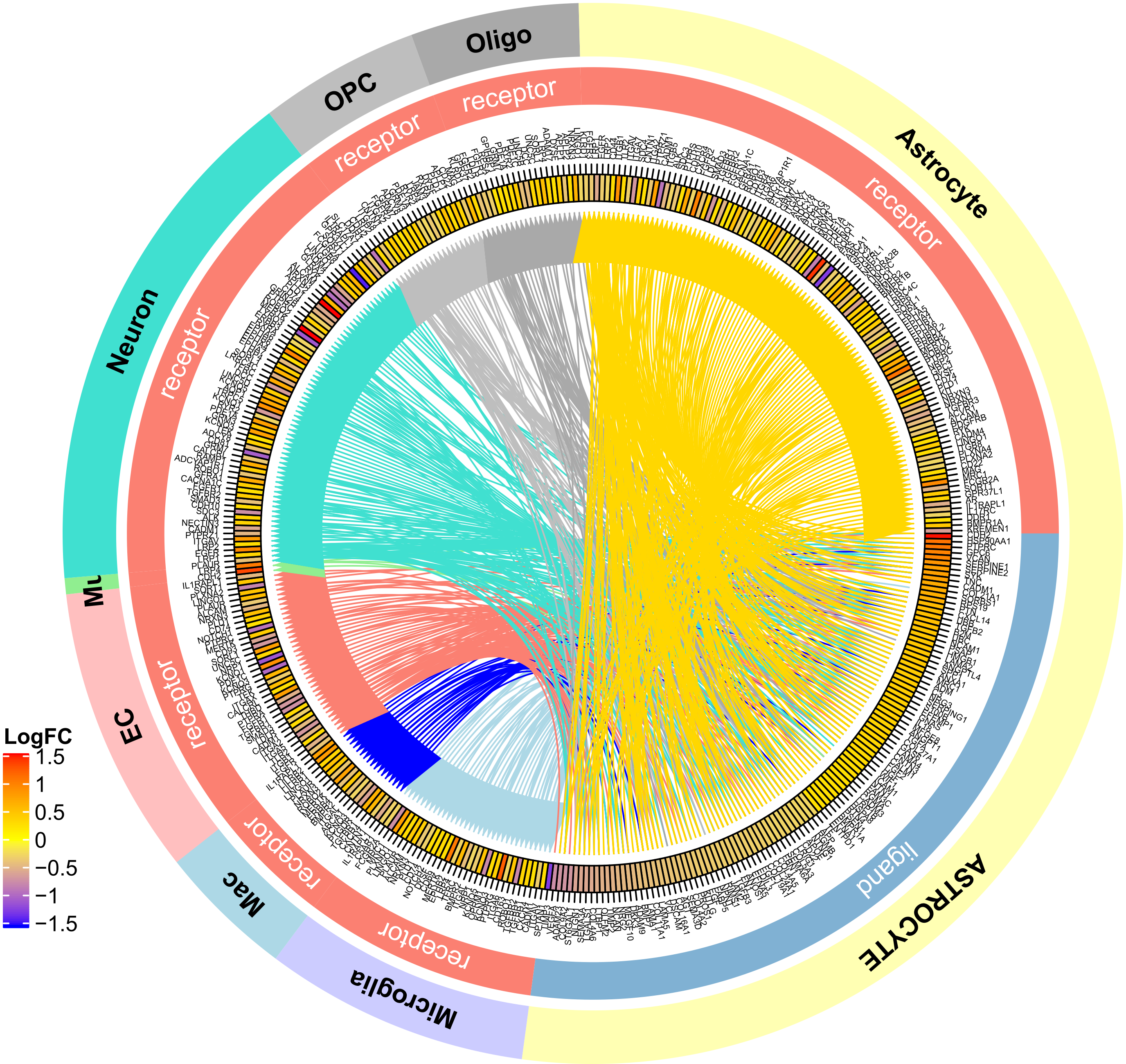

**r**

Astrocyte to other cells in human VaD (identified by mouse L-R pool)

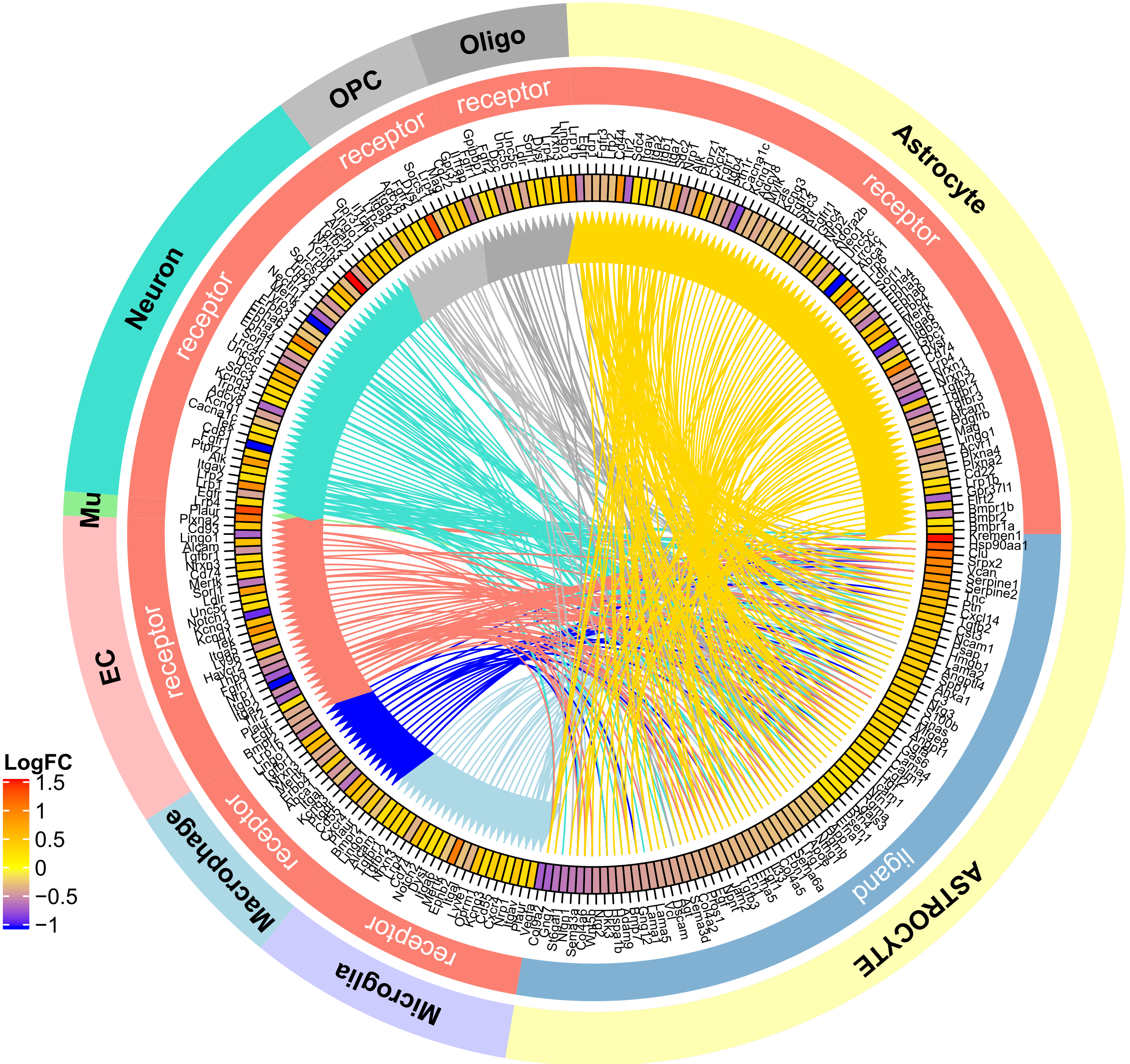
