## Supplementary Figure 9 for "Intercellular Signaling Pathways as Therapeutic Targets for Vascular Dementia Repair"

**a** **Tie2-cre x PHP-CAG-Flex-Rpl22-HA, TRAP (Input and Pulldown in Con and VaD)**

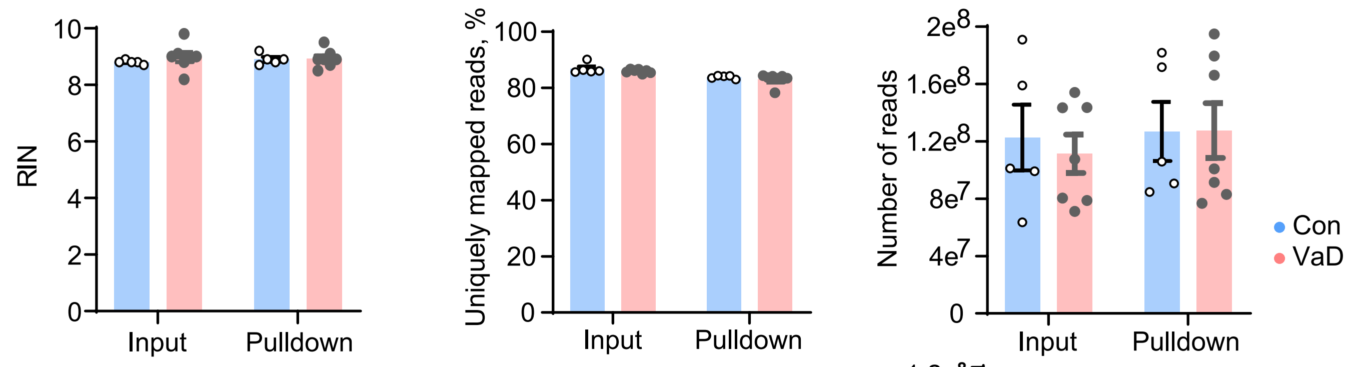


**b** **Tbx18-creER::Rpl22-HA, TRAP (Input and Pulldown in Con and VaD)**

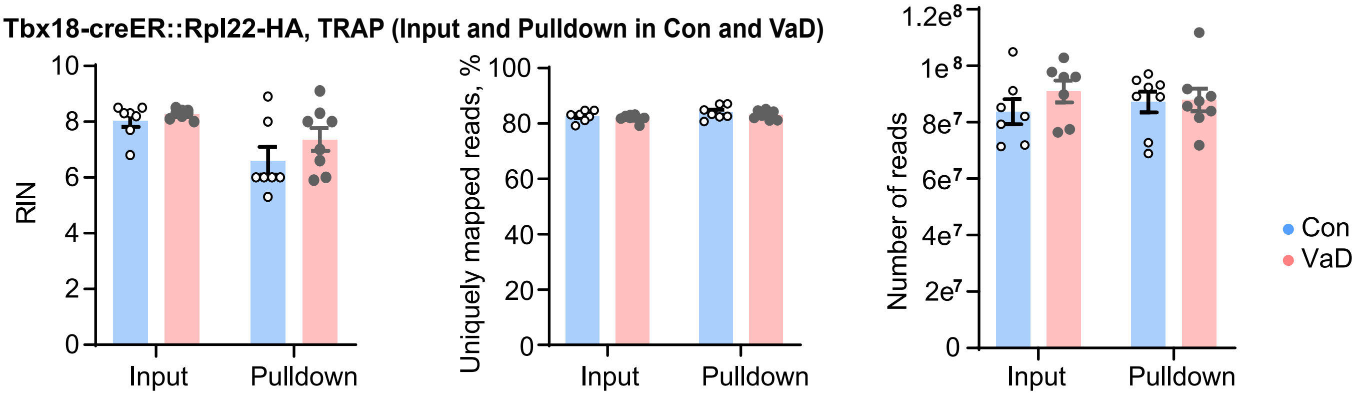


**c** **Ng2-creER::Rpl22-HA, TRAP (Input and Pulldown in Con and VaD)**

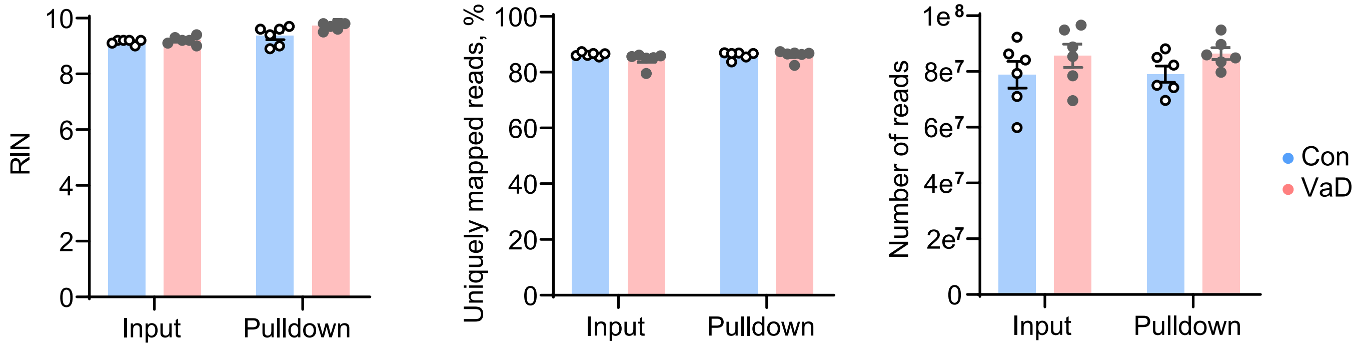


**d** **C57 x Lenti-GfaABC1D-Rpl22-HA, TRAP (Input and Pulldown in Con and VaD)**

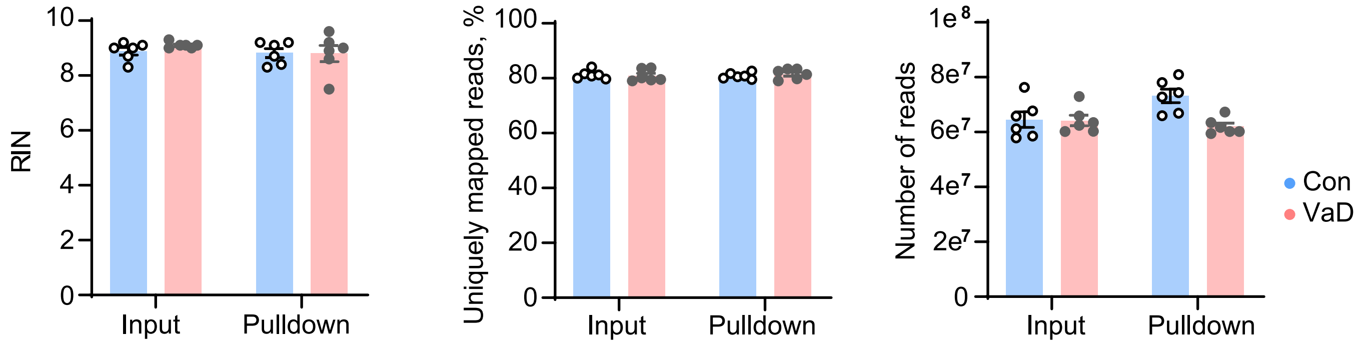
